# Exo1-protected DNA nicks direct crossover formation in meiosis

**DOI:** 10.1101/2021.08.29.458102

**Authors:** Michael Gioia, Lisette Payero, Gianno Pannafino, Jun Jie Chen, Sagar Salim, Ghanim Fajith, Amamah F. Farnaz, Sherikat Momoh, Michelle Scotland, Vandana Raghavan, Carol Manhart, Akira Shinohara, K.T. Nishant, Eric Alani

## Abstract

In most sexually reproducing organisms crossing over between chromosome homologs during meiosis is critical for the viability of haploid gametes. Most crossovers that form in meiosis in budding yeast result from the biased resolution of double Holliday Junction (dHJ) intermediates. This dHJ resolution step involves the actions Rad2/XPG family nuclease Exo1 and the Mlh1-Mlh3 mismatch repair endonuclease. At present little is known about how these factors act in meiosis at the molecular level. Here we show that Exo1 promotes meiotic crossing over by protecting DNA nicks from ligation. We found that structural elements in Exo1 required for interactions with DNA, such as bending of DNA during nick/flap recognition, are critical for its role in crossing over. Consistent with these observations, meiotic expression of the Rad2/XPG family member Rad27 partially rescued the crossover defect in *exo1* null mutants, and meiotic overexpression of Cdc9 ligase specifically reduced the crossover levels of *exo1* DNA binding mutants to levels approaching the *exo1* null. In addition, our work identified a role for Exo1 in crossover interference that appears independent of its resection activity. Together, these studies provide experimental evidence for Exo1-protected nicks being critical for the formation of meiotic crossovers and their distribution.

## INTRODUCTION

Cells in meiosis undergo a single round of DNA replication followed by reductional and equational chromosomal divisions to produce haploid gametes. In most eukaryotes, including budding yeast and humans, the accurate segregation of homologous chromosomes during the first reductional division (Meiosis I) requires the formation of crossovers between homologs. Physical linkages created by crossovers and sister chromosome cohesions distal to the crossover site are critical for proper segregation of chromosome pairs during Meiosis I (Maguire, 1974; Hunter, 2015; Zickler and Kleckner, 2015). The inability to establish these physical connections can lead to improper chromosome segregation and aneuploidy, and in humans is thought to be an important cause of birth defects and miscarriages (Hassold and Hunt, 2001; Nagaoka et al., 2012; Hunter, 2015).

In baker’s yeast crossover formation in meiotic prophase is initiated through the genome-wide formation of roughly 150 to 200 Spo11-induced double-strand breaks (DSBs; Keeney et al., 1997; Pan et al., 2011). These breaks are resected in a 5’ to 3’ direction to form 3’ single-stranded tails (Cao et al., 1990; Padmore et al., 1991). Strand exchange proteins coat the single stranded tails and promote their invasion into homologous sequences in the unbroken homolog (Hunter, 2015). In the major crossover pathway (Class I), the resulting D-loop intermediate is stabilized by ZMM proteins including Zip2-Zip4-Spo16 and Msh4-Msh5 to form a single end invasion intermediate (SEI; Figure 1A; Hunter and Kleckner, 2001; Fung et al., 2004; Borner et al., 2004; Lynn et al., 2007; De Muyt et al., 2018). This recombination intermediate forms concomitantly with the synaptonemal complex, a structure that is thought to remove chromosomal tangles and interlocks during the homology search process (Padmore et al., 1991; Sym et al., 1993; de Boer and Heyting, 2006). DNA synthesis from the SEI, followed by second-end capture, results in the formation of the double-Holliday junction intermediate (dHJ). The dHJ is thought to be stabilized by Msh4-Msh5 and resolved in a biased orientation to form ∼90 crossovers (COs) in the yeast genome that are distributed so that they are evenly spaced (crossover interference) and every homolog pair receives at least one crossover (Figure 1A; Szostak et al., 1983; Sym and Roeder, 1994; Schwacha and Kleckner, 1994, 1995; Borner et al., 2004; Hillers, 2004; Jones and Franklin, 2006; Mancera et al., 2008; Zakharyevich et al., 2012).

**Figure 1.**
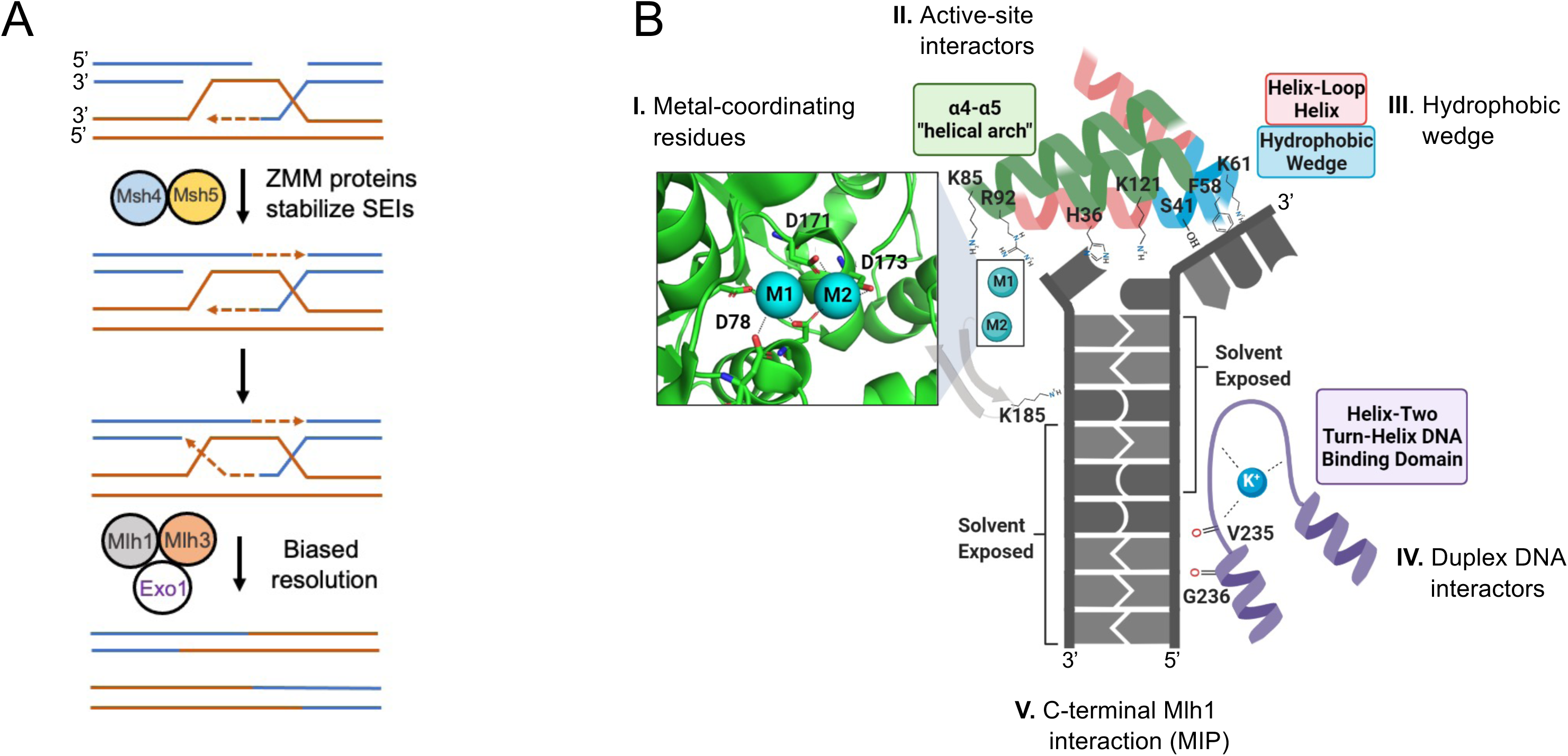
Metal binding, active site interactions, and DNA contact sites of Human Exo1 based on the crystal structure of the Exo1-5’ recessed DNA complex. A. Canonical model showing roles for Msh4-Msh5, Mlh1-Mlh3, and Exo1 in meiotic crossover resolution. See text for details. B. Close-up of the Exo1 active site (adapted from Orans et al. (2011) using crystal structure PDB #3QEA). We highlight the following residues which were mutated in this study (Figure S1): Group I; acidic residues (D78, D171, D173) which coordinate the two metal ions. Group II; residues that are part of the α4-α5 helical arch involved in fraying (H36, K85, K121) and coordinating the scissile bond adjacent to the catalytic metals that interact with the active site (R92). Group III; S41, F58, K61, which are part of a hydrophobic wedge which induces the sharp bend in DNA at the site of a nick. Group IV; K185, G236, residues that interact with duplex DNA (K185, G236). Group V; residues (F447 ,F448) in a region of Exo1 that interact with Mlh1. The *exo1-F447A,F448A* allele is abbreviated in the text as *exo1-MIP*.

How dHJs are resolved in a biased manner to form crossovers is a major unanswered question. Investigators have suggested that the presence of nicks in dHJs ensures biased resolution by creating asymmetric structures that are resolved to form crossover-only products (reviewed in Machin et al., 2020). In support of such ideas, whole genome sequencing of hDNA tracts formed in meiosis inferred a model in which meiotic crossover resolution is biased towards DNA synthesis tracts (Martini et al., 2011; Marsolier-Kergoat et al., 2018). In this model nicks maintained at the ends of synthesis tracts could direct biased and asymmetric cleavage of the dHJ by recruiting a nick-binding protein that acts in the resolution mechanism. However, such a model is inconsistent with a denaturing gel analysis of dHJs that form at a meiotic hotspot in *S. cerevisiae*; this work showed that all single strands within the dHJs are continuous (Schwacha et al., 1994; 1995). It is also inconsistent with recent work in *S. cerevisiae* showing that a vast majority of crossovers initiated at another hotspot displayed evidence of branch migration, with about half of the COs having formed from dHJs located on one side of the initiating double-strand break. In such a situation, nicks should not be present in positions that direct biased resolution (Ahuja et al., 2021). Thus, it remains unclear if nicks participate in meiotic crossover formation.

What factors act in the biased resolution of dHJs? The MMR endonuclease Mlh1-Mlh3 and the XPG/Rad2 family nuclease Exo1 have been shown to act in meiotic crossover resolution, with *mlh3Δ* and *exo1Δ* single and double mutant strains displaying similar crossover defects in crossing over (Khazanehdari and Borts, 2000; Zakharyevich et al., 2010; 2012). Biochemical analyses of Mlh1-Mlh3 indicate that its endonuclease activity is required for its role in crossover formation, but not as a structure-specific nuclease that symmetrically cleaves Holliday junctions (Nishant et al., 2008; Rogacheva et al., 2014; Ranjha et al., 2014; Manhart et al., 2017). Exo1 acts in many steps in DNA metabolism, including creating 3’ single-stranded ends for homologous recombination, telomere maintenance, DNA mismatch repair, DNA replication, and crossover-specific dHJ resolution in meiosis. Exo1 contains an N-terminal Rad2/XPG nuclease domain that is conserved in Rad2/XPG family members and an unstructured C-terminal tail that interacts with the mismatch repair factors Msh2 and Mlh1 (Tishkoff et al., 1997; Tran et al., 2001). *In vitro* studies demonstrated that Exo1 displays a robust and processive 5’ to 3’ exonuclease on the ends of a double-strand break, and on gapped and nicked duplex DNA. In addition, it displays 5’ flap endonuclease activity (Kunkel and Erie, 2015; Goellner et al., 2015; Szankasi and Smith, 1992; Fiorentini et al., 1997; Lee and Wilson, 1999; Tran et al., 2002; Genschel and Modrich, 2003; Zakharyevich et al., 2010).

In meiosis *exo1Δ* strains display a defect in the 5’ to 3’ resection of Spo11-induced DSBs and a meiotic crossover defect. In fact, resection is reduced in *exo1Δ* to an average of 270 nt compared to 800 nt in *wild-type*. Despite showing these defects, *exo1Δ* mutants display *wild-type* timing and levels of meiotic recombination intermediates, including dHJs (Zakharyevich et al., 2010). Genetic analysis showed that disruption of a conserved Mlh1-Interaction Protein sequence (MIP box) in the Exo1 C-terminal domain conferred intermediate defects in meiotic crossing over, suggesting that Exo1 promotes meiotic crossovers through interactions with Mlh1 and possibly other factors (Amin et al., 2001; Argueso et al., 2003; Tran et al., 2004; 2007; Zakharyevich et al., 2010). Curiously, an *exo1* mutation (*D173A*) that disrupts a metal binding site critical for nuclease function was shown to have only a minimal impact on meiotic crossing over. Together these analyses suggested that Exo1’s interactions with Mlh1-Mlh3, but not its nuclease function, are critical for crossover formation (Abdullah et al., 2004; Zakharyevich et al., 2010; Keelagher et al., 2011).

The studies outlined above in addition to recent biochemical analyses have led to the proposal that Mlh1-Mlh3 interacts with Exo1, Msh4-Msh5 and the DNA polymerase processivity factor PCNA for biased resolution of double Holliday junctions (Cannavo et al., 2020; Sanchez et al., 2020; Kulkarni et al., 2020). This proposal suggests that DNA signals are present in dHJ intermediates that are critical for such resolution; however, these studies have not provided direct evidence for such signals. Here we provide genetic evidence that Exo1 acts to protect DNA from being ligated in recombination intermediates during the formation of crossover products. We also show that it plays a critical role in ensuring that meiotic crossovers are widely spaced for proper chromosome segregation in the Meiosis I division. These observations provide evidence for dynamic and distinct roles for Exo1 in both crossover placement and for maintaining a nicked recombination intermediate for the resolution of dHJs into crossovers.

## RESULTS

### Mutations in metal coordinating and active site residues in Exo1 do not disrupt meiotic crossing over

The crystal structure of human Exo1 with 5’ recessed DNA (PDB #3QE9) identified two metals in the catalytic site of the Exo1-DNA structure, with residue D171 assisting D173 in coordinating one metal, and residue D78 coordinating the other, to hydrolyze the phosphodiester backbone of DNA (Figures 1B and S1; Orans et al., 2011; Mueser et al.,1996; Hwang et al., 1998; Feng et al., 2004; Shi et al., 2017). While the *exo1-D173A* mutation in baker’s yeast was shown to disrupt Exo1 nuclease activity (Tran et al., 2002), mutation of other amino acids that coordinate the catalytic metals was not performed. Mutation of other nucleases that act through a two-metal catalysis mechanism suggested that altering a single metal binding residue does not fully ablate function and could create novel functions, perhaps because a water molecule can substitute as a ligand (Schiltz et al., 2019). For example, work by Lee et al. (2002) showed that the human exo1-D78A and exo1-D173A mutant proteins display nuclease activities, though at levels significantly lower than the wild-type protein.

In baker’s yeast meiosis, mutation of a single metal binding residue (*exo1-D173A*) caused a disruption in the 5’ to 3’ resection steps of meiotically induced DSBs, but only minor, if any defects in meiotic crossing over, suggesting that Exo1’s nuclease functions were not required in this step (Abdullah et al., 2004; Zakharyevich et al., 2010). We purified exo1-D173A from baculovirus infected *Sf9* cells (Materials and Methods), but were unable to purify a full length variant (exo1-D78A,D173A) expected to disrupt both metal binding sites (Figure S2). We tested the nuclease activity of exo1-D173A on a 2.7 kb pUC18 substrate containing four pre-existing nicks (Figure S2A) as well as supercoiled plasmids. As shown in Figure S2A and C, exo1-D173A was deficient for exonuclease activity on the substrate containing four pre-existing nicks. However, exo1-D173A displayed a weak DNA nicking activity on closed circular DNA similar to that seen for Mlh1-Mlh3 (∼10% nicking of pUC18 at 20 nM exo1-D173A compared to ∼20% nicking at 20 nM Mlh1-Mlh3), suggesting that a role for Exo1 nuclease activity in crossover resolution was not fully resolved (Manhart et al., 2017). In contrast, wild-type Exo1 did not display such nicking activity, consistent with previous work showing that human Exo1 displayed little or no endonuclease activity on blocked-end DNA substrates (Figure S2B; Lee et al., 2002). Interestingly, the addition of a mutation predicted to be critical for DNA binding, G236D (see below), decreased the nicking activity of the exo1-D173A protein by about two-fold, consistent with previous studies indicating that Exo1 nuclease activity was dependent on its DNA binding activity (Figure S2D; Orans et al., 2011).

To test the effect of mutations in the Exo1 catalytic site we made D78A, D171A, and D173A mutations (Group I, Figure 1B) in combination to disrupt coordination of both metals. We also mutated residues in Exo1 which interact with and position DNA in an orientation to be cleaved (Orans et al., 2011). These residues (H36, K85, R92, K121, Group II) contribute to the fraying of the duplex DNA bases away from its complement and reside within an α4-α5 helical arch microdomain that forms part of the Exo1 active site (Figures 1B, S1). This microdomain is important for catalysis and also defines substrate specificity throughout the flap endonuclease (FEN) superfamily and consequently Exo1 5’ flap binding (Ceska et al., 1996; Devos et al., 2007; Gloor et al., 2010; Orans et al., 2011). Within this region R92 has been shown to be a critical residue for Exo1 catalysis; it interacts with the scissile bond on the DNA to position it adjacent to the catalytic metal core, and the R92A mutation dramatically decreased nuclease activity of human Exo1 *in vitro* to similar levels of the D173A metal-coordinating mutation (Orans et al., 2011). K121 (R in human Exo1) is part of the α5 helix and coordinates passage of the DNA substrate through the active site.

We analyzed meiotic crossing over by tetrad analysis at four consecutive intervals on Chromosome XV (104.9 cM map distance in *wild-type*, 52 cM in *exo1Δ*) and at one interval *(CEN8-THR1)* on Chromosome VIII (∼39% single crossovers in *wild-type*, 20% in *exo1Δ*; Figures 2A and 3A; Thacker et al., 2011). These two chromosomal regions showed defects in crossing over similar to those seen previously (*exo1Δ,* ∼2-fold decreased; *mlh3Δ,* ∼2-fold; *msh5Δ,* ∼3-fold; *exo1Δ mus81Δ*, ∼12-fold) and confirmed the epistatic relationship between *exo1Δ* and *mlh3Δ* (Figure 2B; Argueso et al., 2004; Nishant et al., 2008; Zahkaryevich et al., 2012; Al-Sweel et al., 2017). As shown in Figures 2B and 3B and Tables S1 and S2, disruption of either one or both metal binding sites of Exo1 (Group I) had minor if any effects on meiotic crossing over. There was a small crossover (<10%) reduction in some of the catalytic mutants compared to *wild-type*; this reduction could result from defects in DNA binding that result from perturbation of the active site. In fact, the human exo1-D78A mutant protein showed defects in binding to DNA flap structures (Lee et al., 2002). In addition, the *exo1-H36E*, *exo1-K85A/E*, *exo1-R92A* and *exo1-K121A/E* mutations (Group II) had very modest, if any effect on meiotic crossing over compared to *wild-type*, suggesting that coordination of the scissile bond for catalysis within the active site is not critical for crossing over. The dramatic loss of nuclease activity seen with human Exo1 bearing K85A, R92A or K185A mutations (Orans et al., 2011; Li et al., 2019) further supports the dispensability of Exo1 catalytic activity for crossing over. These observations indicate that the critical function(s) of Exo1 in meiotic crossover resolution are not catalytic in nature.

**Figure 2.**
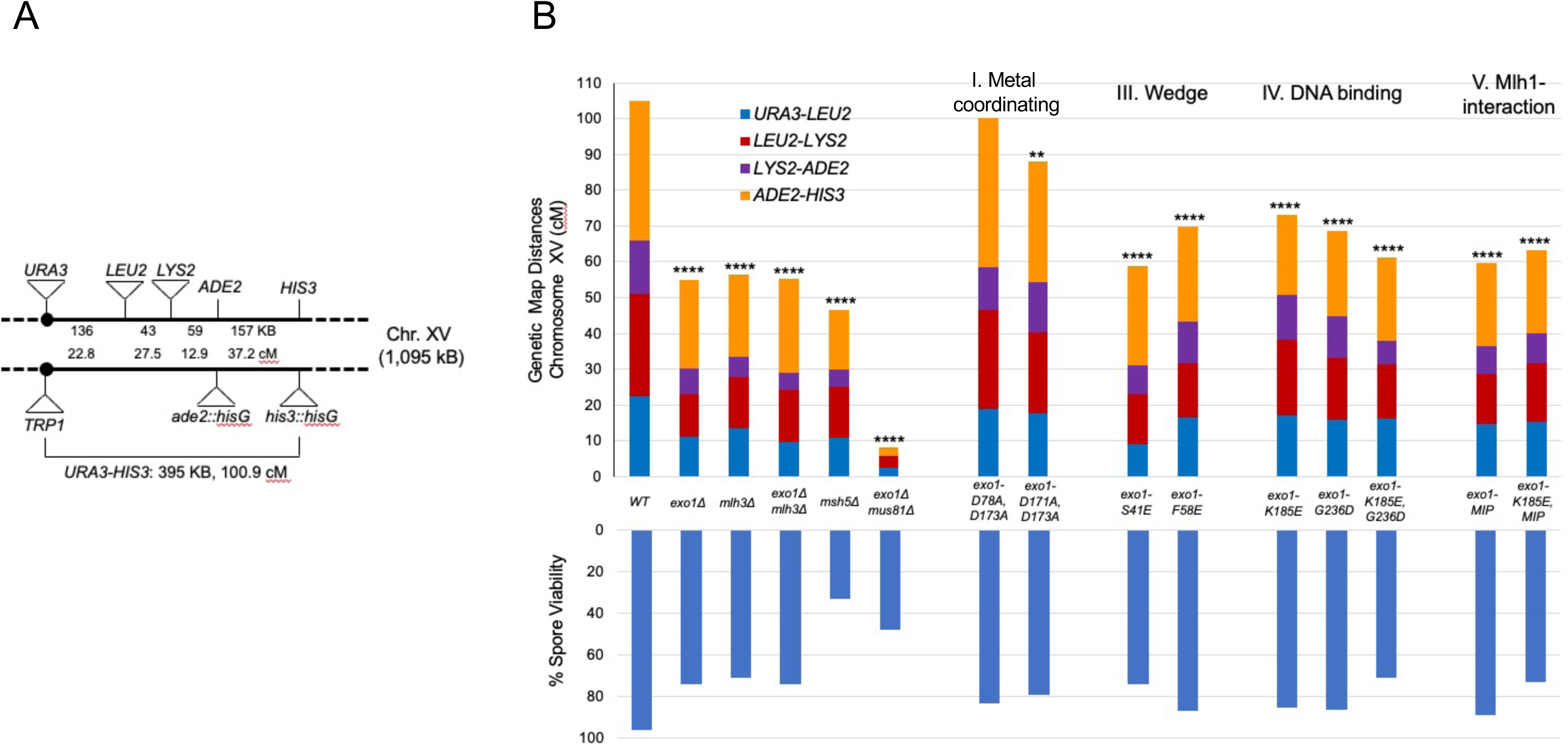
Meiotic crossover phenotypes in *exo1* mutant strains. A. Genetic markers on chromosome XV spanning the *CENXV-HIS3* interval in the EAY1108/1112 strain background (Argueso et al., 2004). The solid circle indicates the centromere. Distances between markers in KB and cM are shown for *wild-type* (not drawn to scale). B. Cumulative genetic distance (cM) in *wild-type* (*WT*) and *exo1* strains. Genetic map distances for the *URA3-HIS3* interval of chromosome XV in *wild-type* and the indicated mutant strains. Each bar is divided into sectors corresponding to genetic intervals in the *URA3-HIS3*, as measured from tetrads (T). The spore viability data obtained from tetrad analysis are shown, with the complete data set presented in Figure S3. The asterisks indicate the number of genetic intervals (0-4) that are distinguishable from *wild-type* in the indicated genotypes as measured using standard error calculated by Stahl Laboratory Online Tools (https://elizabethhousworth.com/StahlLabOnlineTools/; Table S2).

**Figure 3.**
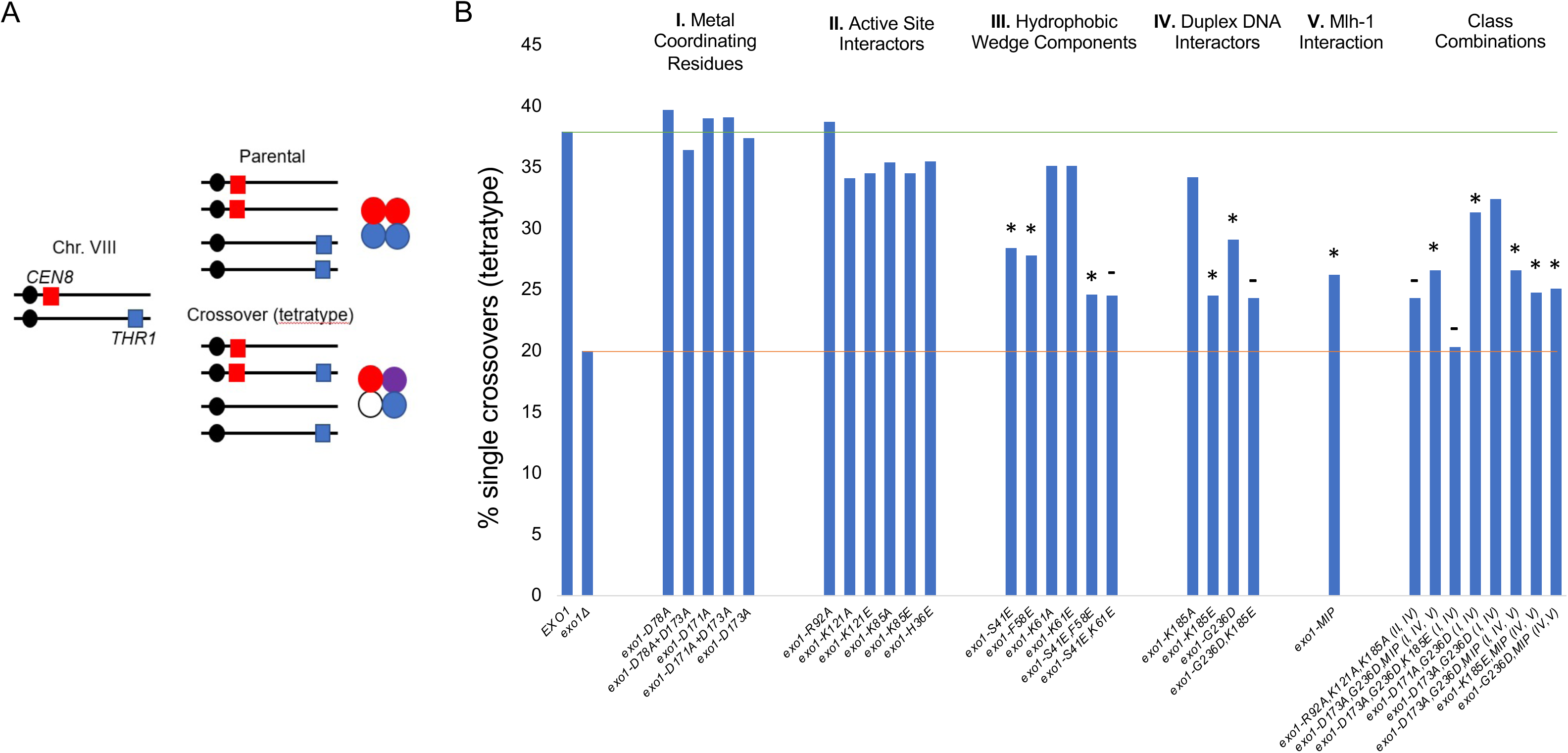
Crossing over for the indicated *exo1* strains was measured in the 20 cM *CEN8* to *THR1* interval on Chr. XV using a spore-autonomous fluorescence assay (Thacker et al., 2011). A. The spore autonomous fluorescence assay was used to measure single meiotic crossover events (tetratypes) in the chromosome VIII *CEN8-THR1* interval (Thacker et al., 2011). B. Single meiotic crossover events in the indicated strains. Mutations are separated into categories based on disruption of specified functions outlined in Figure 1B. *EXO1* and *exo1Δ* levels are indicated by green and red dashed lines, respectively. *, statistically distinguishable from *EXO1* and *exo1Δ;* **-,** distinguishable from *EXO1*, but indistinguishable from *exo1Δ*. See Table S1 for the complete data set.

### Mutation of DNA binding domains of Exo1 reveal a DNA binding role for Exo1 in meiotic crossing over

The structure solved by Orans et al. (2011) revealed that Exo1 makes key contacts with DNA through several defined domains (Figure 1B). For example, G236 (Group IV) is one of several residues in a helix-two turn-helix motif that coordinates a metal ion and forms hydrogen bonds with DNA backbone oxygen residues to stabilize an interaction with Exo1 and the pre-nick duplex DNA. This conserved motif is only slightly modified from observed FEN-1 structures (Ceska et al., 1996; Feng et al., 2004) and is presumed to facilitate exonuclease processivity as the protein moves along the DNA backbone (Pelletier et al., 1996; Orans et al., 2011). K185 is part of a small hairpin loop between strands β6 and β7 and is also thought to be critical for recognition of duplex DNA (Orans et al., 2011; Li et al., 2019). The K185A mutation has been shown to diminish Exo1 nuclease activity several fold *in vitro*, and confer elevated sensitivity to DNA-damaging agents, likely due to a defect in binding duplex DNA (Li et al., 2019). A crucial component of Rad2/XPG members is the hydrophobic wedge (Figure 1B, Group III), a structurally conserved domain which induces a sharp bend at a ds-ssDNA junction, and gives the enzyme family its specificity for gapped/nicked DNA substrates (Orans et al., 2011, Chapados et al., 2004). Several hydrophobic residues within the wedge motif displace the non-substrate strand, as well as two lysine residues which appear to coordinate this portion of the non-substrate strand (Figure 1B).

As shown in Figure 2B and 3B and Tables S1 and S2, the *exo1-K185E* and *exo1-G236D* mutations conferred significant decreases in crossover formation (68 cM, 29.1% tetratype in *exo1-G236D* and 73 cM, 24.5% tetratype in *exo1-K185E*) in the *URA3-HIS3* and *CEN8-THR1* intervals, respectively. Interestingly, the hydrophobic wedge mutations *exo1-S41E* (58.6 cM, 28.4% tetratype), and *exo1-F58E* (69.9 cM, 27.8% tetratype) also conferred crossover defects with double mutation combinations (*exo1-K185E,G236D-*24.2% tetratype*; exo1-S41E,F58E-*24.6% tetratype) conferring more severe phenotypes. We then made a series of double and triple mutants that included a catalytic, DNA binding, and Mlh1-interacting (MIP) mutations (Figure 3B; Table S1). Combining groups did not confer crossover phenotypes equivalent to the *exo1Δ*, and including a catalytic mutation *(-D171A, -D173A*) with any single DNA binding mutation that conferred a crossover phenotype did not further impair crossover formation. However, a triple mutation, *exo1-R92A,K121A,K185A* (24.3% tetratype) conferred a more severe phenotype than the single mutations, and another triple mutation, *exo1-D173A,K185E,G236D* (22.4% tetratype), conferred a phenotype very close to the *exo1Δ*, also suggesting that catalytic mutations could impact DNA binding as indicated above (Figure 3B). The data collected from assaying double and triple mutants validated the results of single catalytic and DNA binding mutations, identified DNA binding mutants that confer a near *exo1Δ* crossover phenotype, and showed that the Exo1 active site is relatively insensitive to mutation for crossover formation. These observations also indicated that the decrease in crossover frequency seen in single mutants is compounded in multiple mutant combinations (Figure 3B).

We then examined the spore viability of *exo1* mutant strains. The *exo1Δ* strain showed a tetrad spore viability pattern (74% spore viability; 4, 2, 0 viable tetrads > 3, 1) consistent with Meiosis I non-disjunction (Figures 2B; S3; Ross-Macdonald and Roeder, 2004; Abdullah et al., 2004). However, decreases in meiotic crossing over and spore viability did not correlate in the *exo1* strains. For example, *exo1* mutants with very similar defects in crossing over showed spore viabilities that ranged from 89% (*exo1-G236D, exo1-MIP*) to 71 to 73% (*exo1-K185E,G236D, exo1-K185E,MIP*). A plausible explanation for these differences is that the *exo1* mutations display other phenotypes in addition to meiotic crossover phenotypes. In fact, some of the *exo1* mutations analyzed above conferred defects in DNA repair, as measured by sensitivity to methyl-methane sulfonate (MMS). However, the MMS phenotypes did not correlate with defects in meiotic crossing over (Figure S4). For example, the *exo1-D78A, exo1-D171A,* and *exo1-D173A* catalytic mutations conferred stronger MMS sensitivities compared to their nearly *wild-type* meiotic CO phenotypes. Similar disparities between DNA repair and CO phenotypes were seen for the active site mutations *exo1-K85E* and *exo1-K121A*, the DNA binding mutant *exo1-K185E* and the MLH interacting mutant *exo1-MIP.* This analysis suggested that the lack of correlation between spore viability and crossover phenotype seen in *exo1* mutants was likely complicated by their defects in DNA repair. Further support for this idea was seen by the lack of a 4, 2, 0 viable tetrads > 3, 1 pattern in the *exo1* mutant alleles, though this pattern was clearly displayed by *exo1Δ* (Figure S3). One explanation for this lack of a pattern in *exo1* mutants with strong crossover defects is that the DNA repair defects in these mutants conferred a pleiotropic decrease in spore viability, obscuring a Meiosis I non-disjunction phenotype. Another potential explanation (discussed below) is that *exo1Δ* strains show increased disjunction as the result of defects in crossover positioning (genetic interference, see below). Together, these observations provide evidence that Exo1 contains distinct DNA repair and meiotic CO functions and DNA binding by Exo1, but not its nuclease activity, is critical for meiotic CO resolution.

### Expression of *RAD27* in meiosis partially complements the crossover defect in *exo1* null strains

The Rad2 family of nucleases consists of four evolutionarily conserved members: *RAD2/XPG* in yeast/humans respectively, *EXO1/EXO1*, *RAD27/FEN-1*, and *YEN1/GEN1.* While all four have distinct roles in DNA metabolism, three members, Exo1, Rad2, and Rad27, possess both 5’ → 3’ exo- and 5’ flap endo-nuclease activity, and Yen1 appears to act exclusively as an endonuclease (Sun et al., 2003, Ip et al., 2008; Tomlinson et al., 2010). In yeast, *RAD27* shares the highest sequence similarly with *EXO1*, suggesting functional overlap. In fact, previous studies have shown that *EXO1* can complement some *RAD27* functions, and the *exo1Δ rad27Δ* double mutant is inviable (Tishkoff et al., 1997, Xie et al., 2001; Qiu et al., 1999). While the substrate preferences of Rad2 family proteins vary, all have been shown to bind nicked, gapped, and/or blunt end DNA, with a particular affinity for single- to double-stranded DNA junctions. They all appear to induce a sharp bend in the DNA substrate upon protein binding (Lee and Wilson, 1999; Genschel and Modrich, 2003; Orans et al., 2011). These observations structurally demonstrate how *RAD2* family proteins can share redundant capacities for endo- and exo-nucleolytic functions.

We reasoned that a protein that mimicked the DNA binding affinity for similar DNA substrates could complement this function in cells lacking Exo1. We therefore tested the ability for Rad27 to complement the meiotic function of Exo1. We did not observe complementation by *RAD27* expressed through its native promoter, but upon placing *RAD27* under control of the *EXO1* promoter (*pEXO1-RAD27)* we saw significant increases in crossing over on both Chromosomes VIII (from 21.5% to 29.9% tetratype; Figure 4A; Table S1B) and XV (Figure 4B; 54 cM map distance in *exo1Δ* to 72 cM *exo1Δ* containing *pEXO1-RAD27*), likely due to the high levels of meiotic expression of the *EXO1* promoter (Figure S5; Brar et al., 2012). Efforts were made to improve *exo1Δ* complementation by fusing a MIP domain, or the entire C-terminus of Exo1 to Rad27 to create a functional Mlh1 interaction; however, they were unsuccessful.

**Figure 4.**
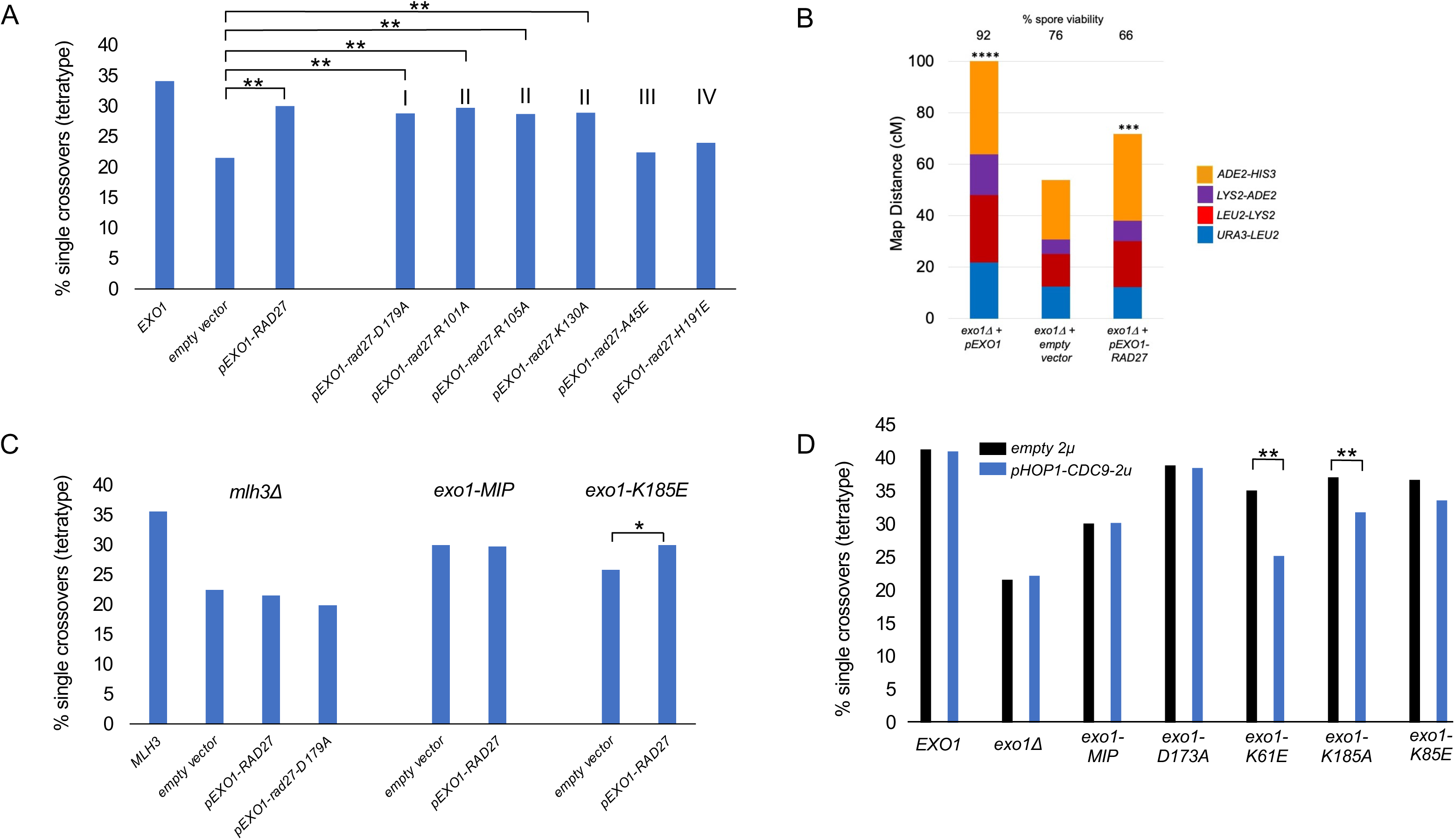
*RAD27* expressed from the *EXO1* promoter can restore crossover functions to *exo1Δ* strains. A. *pEXO1-RAD27*, *ARS-CEN* (pEAA720), the indicated mutant *rad27* derivatives (pEAA724, pEAA727-731), and an empty *ARS-CEN* vector (pRS416), were transformed into an *exo1Δ* strain and examined for crossing over at the *CEN8-THR1* locus. The *rad27* mutations were grouped (I, metal-coordinating; II, active-site; III, hydrophobic wedge; IV, duplex DNA) like those presented for Exo1 (Figure 1B). Significance (*p<0.05; **p<0.01) compared to the *exo1Δ* strain containing an empty vector was determined using a two-tailed Fisher’s Exact Test. B. The *pEXO1-RAD27* plasmid pEAI482 was transformed into *exo1Δ* strains (with *pEXO1, ARS CEN* (pEAI483) and an empty *ARS-CEN* vector (pLZ259) as controls) to measure crossing over in the *URA3-HIS3* interval in the EAY1108/1112 background. Asterisks indicate the number of genetic intervals that are distinguishable from the *exo1Δ* containing the empty vector, as measured using standard error calculated through Stahl Laboratory Online Tools (https://elizabethhousworth.com/StahlLabOnlineTools/; Table S2). C. *mlh3Δ* and the indicated *exo1* strains were transformed with *pEXO1-RAD27* (pEAA720), *pEXO1-rad27-D179A* (pEAA724) and empty vector (pRS416), and examined for crossing over at the *CEN8-THR1* locus. Significance (*p<0.05) compared to the *exo1Δ* strain containing an empty vector (panel A) was determined using a two-tailed Fisher’s Exact Test. D. *CDC9* overexpression in meiosis disrupts the crossover functions of *exo1* DNA binding mutants. Strains with the indicated *exo1* genotypes (Table S5) were transformed with a *2μ URA3* vector containing no insert (*empty 2μ,* pRS426) or *CDC9* expressed from the *HOP1* promoter (*pHOP1-CDC9*, *2μ*, pEAM329) and then assessed for meiotic crossing over in the *CEN8-THR1* interval. Significance is shown between each *empty vector-pHOP1-CDC9* pair using a two-tailed Fisher’s Exact Test, with ** indicating p<0.01.

We reasoned that if Rad27 complemented the meiotic role of Exo1 by binding a specific DNA substrate based on structural similarity divorced from catalytic activity, inactivating Rad27 through mutation of a metal-coordinating aspartic acid D179 (Shen et al., 1996; Gary et al., 1999) would not impact its ability to effect higher crossover frequencies. Indeed, *exo1Δ* cells expressing *pEXO1-RAD27* or *pEXO1-rad27-D179A* showed similar levels of crossover complementation. This observation encouraged us to further test our hypothesis by making five additional *rad27* mutations based on previous biochemical and structural characterization of the human homolog of Rad27, FEN-1. These included *rad27-R101A*; equivalent to *FEN1-R100A*, of which the mutant FEN-1 protein exhibited a strong catalytic defect but remained competent for flap binding and bending (Song et al., 2018), and *rad27-R105A* and *rad27-K130A*, equivalent to FEN-1-R104A and FEN-1-K132A, of which the mutant FEN-1 proteins exhibited 20- and 5-fold reductions in flap cleavage but were not characterized for flap binding or bending (Tsutakawa et al., 2017). Two other mutations were analyzed based on Exo1 and Rad27 homology: *rad27-A45E*, which aligns to a mutation in the Exo1 hydrophobic wedge (*exo1-S41E,* Group III, Figure 1B), and *rad27-H191E*, which aligns to a mutation in the Exo1 DNA binding domain (e*xo1-K185E,* Group IV). As shown in Figure 4A, *rad27-R101A, rad27-R105A and rad27-K130A*, which coordinate the scissile bond for catalysis, complemented the crossover defect in *exo1Δ*, consistent with the phenotypes exhibited by *exo1* Group II mutations. Interestingly, the *rad27-A45E* and *rad27-H191E* mutations were defective in *exo1Δ* complementation, as predicted for their requirements in flap bending and stabilizing the DNA backbone, respectively.

We also tested if *RAD27* expression from the *EXO1* promoter could improve meiotic crossover functions of *exo1* strains bearing mutations within (*exo1-K185E*) or outside of the DNA binding domain (*exo1-MIP*). As shown in Figure 4C, meiotic crossing over in *exo1-K185E,* but not *exo1-MIP,* was increased in cells containing *pEXO1-RAD27*. These observations are consistent with Rad27 being able to substitute for Exo1 DNA binding functions because improved complementation by *pEXO1-RAD27* was seen in a DNA binding mutant (*exo1-K185E*) but not in a mutant predicted to be functional for DNA binding (*exo1-MIP)*, but defective in interacting with other crossover factors.

Finally, we saw no complementation of meiotic crossing over by *pEXO1-RAD27* in strains lacking functional Mlh1-Mlh3 (*mlh3Δ),* indicating that Rad27 complementation was specific to Exo1 function. This observation differs from observations made by Arter et al. (2018), who found that expression of the Rad2/XPG nuclease Yen1 complemented crossover defects in both *exo1Δ* and *mlh3Δ* strains. One explanation for the Yen1 complementation phenotype is that Yen1 Holliday junction resolvase activity could bypass Mlh1-Mlh3-Exo1 dependent dHJ resolution steps.

### Meiotic crossover phenotype of *exo1* DNA binding mutants is significantly reduced when Cdc9 ligase is overexpressed in meiosis

Reyes et al. (2021) et al. recently showed that overexpression of the budding yeast ligase Cdc9 disrupted DNA mismatch repair through the premature ligation of replication-associated nicks that act as critical repair signals. If the role of Exo1 in meiotic recombination involved nick binding/protection, then we reasoned that meiotic overexpression of *CDC9*, the budding yeast DNA ligase involved in DNA replication, could lead to premature ligation of DNA synthesis-associated nicks critical for maintaining biased resolution. We posited that some *exo1* DNA binding mutants that maintained near wild-type levels of crossing over might be especially susceptible to Cdc9 overexpression. During meiosis *CDC9* expression appears to be low relative to *HOP1*, whose expression increases dramatically in meiotic prophase and remains high through dHJ resolution (∼6hrs in meiosis; Figure S5). We thus expressed *CDC9* under control of the *HOP1*. As shown in Figure 4D we saw no disruption of crossing over in *exo1* mutants that contained intact DNA binding domains (*EXO1, exo1-MIP, exo1-D173A)* or in a mutant *(exo1-K85E)* predicted to be defective in steps post-DNA bending (Orans et al., 2011). However, we saw modest to severe losses of crossing over in *exo1* DNA binding mutant hypomorphs. As shown in Figure 4D, *pHOP1-CDC9* reduced single crossovers in *exo1-K185A* from 35.3 to 31.3% and in *exo1-K61E* from 35.1 to 25.2%. These data, in conjunction with the *RAD27* complementation experiments, provide evidence for a nick protection role for Exo1 in crossover formation.

### Interference analysis suggests a role for Exo1 prior to crossover resolution

While expression of *RAD27* under the *EXO1* promoter (*pEXO1-RAD27* plasmid) could partially complement CO defects in *exo1Δ* strains, it did not improve the meiotic spore viability or MMS resistance seen in *exo1Δ* strains (Figures 4B). We performed crossover interference analysis to determine if *exo1Δ* strains showed defects in addition to those seen in DSB resection and CO resolution. As described below, we found that *exo1Δ* strains displayed crossover interference defects that were not complemented by the *pEXO1-RAD27* plasmid.

First, we analyzed *exo1Δ* strains bearing *pEXO1-RAD27* for defects in crossover interference on chromosome XV using the Malkova method, which calculates genetic distances between intervals in the presence and absence of a neighboring crossover (Figure 5; Table S3; Malkova et al., 2004; Martini et al., 2006). These measurements are presented as a ratio, wherein 0 indicates complete interference and 1 indicates no interference. Three pairs of intervals (*URA3-LEU2-LYS2, LEU2-LYS2-ADE2, LYS2-ADE2-HIS3)* were tested for interference. In all three interval pairs tested, *exo1Δ* displayed a loss of interference compared to *wild-type*. Most strikingly, two intervals that displayed strong interference in *wild-type* strains (Malova ratios of 0.48 at *URA3-LEU2-LYS2* and 0.43 at *LEU2-LYS2-ADE2)* displayed a complete loss of interference in *exo1Δ* (1.28 and 0.84 respectively). These results are reminiscent of the interference defects observed previously in *msh4Δ* and *msh5Δ* (Ross-Macdonald and Roeder, 1994; Hollingsworth et al., 1995; Novak et al., 2001; Nishant et al., 2010; Figure 5). Interestingly, a lack of interference was observed in all three intervals in the *exo1Δ* strain containing *pEXO1-RAD27* (Malkova ratios of 1.41, 0.90, and 0.81 in Intervals I, II, III, respectively; Figure 5), supporting the idea that *RAD27* expression in meiosis could complement only Exo1’s crossover functions.

**Figure 5.**
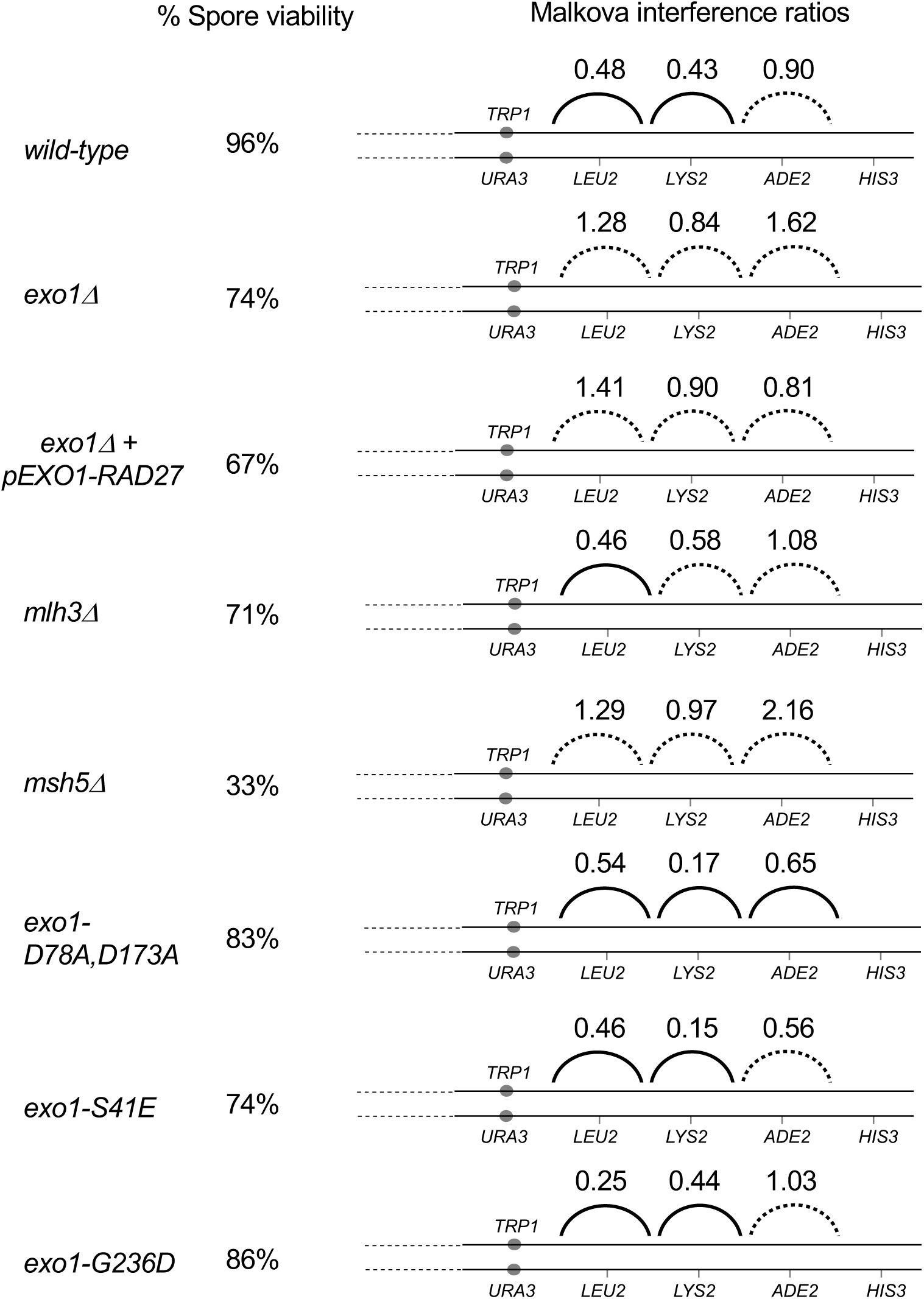
Interference Analysis for pairs of adjacent genetic intervals on Chromosome XV in the EAY1108/EAY1102 strain background. Crossover interference was analyzed on Chromosome XV by measuring centimorgan (cM) distances in the presence and absence of a neighboring crossover (Malkova et al., 2004; Martini et al., 2006; Tables S3A, S3B). Malkova interference is presented as a ratio of cM crossover absent/cM crossover present. *Dashes indicate no detectable positive interference. Significance of differences in tetrad distribution was assessed using a G test. Statistically significant p values (p <0.05) suggest the presence of interference (I) in the genetic interval (Tables S3B).

The interference defect seen in *exo1Δ* (all three intervals showed a lack of interference) was stronger than that seen in the *mlh3Δ* strain (two intervals showed a lack of interference), suggesting a role for Exo1 in promoting interference independent from its association with Mlh1-Mlh3 in crossover resolution. To determine if the early resection role of Exo1 (Zahkaryevich et al., 2010) could account for this interference function, e*xo1-D171A,D173A* and *exo1-D78A,D173A* catalytic mutants were analyzed for interference defects (Figure 5). Strikingly, these mutants displayed interference similar to or stronger than *wild-type*. In fact, the interference defect observed in *exo1Δ* was not recapitulated in any of the *exo1* alleles tested. Interference was also measured using the COC (Coefficient of Coincidence) method (Papazian, 1952; Table S3A). COCs measure the double crossover rate compared to the expected rate in the absence of interference. The COC ratios were consistent with the Malkova ratio analysis, supporting the idea that loss of interference in *exo1Δ* was not recapitulated in any of the mutant alleles. Together the data indicate a previously uncharacterized role for Exo1 in establishing crossover interference and suggest that the pro-interference role of Exo1 is either more robust than the pro-crossover role or involves specific contact or interaction sites that were not examined in this study (see Discussion).

### Genetic interactions involving Msh4-Msh5, Mlh1-Mlh3 and Exo1 also support roles for Exo1 in crossover interference

The finding that *exo1Δ* showed defects in crossover interference encouraged us to determine if we could identify genetic interactions involving factors that interact with Exo1 and play roles in crossover interference. To initiate this work we analyzed *exo1-F447A,F448A* (referred to as *exo1-MIP*), which contains mutations in an Mlh1-interacting peptide box (MIP) that disrupt both Mlh1-Exo1 interactions and meiotic crossing over (Tran et al., 2007; Zakharyevich et al., 2010). In the spore autonomous fluorescence assay we found that the *exo1-MIP* mutation conferred intermediate defects in CO formation (33.3% single crossovers (tetratype) compared to 37.5% in *wild-type*) when both this allele and *MLH3* were present in two copies (Figure S6; Table S4). However, when both *exo1-MIP* and *MLH3* were present in single copies, we observed a two-fold reduction in CO levels (to 22.6% tetratype) that approached levels seen in *mlh3∆* (Figure S6). This observation confirmed interactions between Mlh1-Mlh3 and Exo1 and encouraged us to use gene dosage as an approach to identify additional genetic interactions involving Exo1 using *mlh3* alleles, *mlh3-42* and *mlh3-54,* that confer defects in Mlh3-mediated mismatch repair (MMR) but do not disrupt crossing over. Previous work showed that the *mlh3* alleles disrupted Mlh1-Mlh3 interactions (Al-Sweel et al., 2017). We reduced the gene dosage of eleven meiotic genes from two to one and measured crossing over at the *CEN8-THR1* interval on chromosome VIII (Figure S6; Table S4). *SGS1* and *RMI1* were included because they encode components of a Sgs1-Top3-Rmi1 complex that acts as a pro-crossover factor in meiotic recombination (Jessop et al., 2006; Zakharyevich et al., 2012; Kaur et al., 2015).

As shown in Figure S6 and Table S4, we observed defects for both *mlh3* alleles in crossing over when the gene dosage of *EXO1, MSH4,* or *MSH5* was reduced to one copy. For *MLH1*, we observed such dosage effects with only the *mlh3-54* allele, and for *SGS1* and *RMI1*, with only the *mlh3-42* allele (Figure S6). Interestingly, the residues mutated in *mlh3-54* mapped to the Mlh1-Mlh3 dimerization interface whereas residues mutated in *mlh3-42* mapped to the distal periphery of the dimerization interface (Dai et al., 2021). While this observation might help explain the different effect of gene dosage for *MLH1* in *mlh3-42* and *mlh3-54* backgrounds, it is unclear why the *mlh3-42* allele disrupts the stability of Mlh1-Mlh3 or why it showed gene dosage interactions with *SGS1* and *RMI1*.

*mlh3* allele-specific interactions were not observed when reducing dosage for a group of ZMM family genes (*ZIP1, ZIP3, ZIP4, SPO16, MER3*) which are thought to act upstream of Mlh1-Mlh3 to stabilize early recombination intermediates and promote CO outcomes (Agarwal and Roeder, 2000; Snowden et al., 2004; Borner et al., 2004; Kolas et al., 2005; Argueso et al., 2004; Shinohara et al., 2008; Hatkevich and Sekelsky, 2017). As shown in Figure S6, a reduction of gene dosage for *ZIP1* and *SPO16* did not alter crossing over in any *MLH3* background, and a reduction of dosage for *ZIP3* and *MER3* led to CO decreases in *MLH3, mlh3-42,* and *mlh3-54* backgrounds. *ZIP4* fit a somewhat similar pattern to *ZIP3* and *MER3*, but statistical significance was mixed, with significance for haploinsufficiency seen in only the *mlh3-42* background. Together, these studies support a model in which Msh4-Msh5, Mlh1-Mlh3, and Exo1 form a group that participates in crossover interference (Santucci-Darmanin et al., 2002; Santucci-Darmanin et al., 2000; Zakharyevich et al., 2010; Krishnaprasad et al., 2021).

### Msh5 DNA interactions and foci are not dependent on Exo1

Crossover interference involves the recruitment of ZMM proteins which stabilize and identify a set of dHJs for Class I crossover resolution. Among this class of factors is Msh4-Msh5, which stabilizes SEIs after strand invasion (Boerner et al., 2004). During meiosis, the Msh4-Msh5 complex binds *in vivo* to DSB hotspots, chromosome axes, and centromeres (Krishnaprasad et al., 2021). We previously showed Msh5 can bind resected DSB structures *in vivo* in a mutant defective in strand invasion (*dmc1Δ* mutant; Krishnaprasad et al., 2021). Meiotic DSB resection by Exo1 results in the formation of extensive 3’ overhangs that can promote strand invasion and joint molecule formation stabilized by ZMM proteins (Zakharyevich et al., 2010). However, previous studies have shown that in *exo1Δ*, joint molecule formation is normal, though there is a roughly 50% reduction in crossovers (Khazanehdari and Borts, 2000; Tsubouchi and Ogawa, 2000; Zakharyevich et al., 2010). Since interference and crossover formation is significantly reduced in *msh5Δ*, an explanation for the interference defect in *exo1Δ* is that Msh4-Msh5 recruitment to recombination intermediates is compromised due to reduced resection of DSBs (Zahkaryevich et al., 2010). To address this, we analysed Msh5 binding in an *exo1Δ* mutant using a combination of ChIP-qPCR and cytological methods.

We performed ChIP-qPCR analysis of Msh5 binding in *exo1Δ* at the representative DSB hotspots (*BUD23*, *ECM3*, *CCT6*), chromosomal axes (*Axis I*, *Axis II* and *Axis III*), centromeres (*CENIII*, *CENVIII*), and the DSB coldspot (*YCRO93W;* Krishnaprasad et al., 2021). Enhanced Msh5 binding was observed in *exo1Δ* at some of the representative DSB hotspots (*ECM3, CCT6*) at 4h and 5h relative to the *wild-type* (Figure 6A). Msh5 binding at the axes and centromeres in *exo1Δ* was similar to *wild-type* from 3-5 hrs (Figure 6A).

**Figure 6.**
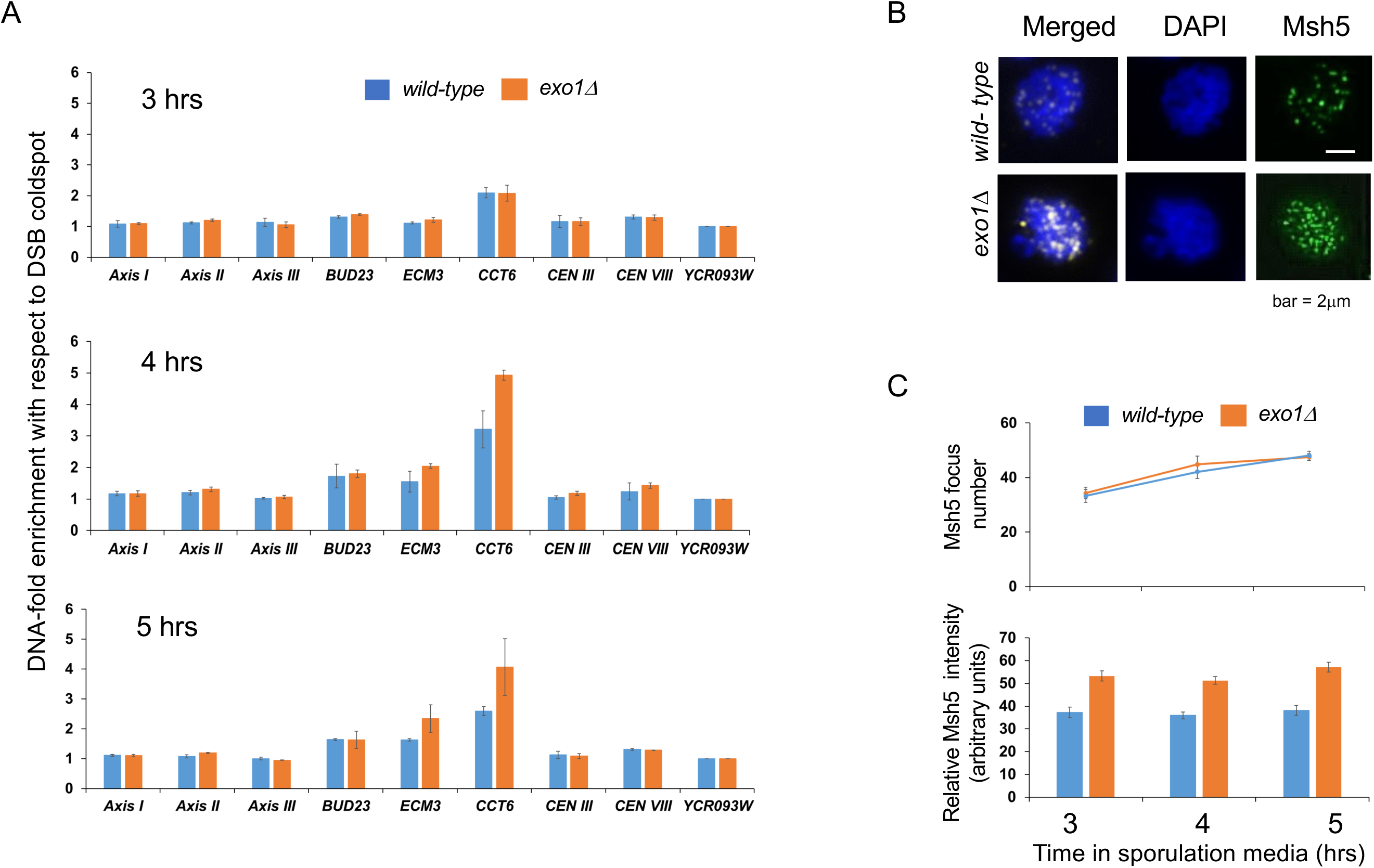
Msh5 localization to chromosomes in *wild-type* and *exo1Δ* strains. A. ChIP-qPCR analysis of Msh5 binding at DSB hotspots (*BUD23*, *ECM3*, and *CCT6*), centromere regions (*CEN III*, *CEN VIII*) and axis regions (*Axis I*, *Axis II*, *Axis III*) relative to DSB coldspot (*YCR093W*) in *wild-type* and *exo1Δ* at 3, 4, and 5 hrs after transfer of cells to sporulation media (see Krishnaprasad et al., 2021 for region assignment). The samples are normalized using input and plotted after dividing with the cold spot value. Error bars represent the standard deviation from two independent biological replicates. B. Representative images of Msh5 staining of chromosome spreads of wild-type and the *exo1* mutant cells at 5-hr incubation in sporulation media. Msh5, green; DAPI, blue. Bar indicates 2 μm. C, top; number of Msh5 foci was counted in Msh5-focus positive spreads at the indicated times. At each time point, 30 nuclei were counted. Mean+/- standard deviation of three independent time courses are shown. C, bottom; relative ratio of Msh5 intensity to DAPI intensity was quantified. At each time point, 30 Msh5-positive nuclei were analyzed. Mean+/- standard deviation of three independent time courses are shown.

Msh5 binding in *exo1Δ* was also analysed by cytological analysis of Msh5 foci (Figure 6B). The average numbers of Msh5 foci per cell in *exo1Δ* at 3 hrs (34), 4 hrs (45) and 5 hrs (48) were comparable to the number of Msh5 foci in *wild-type* at the same time points (33, 42, and 48 respectively) (Figure 6C). However, measurement of the foci intensity showed that the Msh5 foci appeared brighter in *exo1Δ* (Figure 6C). These observations support the ChIP-qPCR data showing enhanced Msh5 binding in *exo1Δ* mutants, especially at DSB hotspots. Together the ChIP and Msh5 localization studies suggest that Msh4-Msh5 localization is not dependent on either the long-range resection activity of Exo1 or interaction with Exo1. This information, in conjunction with interference analysis of *exo1* nuclease defective mutants supports a direct role for Exo1 in establishing interference.

## DISCUSSION

In this study we identified a critical function for Exo1 in meiotic crossing over dependent on its ability to bind to nicked/flapped DNA structures. This conclusion is supported by the finding that meiotic expression of the structurally similar *RAD2* family nuclease Rad27 can partially compensate for the loss of crossovers in the absence of Exo1, and that meiotic overexpression of the Cdc9 ligase conferred a significant crossover defect in *exo1* DNA binding domain mutants. Based on these observations we propose that Exo1 acts in meiotic crossover formation by binding to nicks/flaps analogous to those created during lagging strand DNA synthesis (Figure 7). In contrast to the functions of Rad27 and Exo1 during replication, which cleave 5’ flaps in mechanisms that facilitates ligation of the resulting nick (Balakrishnan and Bambara, 2013), the Exo1/Rad27 meiotic crossover function occurs independently of nuclease activity. Such a nuclease-independent activity likely serves to protect nicks or flaps in recombination intermediates from premature ligation, ensuring their incorporation into a resolution mechanism. In addition, a nick/flap bound Exo1 could act to recruit Mlh1-Mlh3 to the dHJ. In support of this idea, work by Manhart et al. (2017) showed that the presence of Mlh1-Mlh3 polymer at a nicked strand can direct the endonuclease to cut the opposite strand, providing a possible mechanism for how biased resolution could occur.

**Figure 7.**
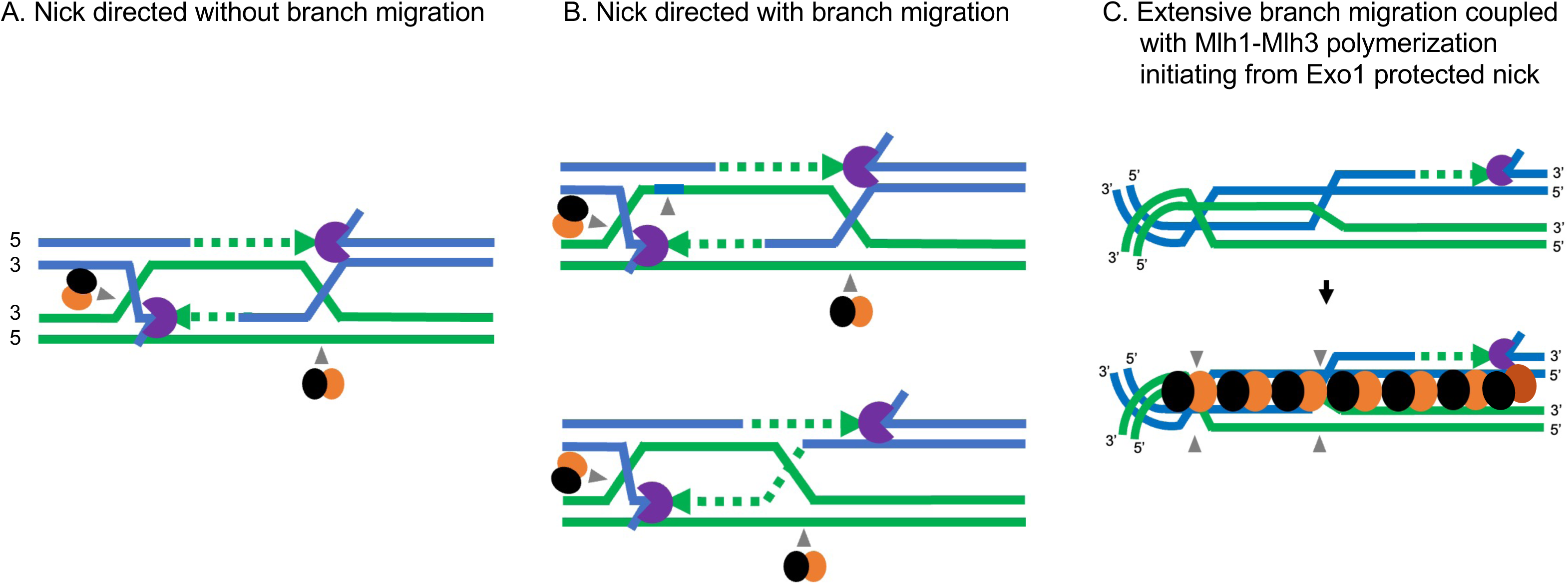
Models for biased resolution of double Holliday junctions. A. Canonical model. In the major interference-dependent crossover pathway, a D-loop intermediate is stabilized by ZMM proteins including Msh4-Msh5 to form a single end invasion intermediate. DNA synthesis from the SEI, followed by second-end capture, results in the formation of the dHJ intermediate which is stabilized by Msh4-Msh5. Biased resolution of the two junctions results in crossover formation. In this model, Exo1 protection of the nick/flap structure recruits Mlh1-Mlh3 to nick the DNA strand opposite the Exo1 protected nick. B. dHJ resolution through limited branch migration, focusing on models adapted from Marsolier-Kergoat et al. (2018; upper panel) and Peterson et al. (2020; lower panel). In these models one or both junctions of the dHJ move prior to resolution. In our adaptation of the Peterson et al. (2020) model, Exo1-protection of nicks recruits Mlh1-Mlh3 as in panel A. In our adaptation of the Marsolier-Kergoat et al. (2018) model, Exo1 protects nicks made by nick translation (resolution independent nicks) and recruits Mlh1-Mlh3 as in panel A. C. dHJ resolution through extended branch migration (Ahuja et al., 2021). Branch migration creates a substrate for Mlh1-Mlh3 polymerization (Manhart et al., 2017). In such a model, the signaling imposed by the binding of Exo1 to nicks acts at a distance. Mlh1-Mlh3 is recruited by Exo1 and forms a polymer with a specific polarity that can displace other factors or be activated upon interaction with such factors. The polymer is activated to introduce a nick on one strand of the duplex DNA on Type II dHJs when it forms a critical length required for stability. See text for details.

### Incorporating nick-protection with models of dynamic dHJs

A role for a nicked recombination intermediate in forming meiotic crossovers has been proposed for many years, with a summary of a few studies provided below. 1. Electron microscopy studies of Holliday junction structures purified from yeast cultures in pachytene failed to reveal open centers expected of fully ligated junctions (Bell and Byers, 1983), though the structure of dHJs *in vivo* is not well understood, and so we cannot exclude the presence of factors that allow centers in fully ligated junctions to open. 2. Nicked HJs are favorable substrates for resolution by resolvase proteins *in vitro* (Fricke et al., 2005), and nicked HJs comprise a large proportion of Holliday junction structures observed in mutants defective in the structure-selective nucleases Yen1 and Mms4-Mus81, suggesting that they represent mitotic recombination intermediates (Garcia-Luis and Machin, 2014). 3. Whole genome sequencing of meiotic spore progeny inferred that the resolution of dHJs is biased towards new DNA synthesis tracts, implying that these tracts contain distinguishing features such as nicks (Marsolier-Kergoat et al., 2018). 4. Biochemical studies have led to models in which nicks persisting during dHJ formation could provide a substrate for continued loading of MMR/replication factors implicated in dHJ resolution (e.g. RFC, PCNA, Msh4-Msh5; Kulkarni et al., 2020; Cannavo et al., 2020). Furthermore, Kulkarni et al. (2020) and Cannavo et al. (2020) showed that PCNA, which is loaded onto primer template junctions during DNA replication, promotes nicking by Msh4-Msh5 and Mlh1-Mlh3. The above observations, however, are challenging to reconcile with observations in *S. cerevisiae* indicating that single strands of DNA within dHJs appear to be continuous (at least at the resolution of denaturing alkaline gels; Schwacha and Kleckner, 1994, 1995) and dHJs are much more dynamic than predicted based on the canonical DSB repair model (Marsolier-Kergoat et al., 2018; Peterson et al., 2020; Ahuja et al., 2021; Figure 7A). However, it is possible that nicked recombination intermediates are not detected because they are transient, yet able to provide the signals critical for crossover formation, such as loading of PCNA.

dHJs have often been portrayed as static intermediates, constrained to the location of the initiating DSB (Figure 7A). While the nick protection mechanism proposed here can be understood in the context of a canonical model in which Exo1 recruits Mlh1-Mlh3 to nick the single-stranded DNA opposite the Exo1 protected nick (Figure 7A), recent work indicated that dHJs undergo significant branch migration *in vivo*. Recently Marsolier-Kergoat et al. (2018), Peterson et al. (2020), and Ahuja et al. (2021) showed in meiosis that one or both junctions of the dHJ can move independently or in concert prior to resolution. Marsolier-Kergoat et al. (2018) estimated the frequency of branch migration to be on the order of 28%, and Ahuja et al. (2021), based on a detailed analysis of a well-defined recombination hotspot containing a high density of single nucleotide polymorphisms, inferred that ∼50% of crossovers occurred in locations where both HJs are located on one side of the initiating DSB, with a much higher number of crossovers showing some migration.

How can nick protection be incorporated into crossover mechanisms that involve branch migration of HJs? One possibility is that nicks are translocated through “nick translation” (Marsolier-Kergoat et al., 2018). For certain types of branch migration, this mechanism would push the nicks to a new dHJ location, allowing bias to be maintained (Figure 7B, upper panel). In one such model (Marsolier-Kergoat et al., 2018), Exo1 nick protection would occur when DNA synthesis encounters a 5’ end and resolution by Mlh1-Mlh3 would occur (Figure 7B). Alternatively, Mlh1-Mlh3 could nick at a distance from the Exo1-protected nick (Peterson et al., 2020, Figure 7B, lower panel), which could be reconciled based on previous studies showing that MLH proteins form polymers on DNA and can make multiple nicks on DNA (Hall and Kunkel, 2001; Manhart et al., 2017; Kim et al., 2019). In the Marsolier-Kergoat (2018) model, the synthesis of new DNA tracts has been hypothesized to be followed by processing of the resultant 5’ flap to create a nick. Though appealing, this model needs to be balanced with our findings that the catalytic activity of Rad27 is not necessary to rescue crossing over in an *exo1Δ* strain.

A key aspect of extensive branch migration is that it should prevent DNA nicks from serving as substrates for biased resolution because they locate away from the resolution site. To reconcile this observation with our analysis of Exo1, such nicks could act as substrates for the activation of an Mlh1-Mlh3 polymer (Figure 7C). Previous work showed that Mlh1-Mlh3 requires a large DNA substrate for nuclease activation and that polymerization barriers impeded its nuclease activity (Manhart et al., 2017). As such, branch migration may provide a way to move the dHJ from a constrained state that is occupied by factors that establish the dHJ such as Msh4-Msh5. In such a model, the signaling imposed by the binding of Exo1 to nicks could act across a distance, and through an initial Exo1-Mlh1-Mlh3 interaction, allowing the Mlh1-Mlh3 polymer to occupy the comparatively unconstrained DNA away from the invasion site (Figure 7C). Thus, we may consider the Exo1-nick interaction site as a nucleation point for Mlh1-Mlh3. This would add asymmetry to the polymer and ensure that Mlh1-Mlh3 nicks in a biased manner. We illustrate this within the context of a model presented by Manhart et al. (2017), in which Mlh1-Mlh3 requires polymerization across multiple kilobases to be catalytically active to cleave Type II Holliday junctions. Variations of such a model have been presented by Kulkarni et al. (2020). These models would also provide an explanation for the importance of Exo1-Mlh1-Mlh3 interactions during meiotic crossing over (but see below). In this model, we see Exo1-nick interactions as a means of guarding essential nicks from premature ligation. This would ensure that the dHJ remains “flexible” if needed for Mlh1-Mlh3 polymerization and activation. These models are not mutually exclusive, and further work is required to understand how resolution factors interact with mobile and static dHJs.

An additional challenge with the models presented in Figure 7 is that while Exo1 and FEN-1 bind flap structures to coordinate tail removal and ligation steps, the endonuclease activities of these proteins do not appear to be required for crossover resolution. However, the finding that ligase overexpression can disrupt crossing over in *exo1* DNA binding hypomorphs suggests that a ligatable nick serves as a critical recombination intermediate. One possibility is that there is a coordinated displacement of Exo1 by Mlh1-Mlh3 that induces Mlh1-Mlh3 nicking on the opposite strand. In such a model there could be other processing events that removal 5’ tails such as one involving Msh2-Msh3 recognition of the flap, followed by endonuclease cleavage by Rad1-Rad10 (Sugawara et al., 1997). It is also worth noting that studies in which we observed complementation of the *exo1Δ* strain with the *pEXO1-rad27-D179A* plasmid contained native *RAD27* that could act to remove 5’ tails.

Does Exo1 direct Mlh1-Mlh3 nicking? A coordinated set of steps are required in meiotic recombination to promote Exo1 mediated resection of DSBs, D-loop formation, DNA polymerase mediated synthesis of the invading 3’ strand, Exo1 protection of flaps/nicks, and ligation of cleaved dHJs. The transitions between these steps are likely to proceed through mechanisms that involve post-translational modifications (e.g. Bhagwat et al., 2021). Recent studies have shown that Exo1 has a key role in the activation of Mlh1-Mlh3 through Cdc5 Kinase (Sanchez et al., 2020), and a protein association/mass spectrometry study (Wild et al., 2019) suggested that Mlh1-Mlh3 meiotic interactions with Exo1 are dynamic. However, we and others have shown that the *exo1-MIP* mutant defective in Mlh1 interactions displays an intermediate defect in meiotic crossing over (Figure S6; Zahkaryevich et al. 2010), suggesting the possibility of other factors/structures facilitating Mlh1-Mlh3 endonuclease activation. Consistent with this, Mlh1-Mlh3 foci appear to form in meiotic prophase in the absence of Exo1 (Sanchez et al., 2020) and *RAD27* complementation of the *exo1Δ* crossover defect was not complete and did not improve crossover interference (Figure 4). One mechanism consistent with the above observations is that a DNA structure or protein barrier forms during meiotic recombination that activates the Mlh1-Mlh3 endonuclease, analogous to that seen for activation of Type I restriction enzymes through head-on collision of two translocating enzymes. (Szczelkun, 2002). Understanding how these transitions occur will require both *in vitro* reconstruction studies using purified proteins and novel *in vivo* approaches to identify nicks in dHJ intermediates.

### A role for Exo1 in promoting genetic interference

In baker’s yeast the ZMM factor Zip3 has been shown to be an early marker for crossover designation and interference, prior to the formation of physical crossovers, and previous work has suggested that crossover interference and crossover assurance are carried out as distinct functions by the ZMMs (Shinohara et al., 2008). These observations indicate that crossover interference is established prior to dHJ resolution (reviewed in Zhang et al., 2014). Interestingly, while *mlh3Δ* mutants lose dHJ resolution bias, residual interference in *mlh3Δ* mutants suggest that biased resolution is not required for interference. In contrast, a more severe loss of crossover interference in *exo1Δ* (Figure 5) suggests a role beyond preserving resolution bias by protecting nicks, analogous to ZMM proteins which designate crossovers and assure interference on the maturing dHJ. The interference role for Exo1 was also reflected in spore viability patterning, as only the full *exo1Δ* displayed a viability pattern consistent with non-disjunction. While it is not possible to determine precisely how crossover patterning is disrupted in our *exo1Δ* data, the strong interference defect and clear non-disjunction pattern seen in *exo1Δ* strains is consistent with ZMM proteins that work early in imposing interference. The nature of this role remains unclear, as none of the *exo1* alleles tested showed the interference defect seen in *exo1Δ*, and in fact some *exo1* mutants showed increased interference. While Exo1 has been observed to interact with Msh2 through a Msh2-interacting-peptide (SHIP) box, direct interaction with Msh4-Msh5 has not been characterized (Goellner et al., 2018). A link between Exo1 and Msh4-Msh5 is also discouraged by the finding that Msh4-Msh5 localization is not dependent on Exo1 (Figure 6). This observation and previous work showing that joint molecule formation occurs at wild-type levels in *exo1Δ* mutants (Zakharyevich et al., 2010) suggest that the interference defect seen in *exo1Δ* mutants does not reflect the defective loading of Msh4-Msh5 to recombination intermediates.

Could the interference defect seen in *exo1Δ* mutants reflect a defect in resection of DSBs? The enhanced Msh5 association with chromosomes in *exo1Δ* could be interpreted as stabilizing DSB repair intermediates that would normally be eliminated and thus contribute to an interference defect. Several points argue against this idea: 1. *exo1Δ* has reduced crossovers despite increased binding of Msh5 (Figure 6; Khazanehdari and Borts, 2000; Tsubouchi and Ogawa, 2000; Zakharyevich et al., 2010). 2. Msh5 enrichment in *exo1Δ* could reflect compensatory/ homeostatic mechanisms to ensure crossover formation when there is a defect in the processing of recombination intermediates (e.g. Cole et al., 2012). 3. As indicated above, a large number of *exo1* mutants containing mutations in catalytic and DNA binding domains (Figure 5) maintain crossover interference, consistent with defects in DSB resection not being the cause of the interference defect seen in *exo1Δ* mutants. 4. We obtained evidence for a set of genetic interactions involving Exo1, Mlh1-Mlh3 Msh4-Msh5 and Sgs1-Top3-Rmi1 (Figure S6) consistent with Exo1 interaction with genes that are thought to function at both early and later stages in the meiotic crossover resolution pathway. Teasing apart how Exo1 coordinates roles in crossover selection and resolution is critical for understanding how biased resolution of dHJs occurs.

## MATERIALS AND METHODS

### Exo1 homology model

The crystal structure of human Exo1 in complex with 5’ recessed DNA (amino acids 2 to 356; Orans et. al., 2011) was used to map residues in yeast Exo1 critical for function. A homology model was constructed (Figure 1B) using the Phyre2 software (http://www.sbg.bio.ic.ac.uk/phyre2/html/page.cgi?id=index). The predicted structure was aligned to human Exo1 (PDB ID: 3QEB) using Pymol (https://pymol.org/2/). Metal binding residues mutated in this study were D78, D171, and D173. Active site residues mutated were H36, K85, R92, K121. Hydrophobic wedge residues mutated were S41, F58, and K61 and DNA binding residues mutated were K185 and G236. For Figure S1 the Exo1 protein sequence from *S. cerevisiae* was submitted to the BLASTP server at NCBI and run against the landmark database. Protein sequences of Exo1 homologs from different model organisms were analyzed and a multiple-sequence alignment was generated with MAFFT using default settings (Katoh et al., 2018).

### Purification of Exo1

Exo1-FLAG variants (Exo1, exo1-D173A, exo1-G236D, exo1-D173A,G236D) were purified from pFastBac1 constructs (Table S6) in the baculovirus/*Sf9* expression system as described by the manufacturer (Invitrogen) with the following modifications (Nicolette et al., 2010). Briefly, 250 ml of *Sf9* cell pellet was resuspended in 7.5 mL of a buffer containing 50 mM Tris pH 7.9, 1 mM EDTA, 0.5 mM PMSF, 0.5 mM β-mercaptoethanol, 20 μg/mL leupeptin, and 0.25x Halt protease inhibitor cocktail (Thermo). The suspension was incubated on ice for 15 min, after which NaCl was added to a final concentration of 100 mM and glycerol was added to a concentration of 18 % (v/v) and incubated on ice for 30 min. The cells were centrifuged at 30,000xg for 30 min. The cleared lysate was applied to a 2 mL SP Sepharose Fast Flow column at a rate of ∼15 mL/hr. The column was washed with 10 mL of a buffer containing 50 mM Tris pH 7.9, 10 % glycerol, 100 mM NaCl, 0.5 mM PMSF, 5 mM β-mercaptoethanol, and 6.7 μg/mL leupeptin. Exo1 variant was eluted with the above buffer containing 700 mM NaCl. Fractions containing Exo1 protein variant were pooled and applied to 0.3 mL of M2 anti-FLAG agarose beads (Sigma) in batch, incubating with rotation for ∼1.5 hours at 4 °C. Unbound protein was isolated by centrifugation at 2,000 RPM for 5 min in a swinging bucket centrifuge at 4 °C. The resin was resuspended in 7 mL of buffer containing 20 mM Tris pH 7.9, 150 mM NaCl, 10 % glycerol, 0.1 % NP40, 0.5 mM PMSF, 0.5 mM β-mercaptoethanol, 6.7 μg/mL leupeptin, and one-third of a Complete Protease Tablet (Roche) for every 100 mL of buffer and flowed into an empty column at ∼15 ml/hr, allowing to pack. The column was then washed with 0.6 ml of the above buffer excluding the NP40 (wash buffer II). Exo1-FLAG variants were eluted using wash buffer II containing 0.1 mg/mL 3x-FLAG peptide (Sigma). After applying elution buffer, the flow was stopped after the first three fractions were collected and incubated for ∼1 hr before resuming flow and collecting fractions. Fractions containing Exo1 variant were pooled, flash frozen in liquid nitrogen, and stored at -80 °C. All purification steps were performed at 4 °C. Protein concentration was determined by the method of Bradford (1976).

### Endonuclease assays

Exo1 endonuclease reactions were performed on supercoiled 2.7 kb pUC18 or 4.3 kb pBR322 DNA (Invitrogen), or pUC18 DNA nicked by incubation with Nt.BstNBI (New England Biolabs; Rogacheva et al., 2014; Manhart et al., 2017). Briefly, 20 μl reactions (0 to 30 nM Exo1 or mutant derivative) were assembled in a buffer containing 20 mM HEPES-KOH pH 7.5, 20 mM KCl, 0.2 mg/ml BSA, 1% glycerol, and 5 mM MgCl_2_ unless otherwise indicated. Reactions (37°C, 1 hr) were stopped by the addition of a stop mix solution containing final concentrations of 0.1 % SDS, 14 mM EDTA, and 0.1 mg/ml Proteinase K (New England Biolabs) and incubated at 37 °C for 20 min. Products were resolved by 1.2% agarose gel containing 0.1 μg/mL ethidium bromide. Samples were prepared and gels were run as described previously (Manhart et al., 2017). Gel quantifications were performed using GelEval (FrogDance Software, v1.37) using negative control reactions as background.

### Media and yeast strains

*S. cerevisiae* SK1 yeast strains used in this study (Table S5) were grown at 30°C in either yeast extract peptone-dextrose (YPD) or synthetic complete media supplemented with 2% glucose (Rose et al., 1990). When required, geneticin (Invitrogen, San Diego) or nourseothricin (Werner BioAgents, Germany) were added to media at recommended concentrations (Goldstein and McCusker, 1999). Meiotic crossing over was analyzed in the SK1 isogenic background using spore-autonomous assays to measure crossing over in the *CEN8-THR1* interval on Chromosome VIII (SKY3576/SKY3575 parental diploids, Thacker et al., 2011) and in the SK1 congenic EAY1108/EAY1112 background (four intervals on Chromosome XV, Argueso et al., 2004). Sporulation media was prepared as described (Argueso et al., 2004).

### Strain constructions

Mutant alleles were transformed into *S. cerevisiae* with integration plasmids, *geneXΔ::KANMX* PCR fragments or on *CEN6-ARSH4* and 2μ plasmids using standard techniques (Gietz et al., 1995; Rose et al., 1990). To confirm integration events, genomic DNA from transformants was isolated as described previously (Hoffman and Winston, 1987). Transformants bearing *EXO1::KANMX* and *exo1::KANMX* mutant derivatives were screened for integration by analyzing DNA fragments created by PCR using primers AO4061 and AO3838. Integration of *exo1* alleles was confirmed by DNA sequencing of the DNA fragments created by PCR using primers AO3666 and AO3399 (Table S7). To confirm integration of *geneXΔ::KANMX* mutations, primers that map outside of the *geneXΔ::KANMX* PCR fragment were used (Table S7). At least two independent transformants for each genotype were made.

### *exo1* integrating and *EXO1, RAD27* and *CDC9* expression plasmids

Plasmids created in this study are shown in Table S6 and the oligonucleotide primers used to make plasmids are shown in Table S7. Genes expressed in plasmids are from the SK1 strain background (Kane and Roth, 1974).

pEAI422 (4.7 KB; *exo1Δ::KANMX)* was built using HiFi DNA Assembly (New England Biolabs). It contains a complete deletion of the *EXO1* open reading frame but retains 280 bp of 5’ flanking and 340 bp of flanking 3’ sequence. This plasmid was digested with *Spe*I and *Sma*I to release the *exo1Δ::KANMX* fragment prior to transformation.

pEAI423 (7.2KB; *EXO1-KANMX*) contains the entire *EXO1* gene with ∼300 bp of promoter sequence and ∼500 bp of sequence downstream of the stop codon linked to the *KANMX* marker. In this construct, there are ∼300 base pairs of immediate downstream sequence to retain the small gene of unknown function that is immediately found after *EXO1*, followed by *KANMX*, followed by downstream homology. pEAI423 was created using HiFi assembly of the following DNA fragments: 1. *Bam*H1 digested pUC18. 2. An *EXO1* gene fragment made by PCR-amplifying SK1 genomic DNA with primers AO4030 and AO4031. 3. A *KANMX* gene fragment made by PCR-amplifying plasmid pFA6 (Bahler et al., 1998) with AO4032 and AO4033. 4. Downstream *EXO1* sequences made by PCR-amplifying SK1 genomic DNA with AO4034 and AO4035. Integration of this construct confers a *wild-type EXO1* genotype. Derivatives of pEAI423 containing mutations in *EXO1* were constructed with the Q5 mutagenesis kit (New England Biolabs) using pEAI423 as template and the oligonucleotides shown in Table S7. The sequence of the entire open reading frame of *EXO1* in *wild-type* and mutant constructs was confirmed by DNA sequencing in the Cornell Bioresource Center using primers AO275, AO643, AO694, AO804, AO2383, AO3886, AO4028. pEAI423 and mutant derivatives were digested with *Spe*I and *Nhe*I to introduce *EXO1::KANMX* or *exo1::KANMX* fragments into SKY3576 and SKY3575 by gene replacement.

pEAA726 (10.5 KB; *MLH3, CEN6-ARSH4, URA3*) an *MLH3* complementation vector, was created by ligating a *Bam*HI-*Sal*I *MLH3-KANMX* fragment from pEAA636 into the pRS416 (*ARS/CEN, URA3;* Christianson et al., 1992) backbone digested with *Bam*HI and *Sal*I.

pEAA722 (6.4 KB; *RAD27, CEN6-ARSH4, URA3*), a *RAD27* complementation vector, was constructed in two steps. First, a fragment of the *RAD27* gene containing 259 bp upstream and 300 bp downstream sequence was created by PCR amplification of SK1 genomic DNA using primers AO4707 + AO4708. The resulting fragment was digested with *Spe*I + *Kpn*I and ligated into pRS416 digested with *Spe*I + *Kpn*I to create pEAA722.

pEAA715 (7.8 KB; *EXO1, CEN6-ARSH4, URA3*) was constructed in two steps. First, a fragment of the *EXO1* gene containing 400 bp upstream and downstream sequence was created by PCR amplification of SK1 genomic DNA using primers AO4631 and AO4636. The resulting fragment was digested with *Spe*I + *Kpn*I and ligated into pRS416 digested with *Spe*I + *Kpn*I to create pEAA715.

pEAA720 (6.8 KB), a *pEXO1-RAD27* (*EXO1* promoter driving *RAD27* expression*), CEN6-ARSH4*, *URA3* vector, was constructed by HiFi assembly (New England Biolabs) using the following fragments: 1. pRS416 (*CEN6-ARSH4, URA3)* digested with *Kpn*I + *Xba*I. 2.*EXO1* promoter region (400 bp immediately upstream ATG) amplified from the SK1 genome using AO4643 + AO4644. 3. The entire *RAD27* ORF amplified from the SK1 genomic DNA using AO4645 + AO4637. 4. The *EXO1* downstream region (400 bp immediately downstream of the stop codon) amplified from the SK1 genomic DNA using AO4638 + AO4636. *rad27* mutant alleles were constructed with the Q5 mutagenesis kit (New England Biolabs) using pEAA720 as template The oligonucleotides used to make the alleles are shown in Table S7). All *RAD27* plasmid constructs were confirmed by DNA sequencing.

pEAM327 (9.3 KB), a *CDC9, 2μ, URA3* plasmid, was constructed in two steps. First a fragment of the *CDC9* ORF, containing 1000 bp upstream and 400 bp downstream sequence was created by PCR amplification of SK1 genomic DNA using primers AO4783 and AO4784. The resulting fragment was digested with *Hind*III and *Kpn*I and then ligated to pRS426 (*2μ, URA3)* backbone also digested with *Hind*III and *Kpn*I to create pEAM327.

pEAM329 (8.8 KB) is a *2μ, URA3* plasmid that expresses *CDC9* from the *HOP1* promoter (*pHOP1-CDC9*). It was constructed through Hifi assembly using the following fragments: 1. A DNA backbone was created by PCR amplification of pEAM327 using primers AO4837 and AO4838; the resulting DNA fragment lacks the *CDC9* promoter. 2. A 500 bp DNA fragment of the *HOP1* promoter (up until the *HOP1* start codon) was created by PCR amplification of SK1 genomic DNA using primers AO4839 and AO4840. The two fragments were then assembled using Hifi Assembly to create pEAM329, which was confirmed by DNA sequencing.

### Tetrad analysis

Diploids derived from EAY1108/EAY1112 were sporulated using the zero*-*growth mating protocol (Argueso et al., 2003). Briefly, haploid parental strains were patched together, allowed to mate overnight on complete minimal plates, and then struck onto selection plates to select for diploids. The resulting diploids were then transferred from single colonies to sporulation plates where they were incubated at 30°C for 3 days. Tetrads were dissected on minimal complete plates and then incubated at 30°C for 3–4 days. Spore clones were replica-plated onto relevant selective plates and assessed for growth after an overnight incubation. Genetic map distances were determined by the formula of Perkins (1949). Interference calculations from three-point intervals were conducted as described (de los Santos et al., 2001; Novak et al., 2001; Shinohara et al., 2003). Statistical analysis was done using the Stahl Laboratory Online Tools (https://elizabethhousworth.com/StahlLabOnlineTools/) and VassarStats (http://faculty.vassar.edu/lowry/VassarStats.html) and the Handbook of Biological Statistics (http://udel.edu/mcdonald/statintro.html).

Interference was measured by the Malkova method (Malkova et al., 2004). This method measures cM distances in the presence and absence of a neighboring crossover. The ratio of these two distances denotes the strength of interference, with a value closer to 1 indicating a loss of interference. Significance in the distribution of tetrads was measured using a G test (McDonald, 2014) and values of p<0.05 were considered indicative of interference. The coefficient of coincidence (C.O.C) was also measured for each interval by calculating the ratio of observed vs expected double crossovers.

### Spore-autonomous fluorescence assay

We analyzed crossover events between spore-autonomous fluorescence reporter constructs at the *CEN8-THR1* locus on Chromosome VIII (SKY3576, SKY3575; Thacker et al., 2011). To produce diploid strains for analysis in the spore autonomous fluorescence assay, haploid yeasts of opposite mating types were mated by patching together on YPD from freshly streaked colonies and allowed to mate for 4 hrs, and then transferred to tryptophan and leucine dropout minimal media plates to select for diploids. Diploids grown from single colonies were patched onto sporulation plates and incubated at 30°C for approximately 72 hours. Diploid strains containing *ARS-CEN* or *2μ* plasmids were also grown on selective media to maintain the plasmids until just prior to patching onto sporulation plates. Spores were treated with 0.5% NP40 and briefly sonicated before analysis using the Zeiss AxioImager.M2. At least 500 tetrads for each genotype were counted to determine the % tetratype. Two independent transformants were measured per allele. A statistically significant difference from *wild-type* and *exo1Δ* controls based on χ2 analysis was used to classify each allele as exhibiting a wild-type, intermediate, or null phenotype. We applied a Benjamini-Hochberg correction at a 5% false discovery rate to minimize α inflation due to multiple comparisons.

### Sensitivity to methyl-methane sulfonate

Yeast strains were grown to saturation in YPD liquid media, after which they diluted in water and spotted in 10-fold serial dilutions (undiluted to 10^-5^) onto YPD media containing 0.04% MMS (v/v; Sigma). Plates were photographed after a 2-day incubation at 30°C.

### Haploinsufficiency screen

We created knockout transformation PCR fragments consisting of a *KANMX4* antibiotic resistance marker flanked by 300 bp of upstream and downstream homology with respect to the open reading frame (ORF) of each gene of interest. These cassettes were amplified by PCR from genomic preps of the appropriate strains from the *Saccharomyces* genome deletion project (Giaever et al., 2014). In this collection, each ORF has been replaced with *KANMX4*.

EAY3486 (Table S5), a *mlh3∆* strain carrying a gene encoding a cyan fluorescent protein (CFP) on chromosome VIII, was transformed with the PCR amplified knockout cassette. Cells were then plated on YPD-G418 plates and grown at 30°C for three days. At least two independent transformants were verified by confirming resistance to G418 and PCR amplification of using genomic preps of G418 resistant transformants. For PCR verification, primers annealing 350 bp upstream and downstream of the ORF of the gene of interest were utilized to ensure integration at the proper locus. Haploids were then mated to four *MLH3* strains each carrying a gene encoding a red fluorescent protein (RFP) on chromosome VIII. These four strains are as follows: EAY3252 (*MLH3*), EAY3255 (*mlh3∆*), EAY3572 (*mlh3-42*), and EAY3596 (*mlh3-54*). Diploids were isolated by selecting on media lacking tryptophan and leucine and analyzed in the spore-autonomous fluorescence assay described below.

Our criteria for allele-specific interactions was one in which there was little to no change in percent tetratype in either an *MLH3* and *mlh3∆* background, but there was a significant drop of percent tetratype in either *mlh3-42* or *mlh3-54* backgrounds. Significance was assessed by *χ*^2^ test between haplosufficient and haploinsufficient conditions. To minimize α inflation due to multiple comparisons, we applied a Benjamini-Hochberg correction at a 5% false discovery rate (Benjamini and Hochberg, 1995).

### Chromatin immunoprecipitation

Yeast strains KRY753, KTY756, KTY757, NHY1162 and NHY1168 used in the ChiP-qPCR and Msh5 localization analyses (Figure 6) are all derivatives of the *S. cerevisiae* SK1 strain. The *exo1Δ:: KanMX4* marker in KTY753, KTY756 and KTY757 was created using homologous recombination based gene knockout approach in the NHY1162/1168 background (Martini et al., 2006). The transformed colonies were verified by PCR using primers designed for the *EXO1* flanking regions. Msh5 ChIP was performed using polyclonal Msh5 antibody (generated in rabbit) and Protein A Sepharose beads (GE Healthcare) on synchronized meiotic cultures as described in Krishnaprasad et al. (2021). The immunoprecipitated DNA was collected at 3h, 4h, and 5h post entry into meiosis and used for ChIP-qPCR. The DNA enrichment for the Msh5 ChIP-qPCR was estimated with reference to the input at each time point. Msh5 enrichment data for the *wild-type* was from Krishnaprasad et al. (2021). ChIP-qPCR was performed on two independent biological replicates of Msh5 immunoprecipitated DNA samples from *exo1Δ* (3h, 4h, and 5h). Msh5 binding was analyzed at representative DSB hotspots (*BUD23*, *ECM3*, *CCT6*), axes (*Axis I*, *Axis II*, *Axis III*), centromeres (*CENIII, CENVIII*), and DSB coldspot (*YCR093W*). Chromosomal coordinates for these regions and the primer sets used for the qPCR are described in Krishnaprasad et al. (2021).

### Cytological analysis of Msh5 foci

Chromosome spreads (3h, 4h and 5h) were prepared from synchronized meiotic cultures (3, 4 and 5hr) as described (Bishop, 1994; Shinohara et al., 2008; Challa et al., 2019). Msh5 staining was performed using primary antibody against Msh5 (Shinohara et al., 2008) at 1:500 dilution, followed by secondary antibody (Alexa fluor 488, Thermo Fisher Scientific) at 1:1500 dilution. The Msh5 stained samples were imaged using an epi-fluorescence microscope (BX51, Olympus) with a 100X objective (NA,1.3). Images were captured by the CCD camera (CoolSNAP, Roper) processed using iVision (Sillicon) software. To quantify Msh5 focus intensity, the mean fluorescence of a whole nucleus was quantified with Fiji (ImageJ). The final fluorescence intensity of Msh5 was normalized with DAPI intensity for each nucleus. Fluorescence intensity refers to pixel intensity per unit area on chromosome spreads.

## ACKNOWLEDGEMENTS

We are grateful to Michael Lichten, Ilya Finkelstein, Jasvinder Ahuja, and Marcus Smolka for helpful discussions, Scott Keeney for the SKY3576/3576 strains, Michael Liskay for Exo1 expression plasmids, and members of the Alani laboratory for helpful comments throughout this work. M.G. G.P. V.R. J.J.C., M.S., and E.A. were supported by the National Institute of General Medical Sciences of the National Institutes of Health: R35GM134872. L.P. was funded by a Sloan Fellowship, S.M was a student in the Molecular Biology and Genetics Research Experience for Undergraduate (MBG-REU) program, supported by the NSF (DBI1659534), and C.M.M. was a supported by National Institutes of Health grant F32GM112435. K.T.N. was funded by a grant from the Department of Science and Technology (https://dst.gov.in/) (CRG/2018/000916). S.S. was funded by a fellowship from the Council for Scientific and Industrial Research, New Delhi (https://www.csir.res.in), and A.F.F. was supported by a fellowship from IISER TVM (https://www.iisertvm.ac.in/). A.S. was supported by a Grant-in-Aid from the JSPS KAKENHI (19H00981). The content of this work is solely the responsibility of the authors and does not necessarily represent the official views of the National Institutes of Health.

## COMPETING INTERESTS

The authors declare no competing interests.

## SUPPLEMENTARY MATERIALS

### Figure Legends

**Figure S1.**
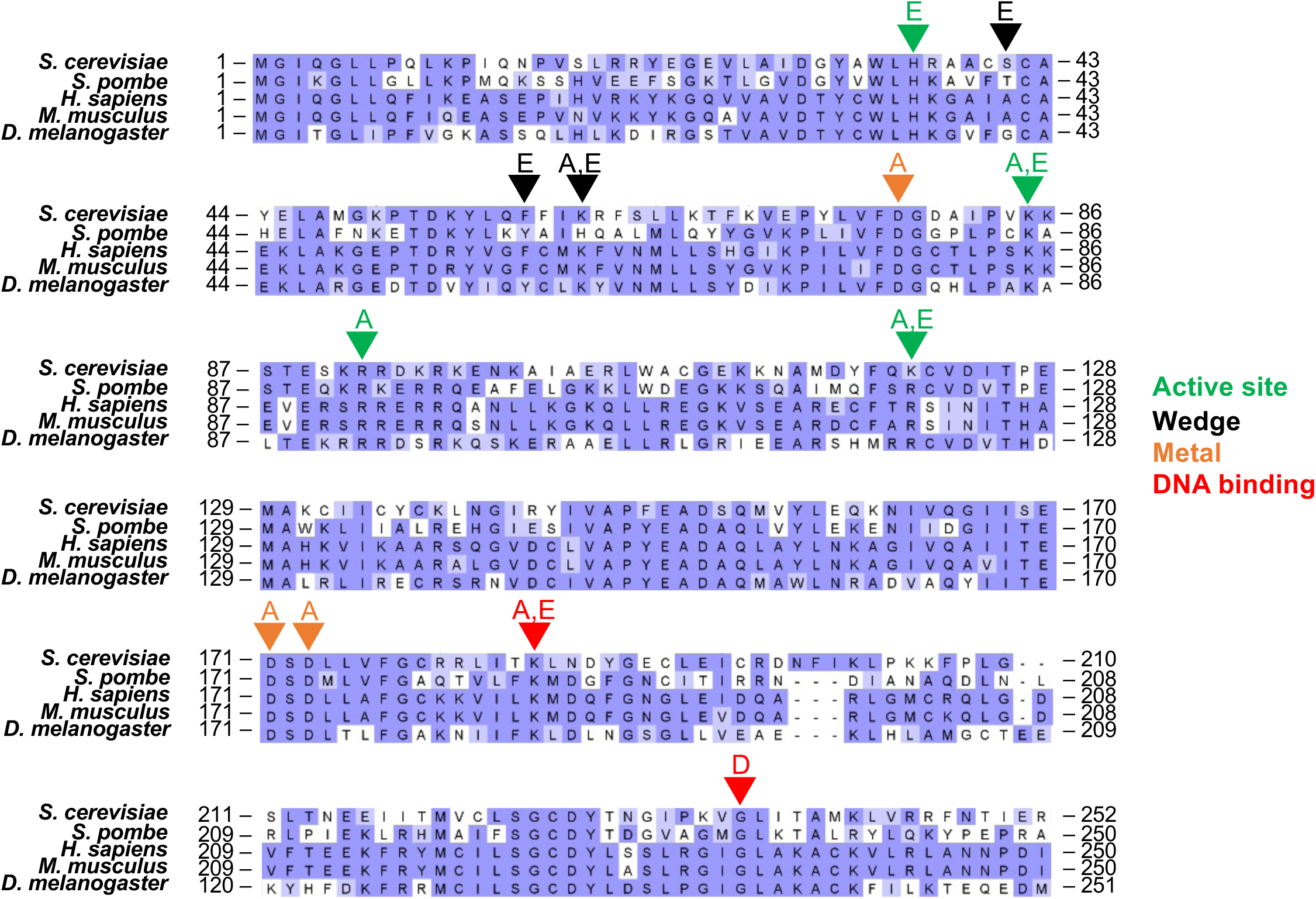
Alignment of Exo1 protein sequences from *S. cerevisiae* (accession # NP_014676)*, S. pombe* (NP_596050.1)*, H. sapiens (*NP_003677)*, M. musculus* (NP_036142) and *D. melanogaster* (NP_477145). Sequence alignment of Exo1 from different species. Triangles indicate mutations made in this study. See Materials and Methods for sequence alignment details.

**Figure S2.**
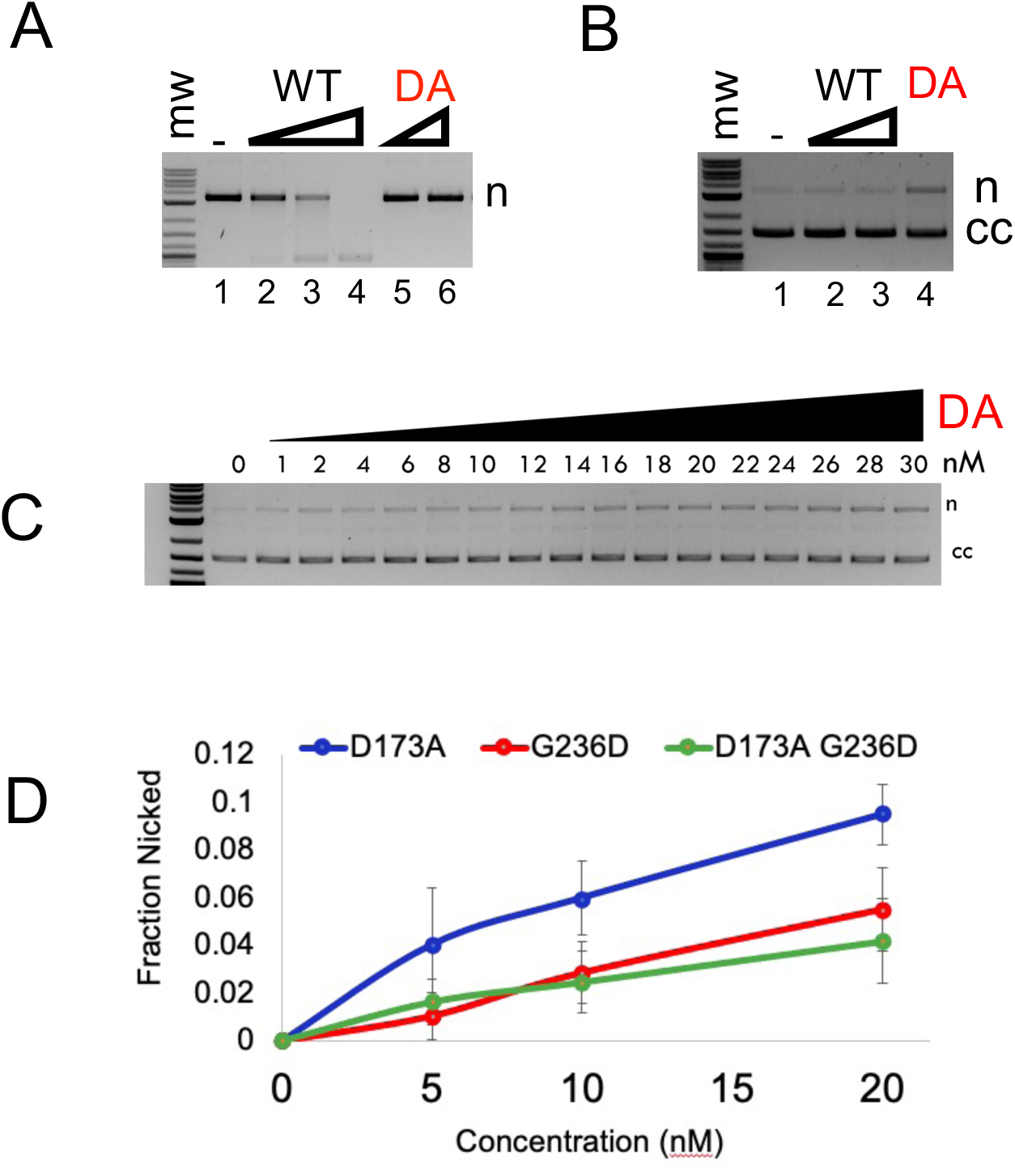
Nuclease activity of Exo1 on plasmid substrates. A. Nuclease activity of Exo1 (WT) and exo1-D173A (DA; Materials and Methods) on a 2.7 kb pUC18 plasmid with four pre-existing nicks. DNA products were resolved by native agarose gel. Exo1 is present at 6 nM, 12 nM, and 24 nM in lanes 2-4, and exo1-D173A is present at 20 and 40 nM in lanes 5-6. B. Exo1 does not show nuclease activity on supercoiled (cc) 2.7 kb pUC18 plasmid. Exo1 is present at 1 nM and 10 nM in lanes 2 and 3, respectively, and exo1-D173A is present at 20 nM in lane 4. B. Titration of exo1-D173A endonuclease activity on supercoiled (cc) pBR322 substrate. D. Titration of exo1-D173A, exo1-G236D and exo1-D173A,G236D endonuclease activity on a supercoiled pBR322 substrate.

**Figure S3.**
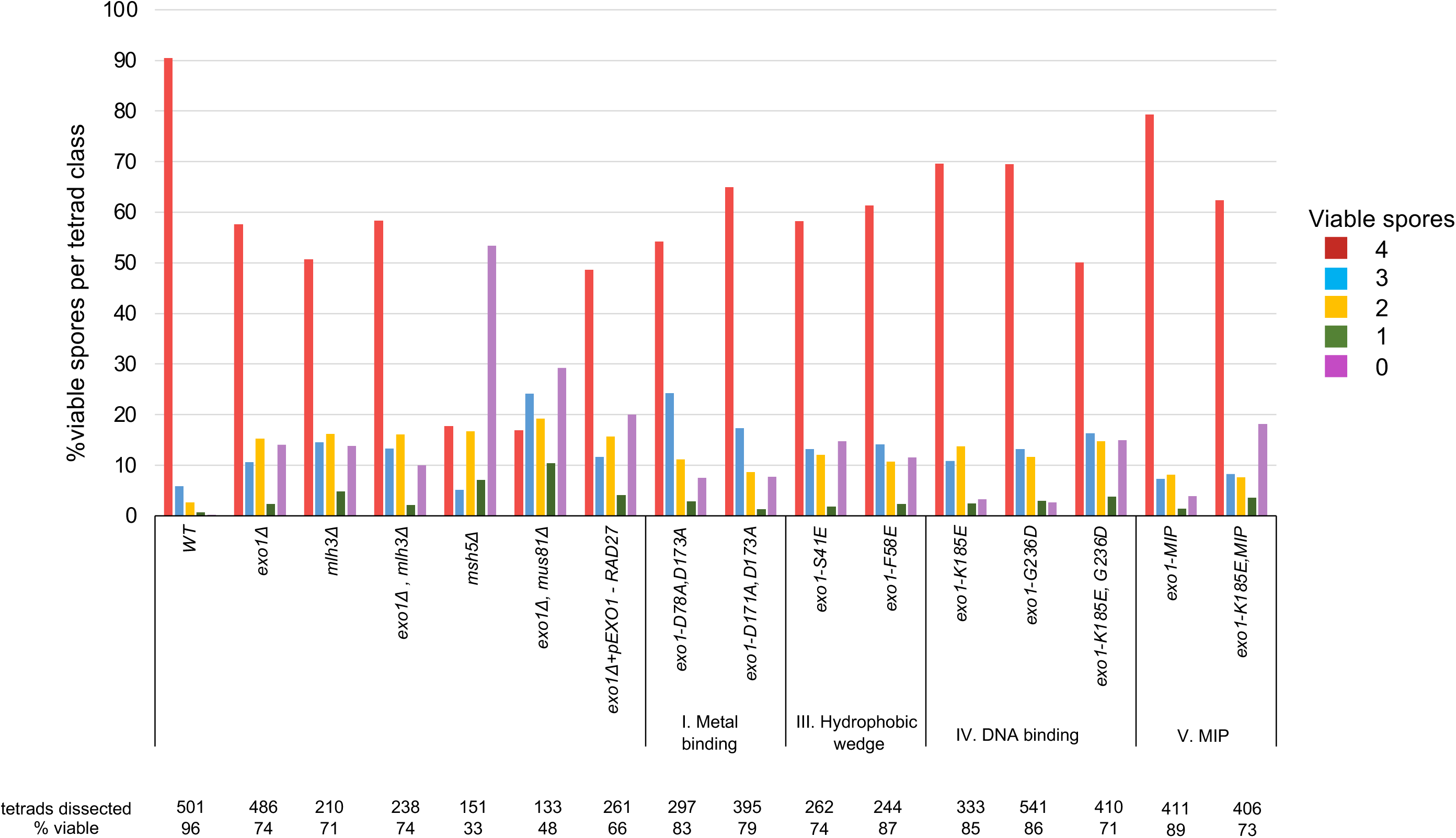
Spore viability profile of *wild-type* and the indicated *exo1* strains in the EAY1108/EAY1112 strain background. The percent of tetrads with 4, 3, 2, 1, and 0 viable spores are shown from the dissections presented in Figure 2 as well as the total number of tetrads dissected and the overall spore viability.

**Figure S4.**
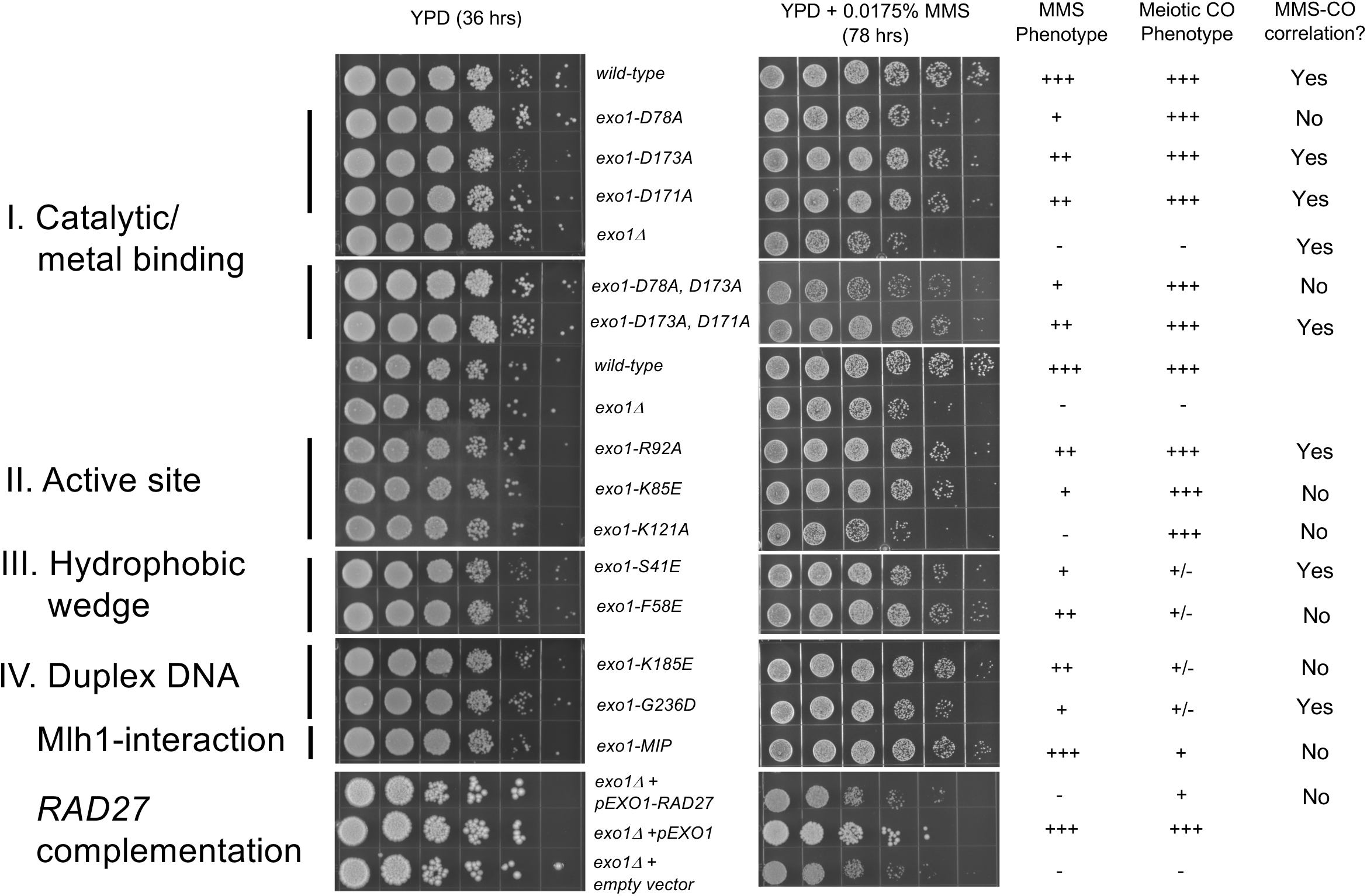
Sensitivity of *exo1* mutants to the DNA damaging agent MMS. *Wild-type* and the indicated *exo1* mutants (Figure 2A) were spotted in 10-fold serial dilutions onto YPD and YPD media containing 0.04% MMS (Materials and Methods). Plates were photographed after a 2-day incubation at 30°C. In the bottom most panel an *exo1Δ* strain (EAY4778) was transformed with an *ARS-CEN* vector containing no insert (pRS416), *EXO1* (pEAA715) or *RAD27* expressed from the *EXO1* promoter (*pEXO1-RAD27,* pEAA720).

**Figure S5.**
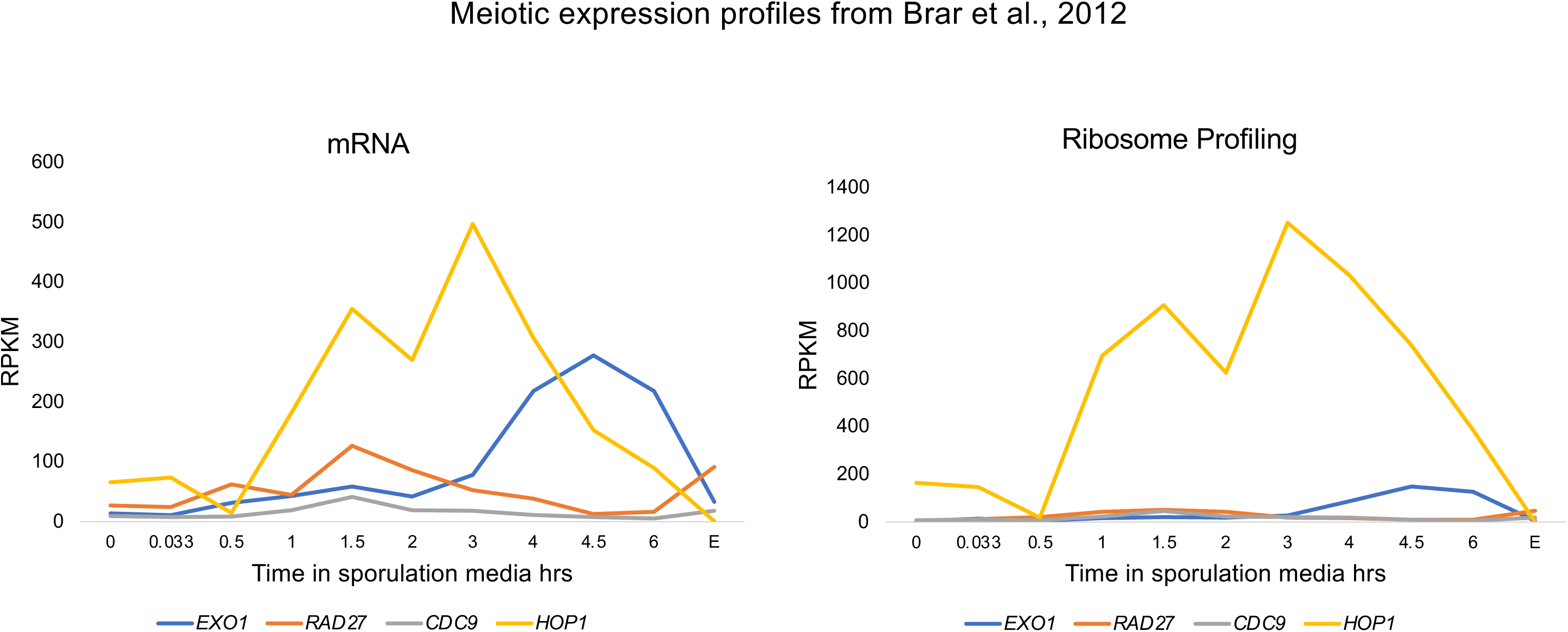
mRNA seq and ribosome profiling of *EXO1, RAD27, CDC9* and *HOP1* expression in SK1 meiosis. Data obtained from Brar et al. (2012). RPKM= Reads per kilobase of coding sequence per million mapped reads.

**Figure S6.**
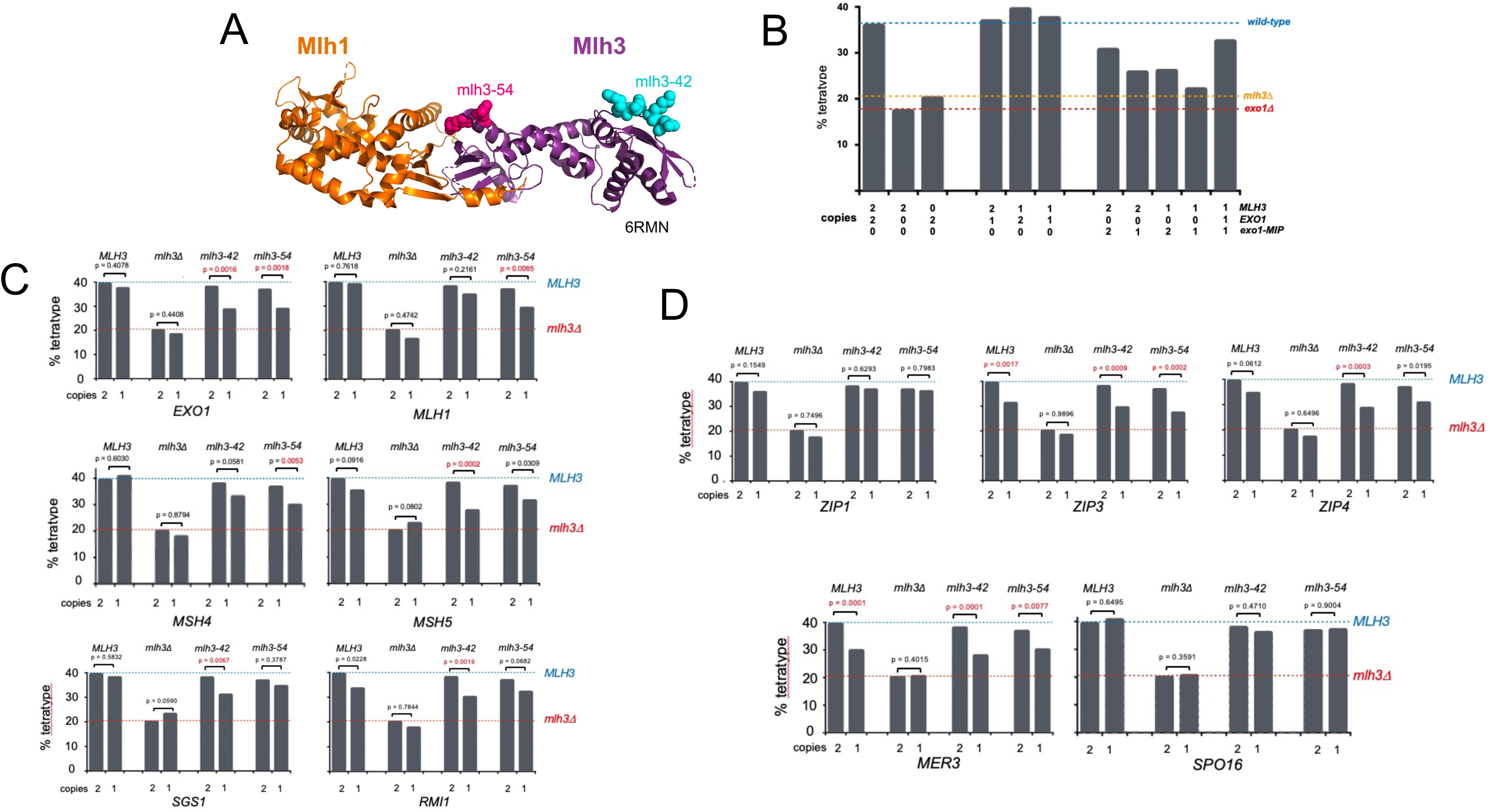
Haploinsufficiency analysis shows genetic interactions between *MLH3* and *MLH1, EXO1, MSH4, MSH5, SGS1*, and *RMI1*, but not between *MLH3* and *ZIP1, ZIP3, ZIP4, SPO16,* and *MER3*. A. The *mlh3-42* and *mlh3-54* mutations analyzed in haploinsufficiency analysis map onto the C-terminal domain of *MLH3.* Each allele confers defects in Mlh1-Mlh3 interactions and Mlh3-dependent DNA mismatch repair, but do not confer strong defects in meiotic crossing over (Al-Sweel et al., 2017). B. Strains containing one or two copies of *EXO1, MLH3, or the exo1-MIP* mutations were analyzed for crossing over in the 20 cM *CEN8* to *THR1* interval using a spore-autonomous fluorescence assay (Thacker et al., 2011). C. A haploinsufficiency screen identified *EXO1, MLH1, MSH4*, *MSH5, SGS1,* and *RMI1* interactions with *MLH3*. Strains containing one or two copies of *EXO1, MLH1, MSH4*, *MSH5*, *SGS1* and *RMI1* were analyzed for crossing over in *wild-type*, *mlh3Δ* and *mlh3-42* and *mlh3-54* strains (Materials and Methods). D. Haploinsufficiency of *ZIP1, ZIP3, ZIP4,* and *MER3* conferred decreases in crossover frequencies that were not *mlh3* alleles-specific, and haploinsufficiency of *SPO16* did not affect CO frequency. Crossing over was also measured in the 20 cM *CEN8* to *THR1* interval. Significance was assessed by *χ*^2^ test between haplosufficient and haploinsufficient conditions. To minimize α inflation due to multiple comparisons, we applied a Benjamini-Hochberg correction at a 5% false discovery rate.

**Table S1A.** Spore Autonomous Meiotic Crossover Analysis of *exo1* mutants. Homozygous mutations were made by crossing two independently constructed strains with the *exo1* variants in the SKY3576 (containing cyan fluorescent protein; Table S5) and SKY3575 (containing red fluorescent protein) backgrounds. Heterozygous mutations were made by crossing two independently constructed strains with *exo1* variants in the SKY3576 and EAY4151 (*exo1Δ*) backgrounds. Diploid strains were induced for meiosis and % tetratype in the *CEN8-THR1* interval was measured, by determining the total tetratypes/sum of tetratypes and parental ditypes). At least 500 tetrads were counted for each allele, and unless indicated (*one transformant analyzed), at least two transformants were analyzed for each background. Significance was assessed by Fisher’s exact test between mutant and *wild-type EXO1 and exo1Δ* tetratype values. To minimize α inflation due to multiple comparisons, we applied a Benjamini-Hochberg correction at a 5% false discovery rate. +, indistinguishable from *wild-type*; -, indistinguishable from *exo1Δ;* INT, distinguishable from both *wild-type* and *exo1Δ*.

| Allele | %Tetratype | Tetrads Counted | Phenotype |
| --- | --- | --- | --- |
| *EXO1/EXO1* | 39.0 | 1071 | + |
| *EXO1/exo1Δ* | 37.9 | 1071 | + |
| *exo1Δ/exo1Δ* | 20.0 | 1054 | - |
| ***Metal Binding (Group 1)*** |  |  |  |
| *exo1-D171A/exo1Δ* | 38.9 | 517 | + |
| *exo1-D171A,D173A/exo1Δ* | 39.1 | 511 | + |
| *exo1-D78A/exo1-D78A* | 39.7 | 531 | + |
| *exo1-D173A/exo1-D173A* | 37.4 | 519 | + |
| *exo1-D78A,D173A/ exo1-D78A,D173A* | 36.4 | 544 | + |
| ***Active Site DNA Interactors (Group II)*** |  |  |  |
| *exo1-H36E/exo1Δ* | 35.4 | 506 | + |
| *exo1-K85A/exo1Δ* | 34.8 | 526 | + |
| *exo1-K85E/exo1Δ* | 34.5 | 533 | + |
| *exo1-R92A/exo1Δ* | 34.8 | 506 | + |
| *exo1-R121A/exo1Δ* | 37.7 | 605 | + |
| *exo1-R121E/exo1Δ* | 34.5 | 765 | + |
| ***Hydrophobic Wedge (Group III)*** |  |  |  |
| *exo1-S41E/exo1Δ* | 28.4 | 506 | INT |
| *exo1-F58E/exo1Δ* | 27.8 | 507 | INT |
| *exo1-K61A/exo1Δ* | 35.1 | 525 | + |
| *exo1-K61E/exo1Δ* | 35.1 | 507 | + |
| *exo1-S41E,F58E/exo1Δ* | 24.6 | 504 | INT |
| *exo1-S41E,K61E/exo1Δ* | 24.5 | 506 | - |
| ***Duplex DNA Interactors (Group IV)*** |  |  |  |
| *exo1-K185A/exo1Δ* | 35.4 | 720 | + |
| *exo1-K185E/exo1Δ* | 24.5 | 649 | INT |
| *exo1-G236D/exo1-G236D* | 29.9 | 521 | INT |
| *exo1-G236D/exo1Δ* | 29.1 | 515 | INT |
| *exo1-G236D,K185E/exo1Δ* | 24.2 | 508 | - |
| ***Mlh1-Interacting (MIP, Group V)*** |  |  |  |
| *exo1- F447A,F448A/exo1-F447A,F448A* | 33.3 | 547 | INT |
| *exo1- F447A,F448A/exo1Δ* | 26.2 | 519 | INT |
| ***Double and Triple Mutants*** |  |  |  |
| *exo1-D171A,G236D/exo1Δ* | 31.1 | 552 | INT |
| *exo1-D173A,G236D/exo1Δ* | 32.7 | 618 | + |
| *exo1-D173A,G236D/ exo1-D173A,G236D* | 35.7 | 532 | + |
| *exo1-D173A,K185E,G236D/exo1Δ* | 22.4 | 553 | - |
| *exo1-G236D,F447A,F448A/exo1Δ* | 25.1 | 617 | INT |
| *exo1-K185E,F447A,F448A/exo1Δ* | 24.8 | 572 | INT |
| *exo1-D173A,G236D,F447A,F448A/exo1Δ* | 26.6 | 500 | INT |
| *exo1-R92A,R121A,K185A/exo1Δ* | 24.3 | 535 | - |

**Table S1B.** Spore autonomous assay: *pEXO1-RAD27* complementation of *exo1Δ* and *mlh3Δ* strains. Diploids of the indicated genotype that contain markers to measure crossing over in the *CEN8-THR1* interval (Table S5) were transformed with the indicated plasmids (pEAA715-*EXO1, URA3, CEN6-ARSH4;* pRS416-*URA3,CEN6-ARSH4;* pEAA722-*RAD27*, *URA3, CEN6-ARSH4;* pEAA720-*pEXO1-RAD27, URA3, CEN6-ARSH4;* pEAA724-*pEXO1-rad27-D179A, URA3, CEN6-ARSH4;* pEAA727*-rad27-A45E, URA3, CEN6-ARSH4;* pEAA728*-rad27-R101A, URA3, CEN6-ARSH4;* pEAA729*-rad27-R105A, URA3, CEN6-ARSH4;* pEAA73*0-rad27-K130A, URA3, CEN6-ARSH4;* pEAA731*-rad27-H191E, URA3, CEN6-ARSH4*) and selected for plasmid retention. The resulting strains were induced for meiosis and % tetratype (single crossovers) in the *CEN8-THR1* interval was measured, by determining the total tetratypes/sum of tetratypes and parental ditypes. At least 500 tetrads were counted for each allele/plasmid combination, and at least two transformants were analyzed for each condition. Significance (presented in Figure 4A, C) was assessed by Fisher’s Exact Test between *exo1Δ* strains containing pRS416 (empty vector) and test conditions with the indicated plasmids. To minimize α inflation due to multiple comparisons, we applied a Benjamini-Hochberg correction at a 5% false discovery rate. The significance of % tetratype in *exo1-K185E* and *exo1-F447A,F448A (MIP)* strains containing pRS416 (empty vector) and pEAA720 (*pEXO1-RAD27*) was determined using Fisher’s exact test. N/A, not applicable.

| Genotype | Plasmid | %Tetratype | Tetrads Counted | Phenotype |
| --- | --- | --- | --- | --- |
| *exo1Δ/exo1Δ* | *EXO1* | 34.1 | 557 | + |
| *exo1Δ/exo1Δ* | *empty vector* | 21.5 | 512 | - |
| *exo1Δ/exo1Δ* | *RAD27* | 22.6 | 1032 | - |
| *exo1Δ/exo1Δ* | *pEXO1-RAD27* | 29.9 | 521 | + |
| *exo1Δ/exo1Δ* | *pEXO1-rad27-D179A* | 28.8 | 510 | + |
| *exo1Δ/exo1Δ* | *pEXO1-rad27-A45E* | 22.4 | 511 | - |
| *exo1Δ/exo1Δ* | *pEXO1-rad27-R101A* | 29.7 | 542 | + |
| *exo1Δ/exo1Δ* | *pEXO1-rad27-R105A* | 28.7 | 521 | + |
| *exo1Δ/exo1Δ* | *pEXO1-rad27-K130A* | 28.9 | 505 | + |
| *exo1Δ/exo1Δ* | *pEXO1-rad27-H191E* | 24.0 | 530 | - |
| *mlh3Δ/mlh3Δ* | *MLH3* | 35.6 | 508 | + |
| *mlh3Δ/mlh3Δ* | *empty vector* | 22.5 | 528 | - |
| *mlh3Δ/mlh3Δ* | *pEXO1-RAD27* | 21.5 | 512 | - |
| *mlh3Δ/mlh3Δ* | *pEXO1-rad27-D179A* | 19.9 | 513 | - |
| *exo1-K185E/exo1Δ* | *empty vector* | 25.4 | 1538 | N/A |
| *exo1-K185E/exo1Δ* | *pEXO1-RAD27* | 29.0 | 1541 | N/A |
| *exo1-F447A,F448A/exo1Δ* | *empty vector* | 30.1 | 512 | N/A |
| *exo1-F447A,F448A/exo1Δ* | *pEXO1-RAD27* | 29.7 | 526 | N/A |

**Table S1C.**
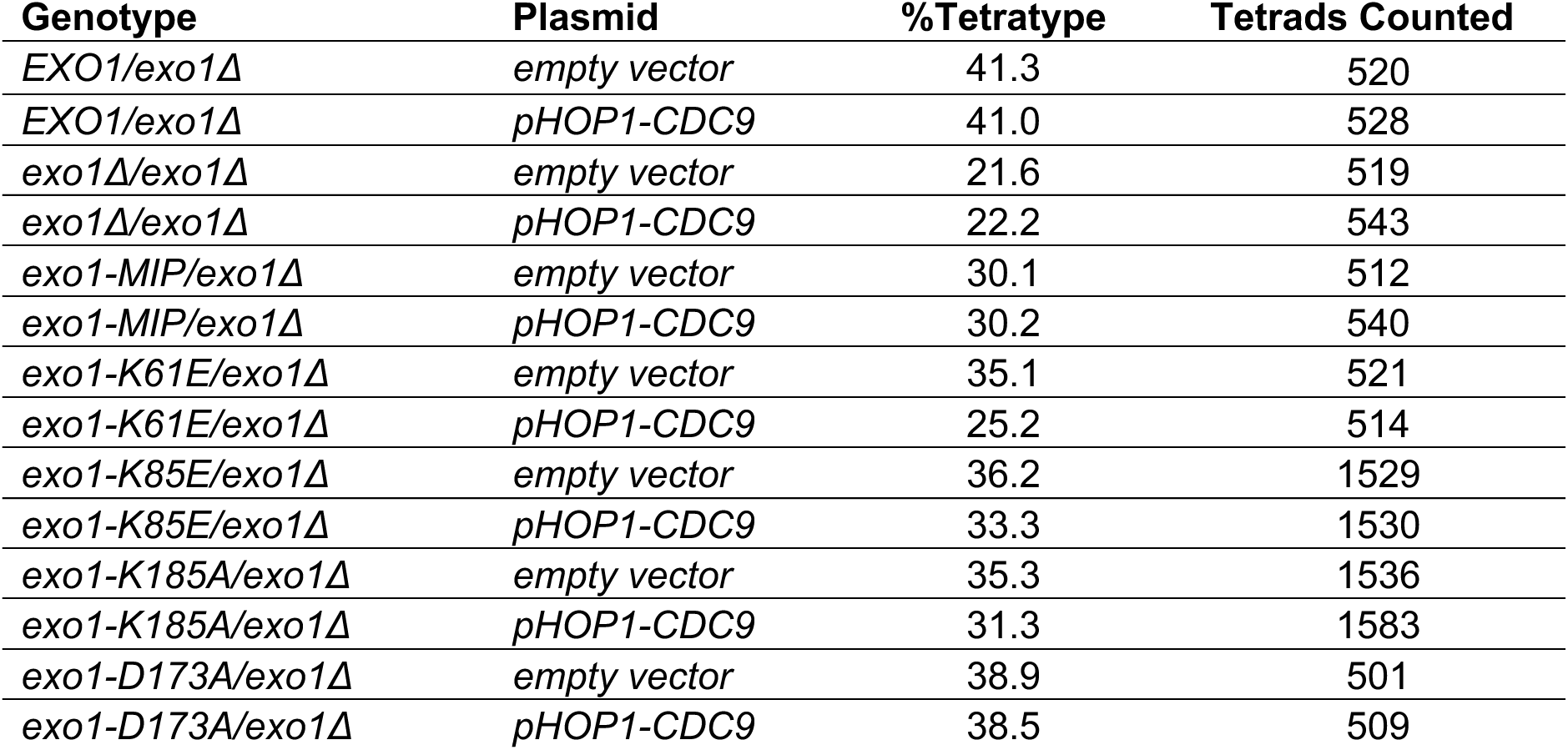
Effect of *pHOP1-CDC9* expression on meiotic crossing over in *exo1* strains. Diploids of the annotated genotype were transformed with the indicated plasmid (pRS426-*URA3, 2μ;* pEAM329*-pHOP1-CDC9, URA3, 2μ*) and selected for diploidy and plasmid retention. Diploid strains were induced for meiosis and % Tetratype in the *CEN8-THR1* interval was measured by determining the total tetratypes/sum of tetratypes and parental ditypes. At least 500 tetrads were counted for each allele/plasmid combination, and at least two transformants were analyzed for each condition. Significance was assessed by Fisher’s exact test between pRS426 value and pEAM329 value and is shown in Figure 4D.

**Table S2.** Genetic map distances (cM) and the distribution of parental and recombinant progeny for the EAY1108/EAY1112 strain background in *WT, mlh3Δ, msh5Δ,* and *exo1* strains on Chromosome XV. Mutants are isogenic derivatives of EAY1108/EAY1112. Genetic intervals correspond to the genetic distance calculated from tetrads +/- one standard error. Standard error was calculated using the Stahl Laboratory Online Tools website (https://elizabethhousworth.com/StahlLabOnlineTools/). For single spore analysis, data are shown as 95% confidence intervals around the recombination frequency. For tetrad analysis the centimorgan (cM) map distance was calculated using the formula of Perkins (1949): 50{TT+(6NPD)}/(PD+TT+NPD). To compare to the tetrad data, recombination frequencies obtained from single spores (Parental/(Parental+Recombinant)) were multiplied by 100 to yield genetic map distances (cM).

| Table S2. Genetic map distances (cM) and the distribution of parental and recombinant progeny for the EAY1108/EAY1112 strain background in *WT*, *mlh3Δ*, *msh5Δ*, and *exo1* strains on Chromosome XV. |  |  |  |  |  |  |  |  |  |
| --- | --- | --- | --- | --- | --- | --- | --- | --- | --- |
| Relevant genotype | Tetrads |  |  |  |  | Single spores |  |  |  |
|  | Number analyzed | cM | PD | TT | NPD | Number analyzed | cM (95% CI) | Parental | Recombinant |
| *URA3-LEU2*: |  |  |  |  |  |  |  |  |  |
| *wild-type* | 501 | 22.4+/-1.5 | 292 | 206 | 3 | 2285 | 22.2 (19.8-23.2) | 1794 | 491 |
| *exo1Δ* | 486 | 11.2+/-1.4 | 392 | 91 | 3 | 2510 | 9.6 (8.5-10.8) | 2267 | 241 |
| *exo1-K185E* | 333 | 17.1+/-2.5 | 227 | 96 | 12 | 1549 | 18.1 (16.2-20.1) | 1269 | 280 |
| *exo1-G236D* | 541 | 15.8+/-1.6 | 388 | 129 | 6 | 2676 | 14.2 (12.9-15.6) | 2296 | 380 |
| *exo1-K185E G236D* | 410 | 16.3+/-1.5 | 286 | 122 | 2 | 2409 | 15.5 (14.1-17.0) | 2033 | 374 |
| *exo1-MIP* | 411 | 14.8+/-1.6 | 304 | 104 | 3 | 1915 | 12.8 (11.4-14.4) | 1669 | 246 |
| *exo1-K185E,MIP* | 406 | 14.9+/-1.6 | 300 | 103 | 3 | 4036 | 12.7 (11.3-14.3) | 1934 | 246 |
| *exo1-D78A,D173A* | 297 | 18.9+/-2.1 | 200 | 94 | 3 | 1760 | 16.2 (14.4-17.9) | 1471 | 284 |
| *exo1-D171A,D173A* | 395 | 18.4+/-1.2 | 250 | 145 | 0 | 2049 | 18.9 (17.2-20.6) | 1662 | 387 |
| *exo1-F58E* | 244 | 16.6+/-1.5 | 163 | 81 | 0 | 1112 | 19.4 (17.1-21.9) | 896 | 216 |
| *exo1-S41E* | 262 | 10.3+/-1.2 | 208 | 54 | 0 | 1362 | 11.1 (9.5-12.9) | 1211 | 151 |
| *mlh3Δ* | 210 | 13.6+/-2.7 | 168 | 39 | 3 | 1191 | 10.5 (8.8-12.4) | 1066 | 125 |
| *msh5Δ* | 151 | 10.9+/-1.7 | 118 | 33 | 0 | 1111 | 8.5 (6.9-10.2) | 1017 | 94 |
| *exo1Δ mus81Δ* | 133 | 2.8+/-0.9 | 128 | 6 | 0 | 1767 | 1.2 (0.7-1.8) | 1745 | 21 |
| *exo1Δ mlh3Δ* | 238 | 11+/-1.3 | 184 | 52 | 0 | 1221 | 10.4 (8.7-12.2) | 1092 | 127 |
| *exo1Δ + pRAD27 (EXO1 Promoter)* | 261 | 12.3+/-1.3 | 197 | 64 | 0 | 1433 | 12.6 (10.9-14.4) | 1253 | 180 |
| *exo1Δ + pEXO1* | 208 | 21.9+/-2.5 | 127 | 79 | 2 | 1072 | 19.4 (17.1-21.9) | 864 | 208 |
| *exo1Δ + pEmpty Vector* | 220 | 12.5+/-2.3 | 175 | 43 | 2 | 1149 | 10.2 (8.5-12.1) | 1032 | 117 |
| *LEU2-LYS2*: |  |  |  |  |  |  |  |  |  |
| *wild-type* | 501 | 28.7+/-1.5 | 233 | 264 | 4 | 2285 | 27.6 (25.7-29.4) | 1655 | 630 |
| *exo1Δ* | 486 | 11.1+/-1.1 | 377 | 108 | 1 | 2510 | 11.9 (10.6-13.2) | 2210 | 298 |
| *exo1-K185E* | 333 | 21.2+/-2.9 | 272 | 56 | 7 | 1549 | 9.8 (8.4-11.4) | 1397 | 152 |
| *exo1-G236D* | 541 | 17.3+/-1.6 | 372 | 145 | 6 | 2676 | 16.2 (14.8-17.7) | 2242 | 434 |
| *exo1-K185E G236D* | 410 | 15.1+/-1.5 | 296 | 112 | 2 | 2409 | 13.6 (12.2-15.0) | 2080 | 327 |
| *exo1-MIP* | 411 | 13.9+/-1.6 | 312 | 96 | 3 | 1915 | 12.5 (11.0-14.0) | 1676 | 239 |
| *exo1-K185E,MIP* | 406 | 16.3+/-1.2 | 274 | 132 | 0 | 1934 | 16.3 (14.7-18.0) | 1619 | 315 |
| *exo1-D78A,D173A* | 297 | 27.6+/-2.3 | 153 | 140 | 4 | 1760 | 24.7 (22.6-26.7) | 1322 | 433 |
| *exo1-D171A.D173A* | 395 | 22.7+/-1.2 | 216 | 179 | 0 | 2049 | 21.9 (20.1-23.7) | 1601 | 448 |
| *exo1-F58E* | 244 | 14.9+/-1.8 | 176 | 67 | 1 | 1112 | 15.5 (13.4-17.7) | 940 | 172 |
| *exo1-S41E* | 262 | 13.4+/-2.0 | 202 | 58 | 2 | 1362 | 13.4 (11.7-15.4) | 1179 | 183 |
| *mlh3Δ* | 210 | 14.3+/-2.0 | 155 | 54 | 1 | 1191 | 13.1 (11.2-15.1) | 1035 | 156 |
| *msh5Δ* | 151 | 9.6+/-1.7 | 122 | 29 | 0 | 1111 | 9.5 (7.9-11.4) | 1005 | 106 |
| *exo1Δ mus81Δ* | 133 | 3.4+/-1.1 | 126 | 9 | 0 | 1767 | 2.6 (1.9-3.4) | 1722 | 45 |
| *exo1Δ mlh3Δ* | 238 | 10.4+/-1.3 | 187 | 49 | 0 | 1221 | 11.5 (9.7-13.4) | 1079 | 140 |
| *exo1Δ + pRAD27*<br>(*EXO1 Promoter*) | 261 | 17.8+/-2.1 | 178 | 81 | 2 | 1433 | 15.2 (13.4-17.2) | 1215 | 218 |
| *exo1Δ + pEXO1* | 208 | 26.2+/-2.9 | 114 | 91 | 3 | 1072 | 22.7 (20.2-25.3) | 829 | 243 |
| *exo1Δ + pEmpty Vector* | 220 | 12.7+/-1.9 | 169 | 50 | 1 | 1149 | 12.5 (10.7-14.6) | 1005 | 144 |
| *LYS2-ADE2:* |  |  |  |  |  |  |  |  |  |
| *wild-type* | 501 | 14.8+/-1.2 | 358 | 142 | 1 | 2285 | 14.8 (13.4-16.3) | 1947 | 338 |
| *exo1Δ* | 486 | 7.3+/-1.1 | 425 | 59 | 2 | 2510 | 6.6 (5.7-7.7) | 2342 | 166 |
| *exo1-K185E* | 333 | 12.3+/-1.8 | 267 | 67 | 1 | 1549 | 10.3 (8.8-11.9) | 1390 | 159 |
| *exo1-G236D* | 541 | 11.8+/-1.4 | 420 | 99 | 4 | 2676 | 10.0 (8.9-11.2) | 2409 | 267 |
| *exo1-K185E G236D* | 410 | 6.6+/-1.3 | 366 | 42 | 2 | 2409 | 5.9 (4.9-6.9) | 2266 | 141 |
| *MIP* | 411 | 7.8+/-0.9 | 347 | 64 | 0 | 1915 | 7.4 (6.3-8.7) | 1773 | 142 |
| *exo1-K185E, MIP* | 406 | 8.3+/-0.9 | 339 | 67 | 0 | 1934 | 7.7 (6.5-9.0) | 1785 | 149 |
| *exo1-D78A,D173A* | 297 | 12.1+/-1.5 | 230 | 66 | 1 | 1760 | 11.3 (9.8-12.8) | 1557 | 198 |
| *exo1-D171A,D173A* | 395 | 8.9+/-1.0 | 325 | 70 | 0 | 2049 | 10.2 (8.9-11.5) | 1841 | 208 |
| *exo1-F58E* | 244 | 11.7+/-1.3 | 148 | 64 | 0 | 1112 | 14.7 (12.7-17.0) | 948 | 164 |
| *exo1-S41E* | 262 | 8.6+/-1.2 | 217 | 45 | 0 | 1362 | 8.1 (6.7-9.7) | 1251 | 111 |
| *mlh3Δ* | 210 | 5.5+/-1.1 | 187 | 23 | 0 | 1191 | 5.1 (3.9-6.5) | 1130 | 61 |
| *msh5Δ* | 151 | 4.6+/-1.3 | 137 | 14 | 0 | 1111 | 3.7 (2.7-5.0) | 1070 | 41 |
| *exo1Δ mus81Δ* | 133 | 0+/-0 | 133 | 0 | 0 | 1763 | 1.4 (0.9-2.0) | 1738 | 24 |
| *exo1Δ mlh3Δ* | 238 | 6.6+/-1.6 | 210 | 25 | 1 | 1221 | 6.1 (4.8-7.5) | 1145 | 74 |
| *exo1Δ + pRAD27*<br>(*EXO1 Promoter*) | 261 | 8.0+/-1.1 | 219 | 42 | 0 | 1433 | 8.6 (7.2-10.2) | 1310 | 123 |
| *exo1Δ + pEXO1* | 208 | 15.8+/-2.5 | 152 | 54 | 2 | 1072 | 14.2 (12.1-16.4) | 920 | 152 |
| *exo1Δ + pEmpty Vector* | 220 | 5.4+/-1.0 | 196 | 24 | 0 | 1149 | 6.5 (5.2-8.1) | 1074 | 75 |
| *ADE2-HIS4:* |  |  |  |  |  |  |  |  |  |
| *wild-type* | 501 | 39+/-2.1 | 170 | 319 | 12 | 2285 | 35.1 (33.1-37.1) | 1483 | 802 |
| *exo1Δ* | 486 | 24.8+/-2.1 | 295 | 181 | 10 | 2510 | 20.9 (19.3-22.5) | 1985 | 523 |
| *exo1-K185E* | 333 | 22.5+/-2.3 | 203 | 131 | 1 | 1549 | 20.7 (18.7-22.8) | 1229 | 320 |
| *exo1-G236D* | 541 | 23.8+/-2.2 | 339 | 171 | 13 | 2676 | 20.0 (18.6-21.6) | 2139 | 537 |
| *exo1-K185E G236D* | 410 | 23.3+/-2.1 | 254 | 149 | 7 | 2409 | 20.1 (18.5-21.7) | 1923 | 484 |
| *exo1-MIP* | 411 | 23.2+/-1.7 | 235 | 173 | 3 | 1915 | 21.6 (19.8-23.5) | 1501 | 414 |
| *exo1-K185E,MIP* | 406 | 23.4+/-2.1 | 246 | 154 | 6 | 1934 | 19.6 (17.9-21.5) | 1554 | 380 |
| *exo1-D78A,D173A* | 297 | 41.6+/-3.2 | 105 | 181 | 11 | 1760 | 34.4 (32.0-36.5) | 1152 | 603 |
| *exo1-D171A D173A* | 395 | 35.2+/-2.7 | 182 | 200 | 13 | 2049 | 29.0 (27.1-31.1) | 1454 | 595 |
| *exo1-F58E* | 244 | 26.6+/-2.5 | 92 | 120 | 0 | 1112 | 28.4 (25.8-31.2) | 796 | 316 |
| *exo1-S41E* | 262 | 26.3+/-3.2 | 159 | 96 | 7 | 1362 | 23.0 (20.8-25.3) | 1049 | 313 |
| *mlh3Δ* | 210 | 22.9+/-3.1 | 134 | 72 | 4 | 1191 | 18.9 (16.7-21.2) | 966 | 225 |
| *msh5Δ* | 151 | 16.9+/-3.4 | 110 | 39 | 2 | 1111 | 13.0 (11.0-15.1) | 967 | 144 |
| *exo1Δ mus81Δ* | 133 | 2.2+/-0.9 | 126 | 6 | 0 | 1763 | 1.2 (0.7-1.8) | 1710 | 21 |
| *exo1Δ mlh3Δ* | 238 | 25.8+/-3.1 | 139 | 92 | 5 | 1221 | 21.3 (19.0-23.7) | 959 | 260 |
| *exo1Δ + pRAD27*<br>*(EXO1 Promoter)* | 261 | 33.7+/-4.0 | 150 | 98 | 13 | 1433 | 23.0 (20.9-25.3) | 1103 | 330 |
| *exo1Δ + pEXO1* | 208 | 36.1+/-3.3 | 83 | 120 | 5 | 1072 | 32.1 (29.3-35.0) | 728 | 344 |
| *exo1Δ + pEmpty Vector* | 220 | 23.2+/-2.4 | 128 | 90 | 2 | 1149 | 21.2 (18.9-23.7) | 905 | 244 |

**Table S3A.** Interference measurements on Chromosome XV. The Malkova ratio and coefficient of coincidence (COC, ratio of double crossovers observed/double crossovers expected) were performed for the indicated genotypes in the EAY1108/EAY1112 strain background (Materials and Methods, strains listed in Table S5). These methods were performed for intervals I (*URA3-LEU2-LYS2)*, II (*LEU2-LYS2-ADE2)*, and III *(LYS2-ADE2-HIS3)*. 0 = Absolute Interference; 1= No interference. Significance of differences in tetrad distribution was assessed using a G test. Differences in distribution with p<0.05 were considered to be significant evidence of interference. Intervals with ratios significantly above 1 were observed and denoted with * to indicate potential negative interference. Detailed analysis of the Malkova ratio calculation is presented in Table S3B.

| Intervals | I | II | III | # tetrads | Interference |
| --- | --- | --- | --- | --- | --- |
| *wild-type* |  |  |  |  |  |
| Malkova | 0.48 | 0.43 | 0.90 | 501 | Intervals I, II |
| C.O.C | 0.66 | 0.52 | 0.92 |  |  |
| *exo1Δ* |  |  |  |  |  |
| Malkova | 1.28 | 0.84 | 1.62 | 486 | No intervals |
| C.O.C | 1.10 | 0.78 | 1.10 |  |  |
| *exo1-D171A,D173A* |  |  |  |  |  |
| Malkova | 0.37 | 0.53 | 0.40 | 395 | All intervals |
| C.O.C | 0.50 | 0.63 | 0.58 |  |  |
| *exo1-D78A, D173A* |  |  |  |  |  |
| Malkova | 0.54 | 0.17 | 0.65 | 297 | All intervals |
| C.O.C | 0.59 | 0.28 | 0.9 |  |  |
| *exo1-S41E* |  |  |  |  |  |
| Malkova | 0.46 | 0.15 | 0.56 | 262 | Intervals I, II |
| C.O.C | 0.56 | 0.19 | 0.73 |  |  |
| *exo1-F58E* |  |  |  |  |  |
| Malkova | 0.60 | 0.30 | 0.78 | 244 | Interval II |
| C.O.C | 0.79 | 0.21 | 0.99 |  |  |
| *exo1-G236D* |  |  |  |  |  |
| Malkova | 0.25 | 0.44 | 1.03 | 541 | Intervals I, II |
| C.O.C | 0.33 | 0.53 | 0.93 |  |  |
| *exo1-K185E* |  |  |  |  |  |
| Malkova | 0.47 | 0.40 | 1.70* | 333 | Intervals I, II |
| C.O.C. | 0.60 | 0.49 | 1.2 |  |  |
| *exo1-G236D,K185E* |  |  |  |  |  |
| Malkova | 0.50 | 0.82 | 1.19 | 410 | Intervals I |
| C.O.C | 0.64 | 0.98 | 0.89 |  |  |
| *exo1-MIP* |  |  |  |  |  |
| Malkova | 0.92 | 0.48 | 0.59 | 411 | Interval II |
| C.O.C. | 0.66 | 0.58 | 0.69 |  |  |
| *exo1Δ + pEXO1-RAD27* |  |  |  |  |  |
| Malkova | 1.41* | 0.90 | 0.81 | 261 | No intervals |
| C.O.C | 1.3 | 0.97 | 0.84 |  |  |
| *mlh3Δ* |  |  |  |  |  |
| Malkova | 0.46 | 0.58 | 1.08 | 210 | Interval I |
| C.O.C | 0.63 | 0.66 | 0.96 |  |  |
| *msh5Δ* |  |  |  |  |  |
| Malkova | 1.29 | 0.97 | 2.16* | 151 | No intervals |
| C.O.C | 1.1 | 1.12 | 1.84 |  |  |

**Table S3B.**
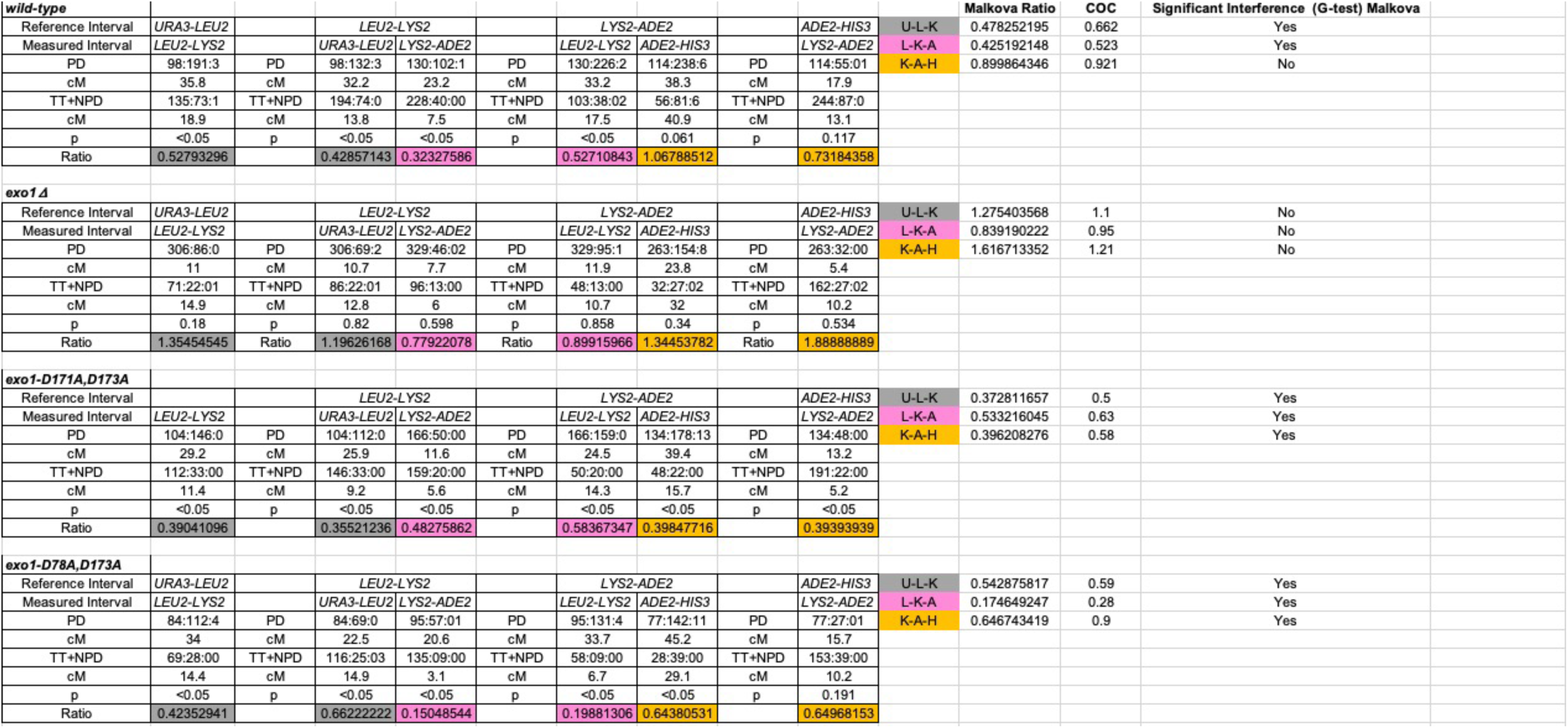
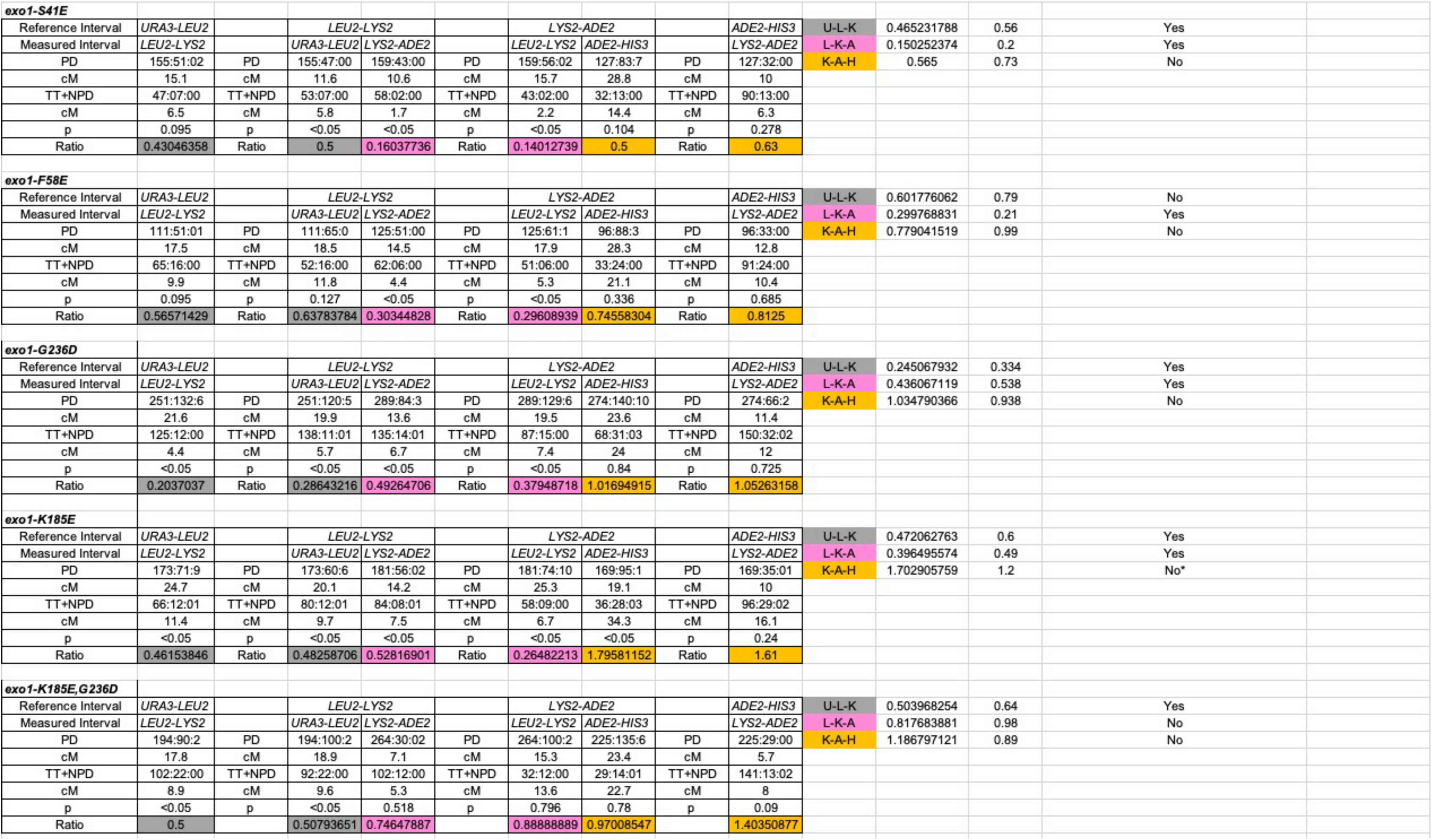
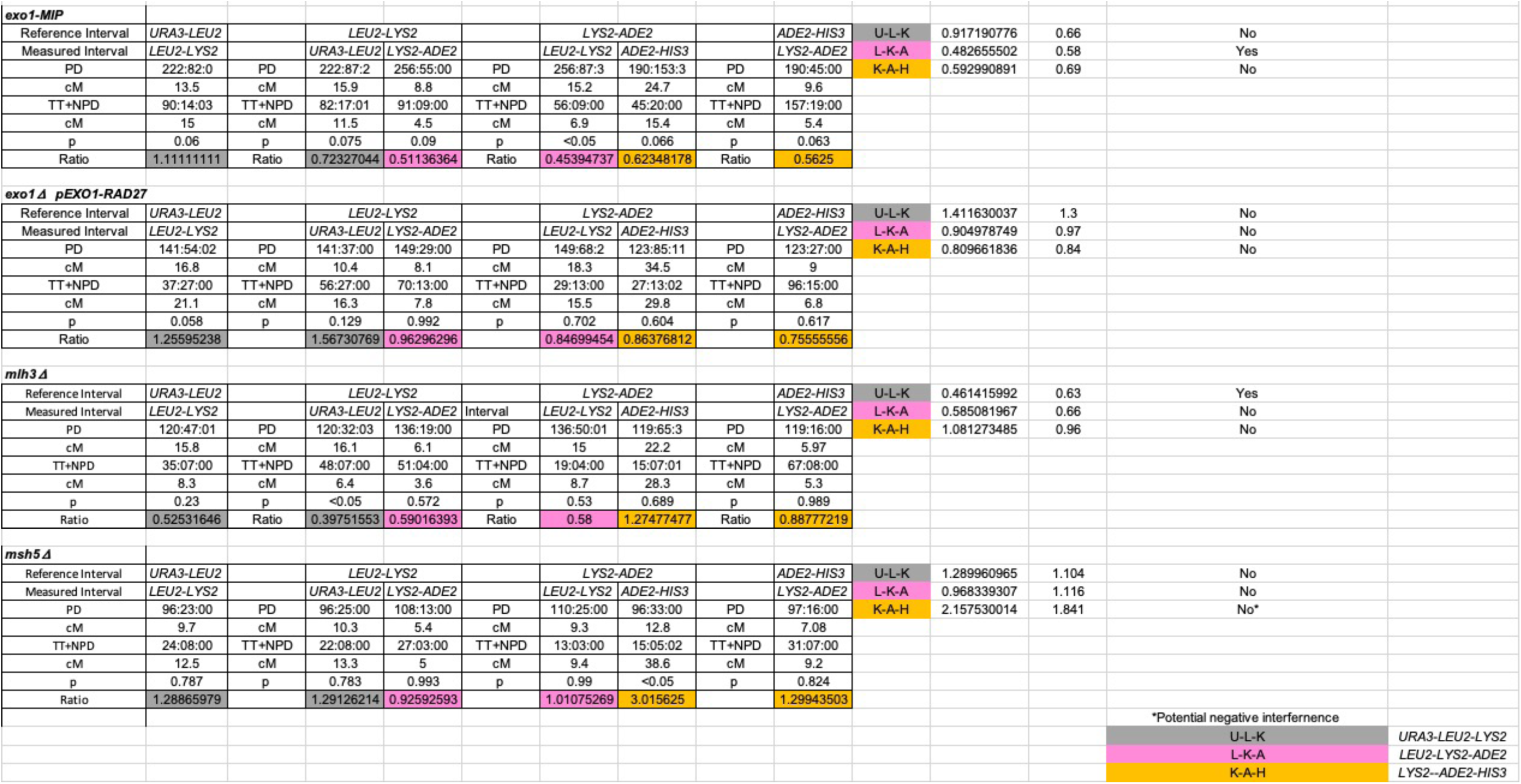
Detailed calculations of Malkova ratios presented in Figure 5 and Table S3A. Crossover interference was analyzed using the Malkova method (Malkova et al., 2004; Martini et al., 2006) for chromosome XV. For each genetic interval, tetrads were divided based on the presence or absence of a recombination event in a reference interval. For each reference interval, the map distance was measured in the adjacent intervals, thus obtaining two map distances for each interval. The significance of differences in tetrad distribution was assessed using a G test. Differences in distribution, with p<0.05, were considered to be evidence of interference. The data are presented as the average ratio of the two map distances in each neighboring interval, with a smaller ratio indicating stronger interference. An interval was considered to have a “loss of positive interference” phenotype when both adjacent intervals displayed no detectable positive interference. Ratios significantly greater than 1 are indicated with * to denote potential negative interference. TT, tetratype; NPD. nonparental ditype; PD, parental ditype.

**Table S4.**
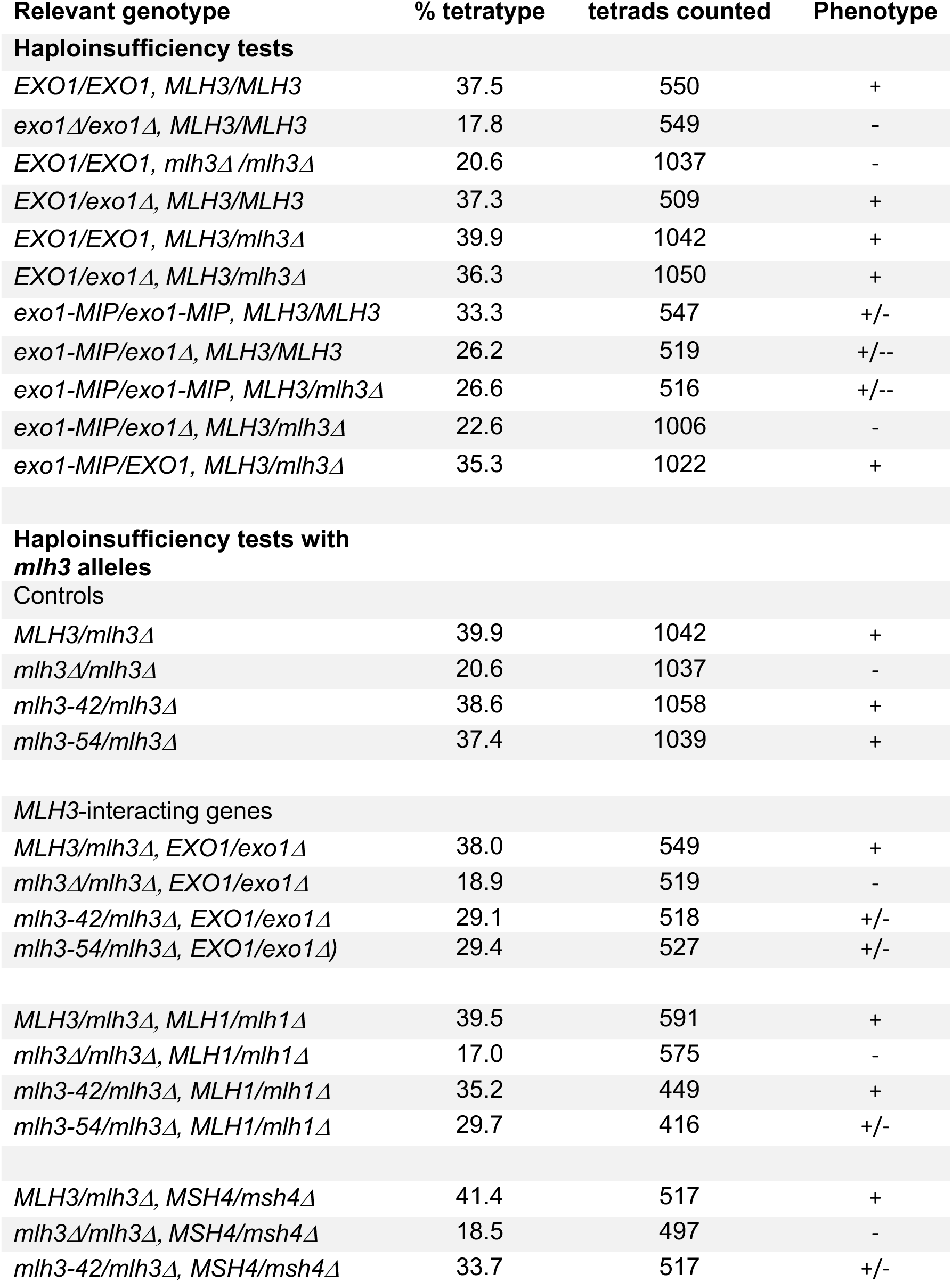
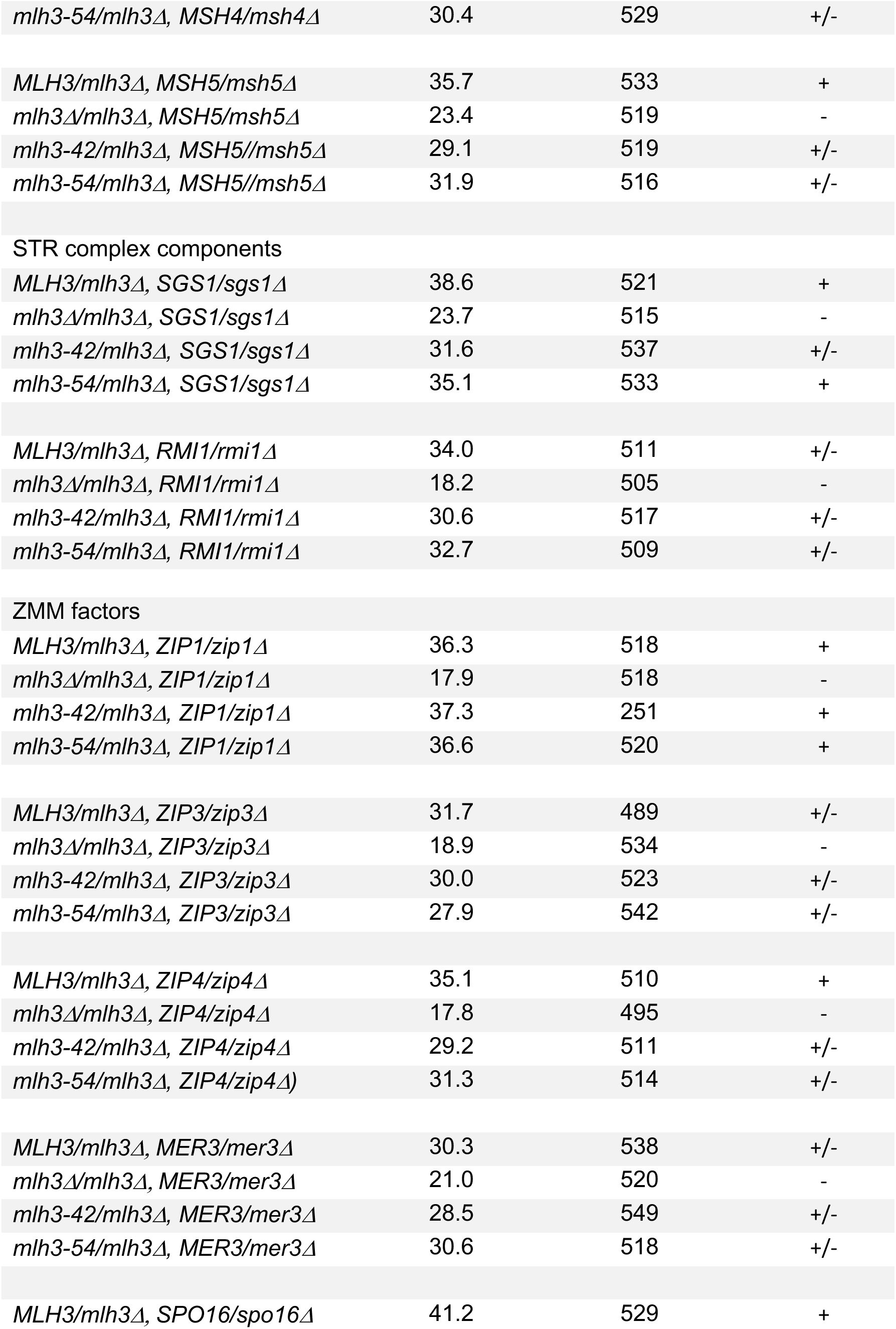
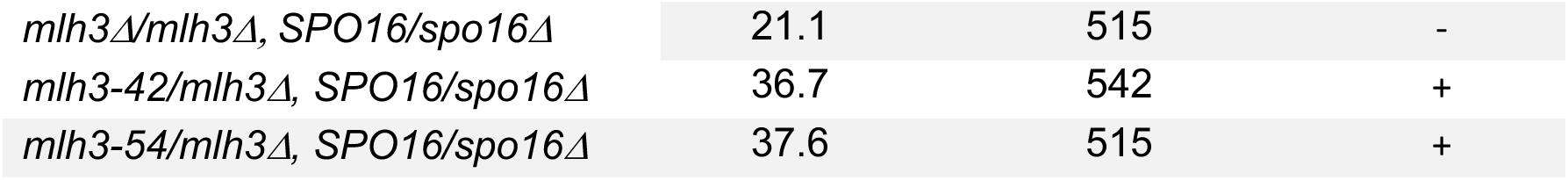
Analysis in diploid strains containing haploinsufficiency of *EXO1, MLH1, MSH4, MSH5, SGS1, RMI1, ZIP1, ZIP3, ZIP4, SPO16, MER3* genes in *mlh3-42* and *mlh3-54* strain backgrounds. Strains with the indicated relevant genotypes (Table S5) containing the *THR1::m-Cerulean-TRP1* and *CEN8::tdTomato-LEU2* markers on chromosome VIII were induced for meiosis and % tetratype in the *CEN8-THR1* interval was measured by determining the total tetratypes/sum of tetratypes and parental ditypes). At least two transformants were analyzed for each background. Significance was assessed by *χ*^2^ test between mutant and wild-type *EXO1 and exo1Δ* tetratype values. To minimize α inflation due to multiple comparisons, we applied a Benjamini-Hochberg correction at a 5% false discovery rate. +, indistinguishable from WT; -, indistinguishable from *exo1Δ*; +/-, distinguishable from both *wild-type* and *exo1Δ*.

**Table S5.**
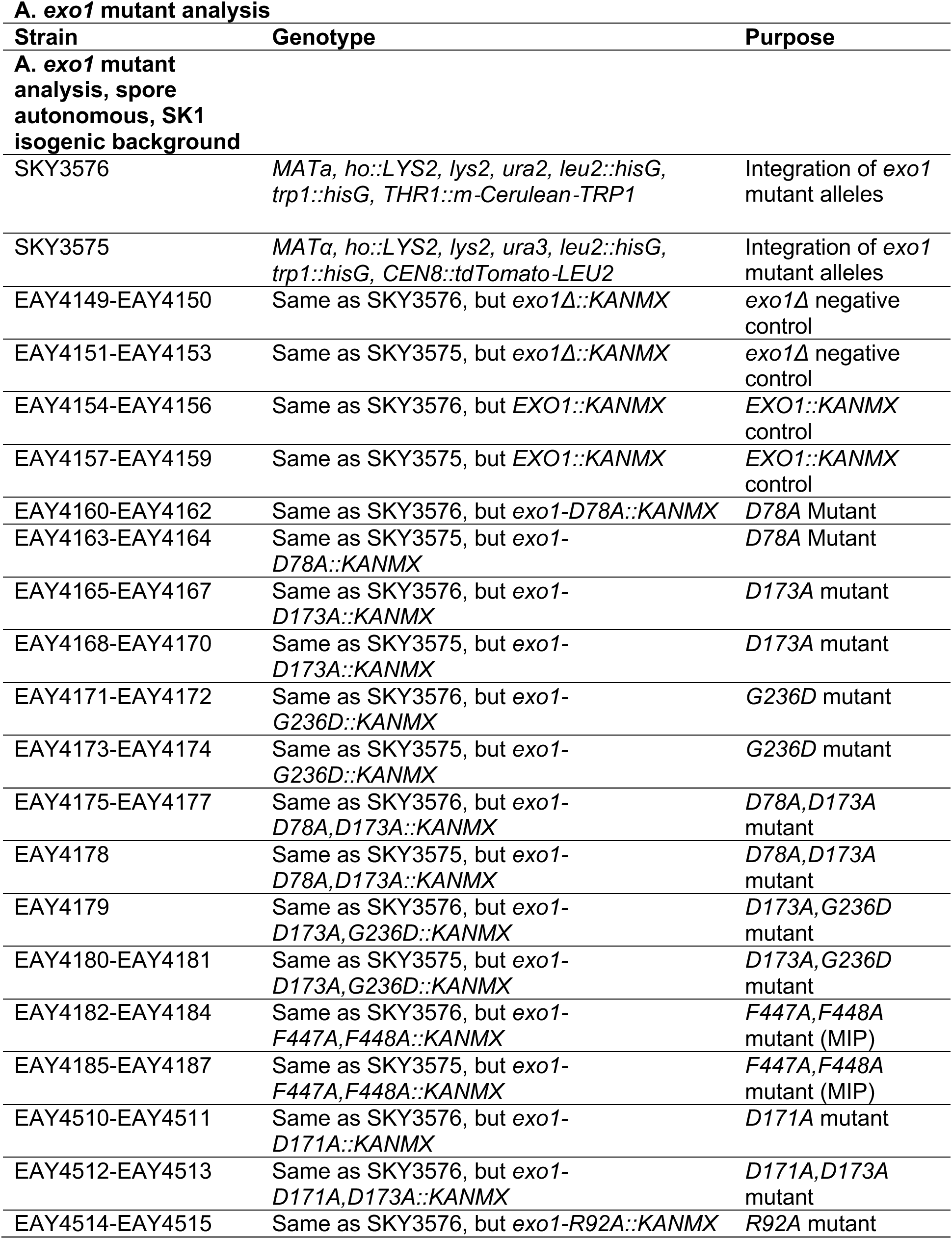
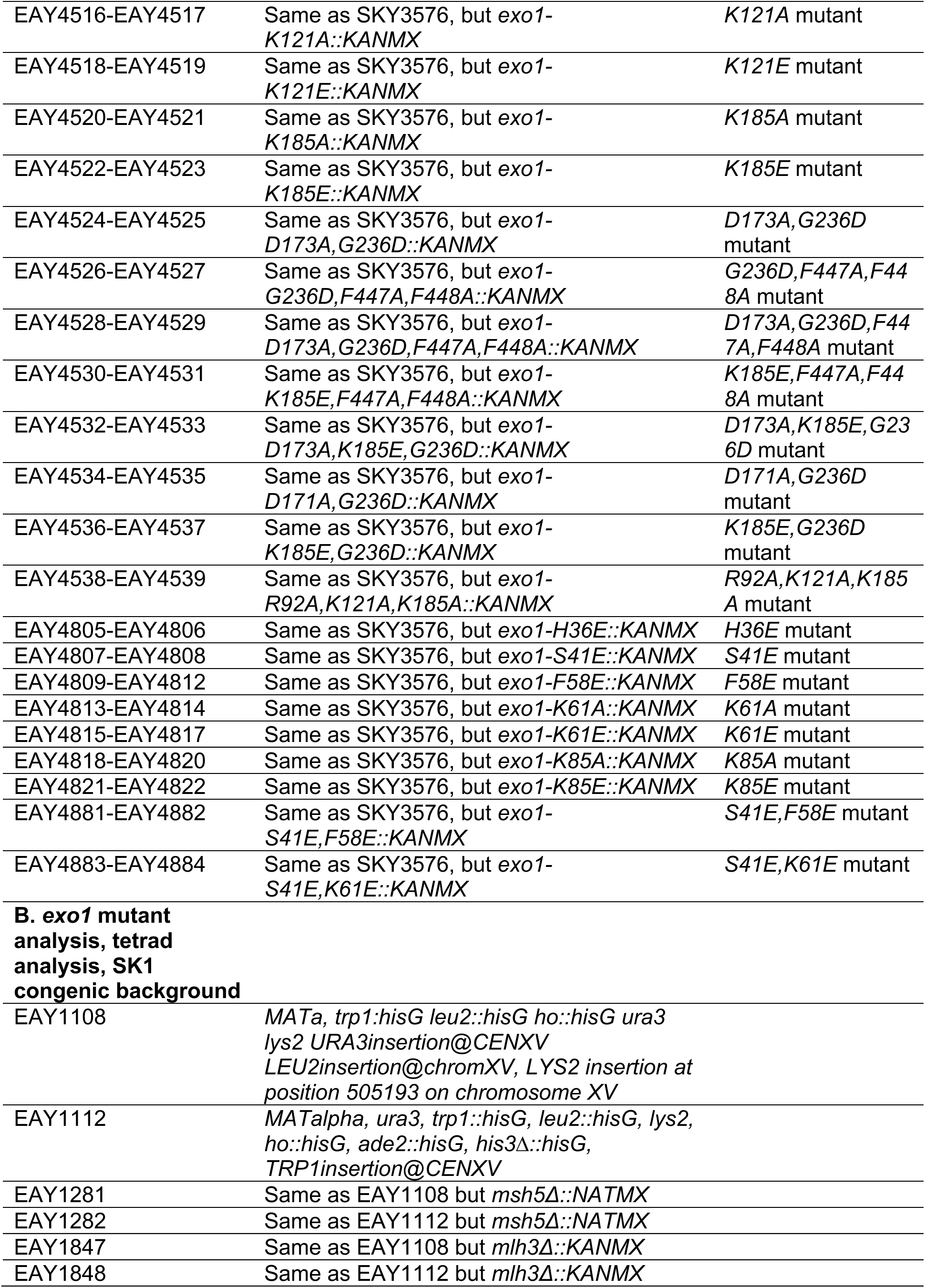
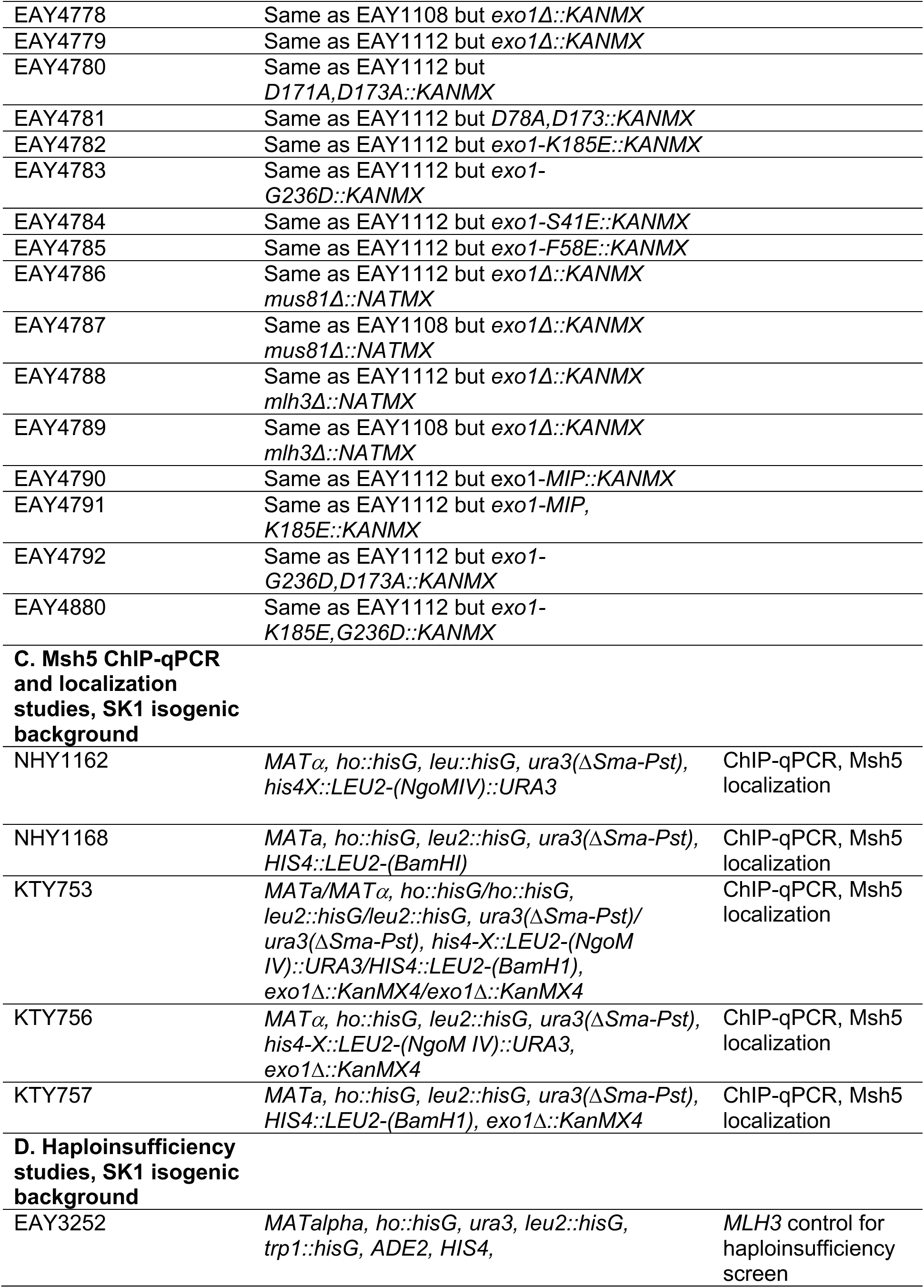
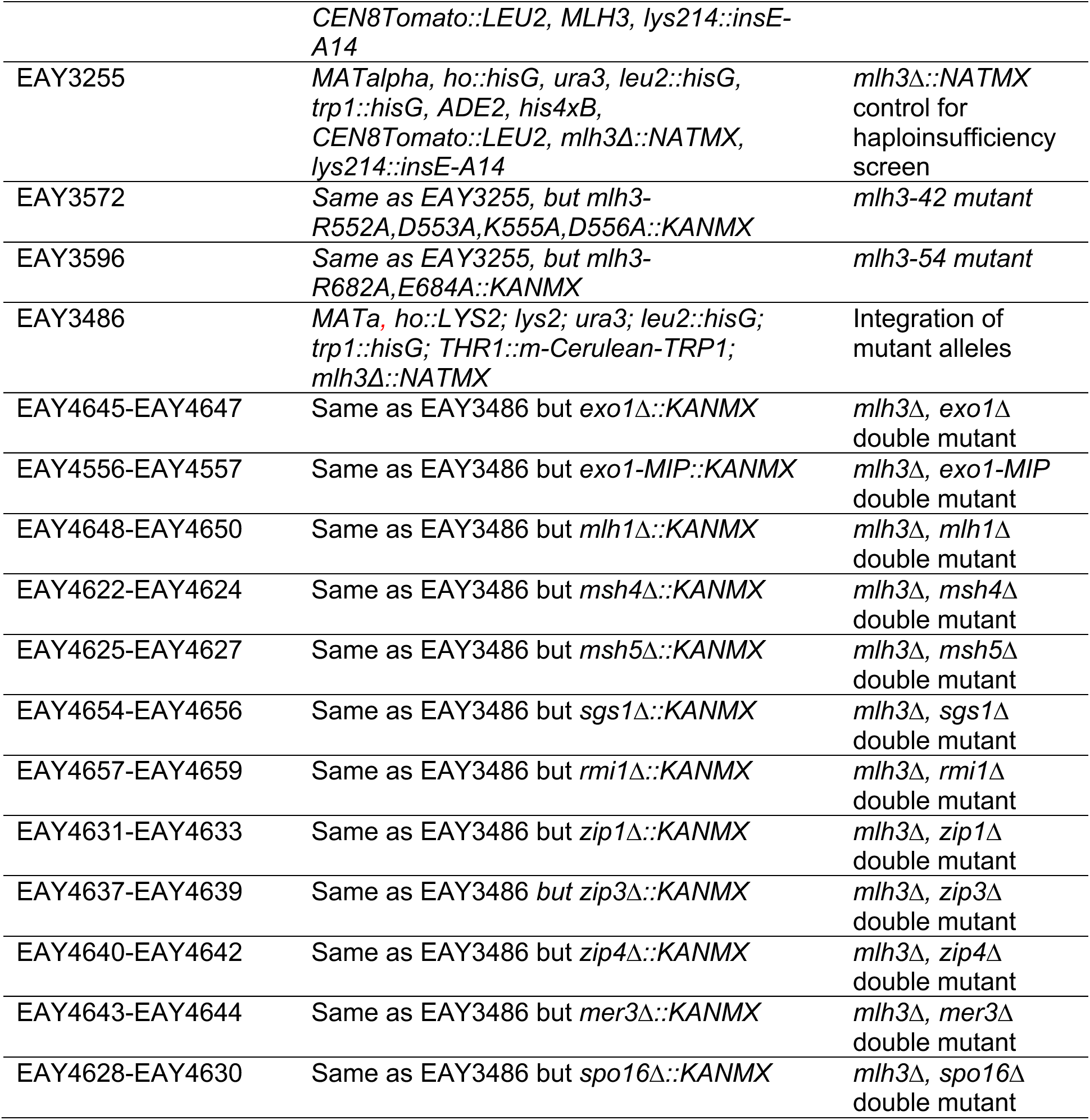
Strains used in this study.

**Table S6.** Plasmids used in this study.

| **Plasmid** | **Markers** | **Purpose** |
| --- | --- | --- |
| pUC18 | *amp<sup>R</sup>* | Exo1 endonuclease assay substrate |
| pBR322 | *amp<sup>R</sup>* | Exo1 endonuclease assay substrate |
| pRS416 | *amp<sup>R</sup>, URA3, CEN6-ARSH4* | Empty vector control |
| pLZ259 | *amp<sup>R</sup>, NATMX, CEN6-ARSH4* | Empty vector control |
| pRS426 | *amp<sup>R</sup>, URA3, 2μ* | Empty vector control |
| pEAI422 | *amp<sup>R</sup>, KANMX* | Integration of *exo1Δ-KANMX* |
| pEAI423 | *amp<sup>R</sup>, KANMX* | Integration of *EXO1-KANMX* |
| pEAI442 | *amp<sup>R</sup>, KANMX* | Integration of *exo1-H36E* |
| pEAI471 | *amp<sup>R</sup>, KANMX* | Integration of *exo1-S41E* |
| pEAI472 | *amp<sup>R</sup>, KANMX* | Integration of *exo1-F58E* |
| pEAI473 | *amp<sup>R</sup>, KANMX* | Integration of *exo1-K61A* |
| pEAI474 | *amp<sup>R</sup>, KANMX* | Integration of *exo1-K61E* |
| pEAI444 | *amp<sup>R</sup>, KANMX* | Integration of *exo1-K85A* |
| pEAI475 | *amp<sup>R</sup>, KANMX* | Integration of *exo1-K85E* |
| pEAI476 | *amp<sup>R</sup>, KANMX* | Integration of *exo1-S41E,F58E* |
| pEAI478 | *amp<sup>R</sup>, KANMX* | Integration of *exo1-S41E,K61E* |
| pEAI424 | *amp<sup>R</sup>, KANMX* | Integration of *exo1-D78A* |
| pEAI445 | *amp<sup>R</sup>, KANMX* | Integration of *exo1-R92A* |
| pEAI446 | *amp<sup>R</sup>, KANMX* | Integration of *exo1-K121A* |
| pEAI448 | *amp<sup>R</sup>, KANMX* | Integration of *exo1-K121E* |
| pEAI447 | *amp<sup>R</sup>, KANMX* | Integration of *exo1-D171A* |
| pEAI425 | *amp<sup>R</sup>, KANMX* | Integration of *exo1-D173A* |
| pEAI450 | *amp<sup>R</sup>, KANMX* | Integration of *exo1-K185A* |
| pEAI451 | *amp<sup>R</sup>, KANMX* | Integration of *exo1-K185E* |
| pEAI426 | *amp<sup>R</sup>, KANMX* | Integration of *exo1-G236D* |
| pEAI437 | *amp<sup>R</sup>, KANMX* | Integration of *exo1-F447A,F448A (MIP)* |
| pEAI427 | *amp<sup>R</sup>, KANMX* | Integration of *exo1-D78A,D173A* |
| pEAI449 | *amp<sup>R</sup>, KANMX* | Integration of *exo1-D171A,D173A* |
| pEAI456 | *amp<sup>R</sup>, KANMX* | Integration of *exo1-D171A,G236D* |
| pEAI436 | *amp<sup>R</sup>, KANMX* | Integration of *exo1-D173A,G236D* |
| pEAI458 | *amp<sup>R</sup>, KANMX* | Integration of *exo1-D173A,G236D,F447A,F448A (MIP)* |
| pEAI452 | *amp<sup>R</sup>, KANMX* | Integration of *exo1-G236D,F447A,F448A (MIP)* |
| pEAI467 | *amp<sup>R</sup>, KANMX* | Integration of *exo1-K185E,F447A,F448A (MIP)* |
| pEAI460 | *amp<sup>R</sup>, KANMX* | Integration of *exo1-D173A,K185E,G236D* |
| pEAI461 | *amp<sup>R</sup>, KANMX* | Integration of *exo1-K185E,G236D* |
| pEAI466 | *amp<sup>R</sup>, KANMX* | Integration of *exo1-R92A,K121A,K185A* |
| pEAA715 | *amp<sup>R</sup>, URA3, CEN6-ARSH4, EXO1* | *EXO1* complementation |
| pEAI483 | *amp<sup>R</sup>, NATMX, CEN6-ARSH4, EXO1* | *EXO1* complementation |
| pEAA726 | *amp<sup>R</sup>, URA3, CEN6-ARSH4, MLH3* | *MLH3* complementation |
| pEAA636 | *amp<sup>R</sup>, HIS3, CEN6-ARSH4, MLH3, KANMX* | *MLH3* complementation |
| pEAA722 | *amp<sup>R</sup>, URA3, CEN6-ARSH4, RAD27* | *RAD27* expression, native promoter |
| pEAA720 | *amp<sup>R</sup>, URA3, CEN6-ARSH4, pEXO1-RAD27* | *RAD27* expression under *EXO1* promoter |
| pEAI482 | *amp<sup>R</sup>, NATMX, CEN6-ARSH4, pEXO1-RAD27* | Expression of *RAD27* under *EXO1* promoter |
| pEAA727 | *amp<sup>R</sup>, URA3, CEN6-ARSH4, pEXO1-rad27-A45E* | *rad27-A45E* expression under *EXO1* promoter |
| pEAA728 | *amp<sup>R</sup>, URA3, CEN6-ARSH4, pEXO1-rad27-R101A* | *rad27-R101A* expression under *EXO1* promoter |
| pEAA729 | *amp<sup>R</sup>, URA3, CEN6-ARSH4, pEXO1-rad27-R105A* | *rad27-R105A* expression under *EXO1* promoter |
| pEAA730 | *amp<sup>R</sup>, URA3, CEN6-ARSH4, pEXO1-rad27-K130A* | *rad27-K130A* expression under *EXO1* promoter |
| pEAA724 | *amp<sup>R</sup>, URA3, CEN6-ARSH4, pEXO1-rad27-D179A* | *rad27-D179A* expression under *EXO1* promoter |
| pEAA731 | *amp<sup>R</sup>, URA3, CEN6-ARSH4, pEXO1-rad27-H191E* | *rad27-H191E* expression under *EXO1* promoter |
| pEAM327 | *amp<sup>R</sup>, URA3, 2μ, CDC9* | *CDC9* expression, native promoter |
| pEAM329 | *amp<sup>R</sup>, URA3, 2μ, pHOP1-CDC9* | Overexpression of *CDC9* under the *HOP1* promoter |
| pFB-EXO1-FLAG | *amp<sup>R</sup>, Gm<sup>R</sup>, EXO1-FLAG* | *EXO1* expression from pFastBac (From Michael Liskay) |
| pFB-exo1-D173A-FLAG | *amp<sup>R</sup>, Gm<sup>R</sup>, exo1-D173A-FLAG* | *exo1-D173A* expression from pFastBac (From Michael Liskay) |
| pEAE422 | *amp<sup>R</sup>, Gm<sup>R</sup>, exo1-G236D-FLAG* | *exo1-G236D* expression from pFastBac |
| pEAE423 | *amp<sup>R</sup>, Gm<sup>R</sup>, exo1-D173A-G236D-FLAG* | *exo1-D173A, G236D* expression from pFastBac |

**Table S7.**
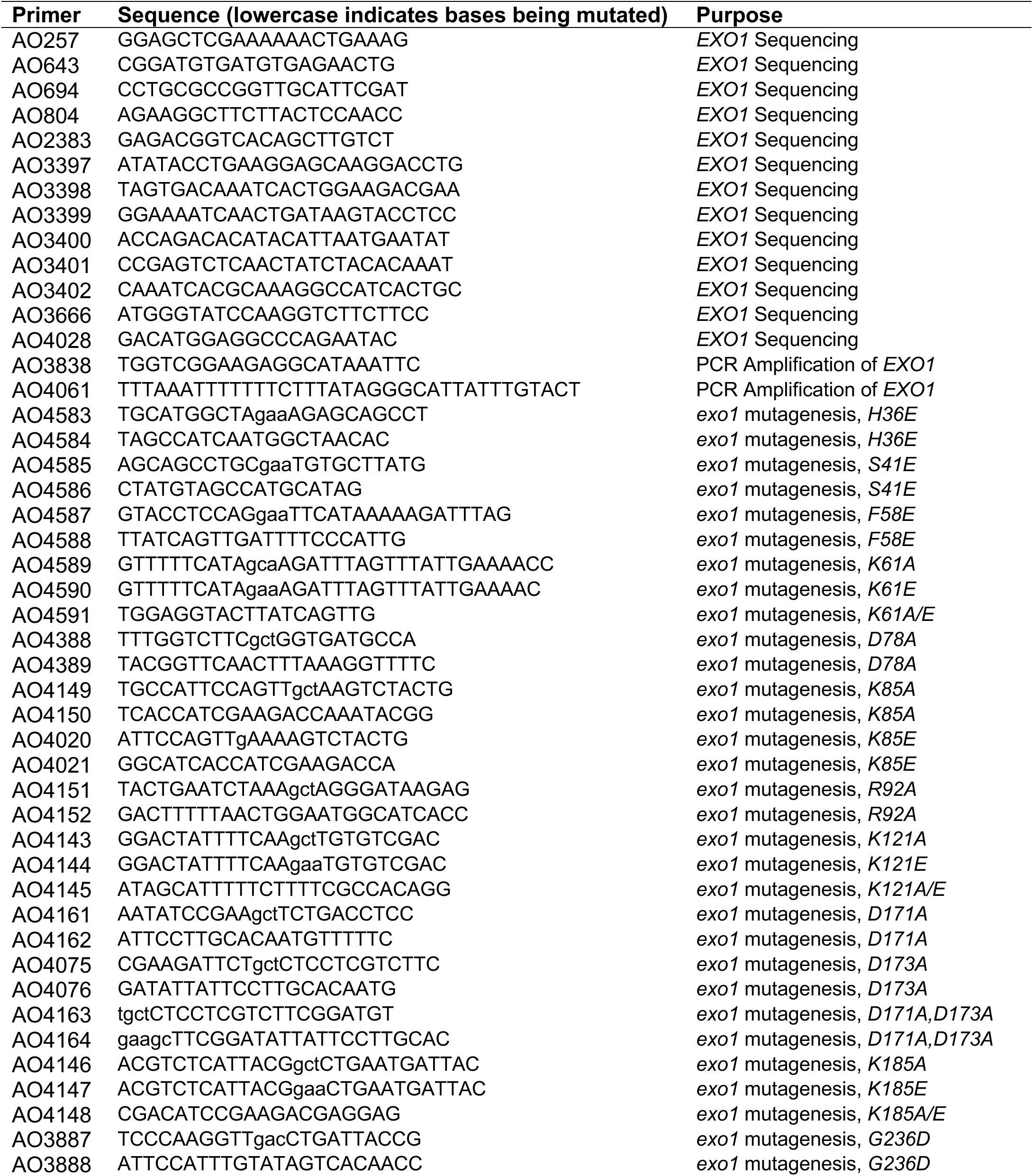
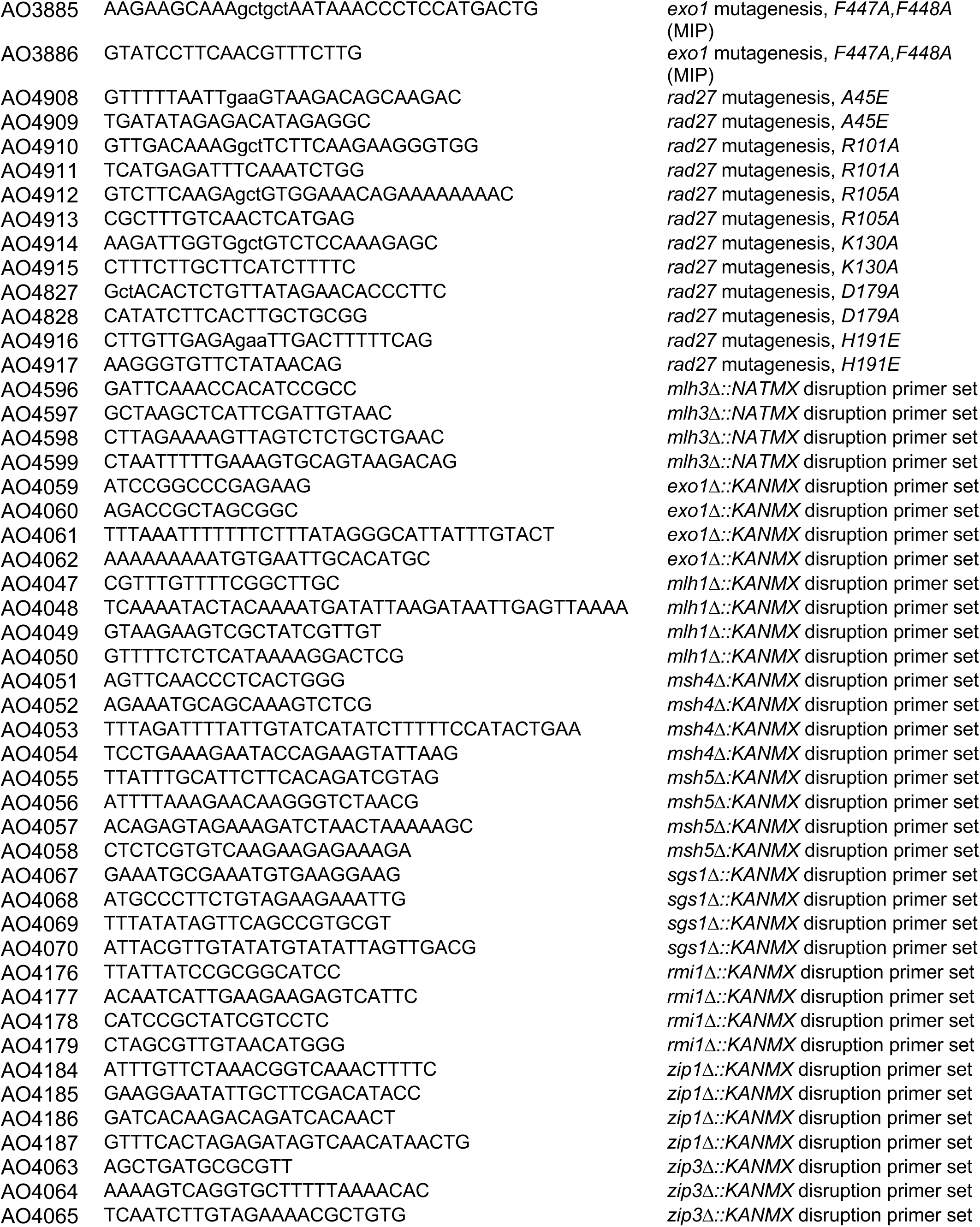
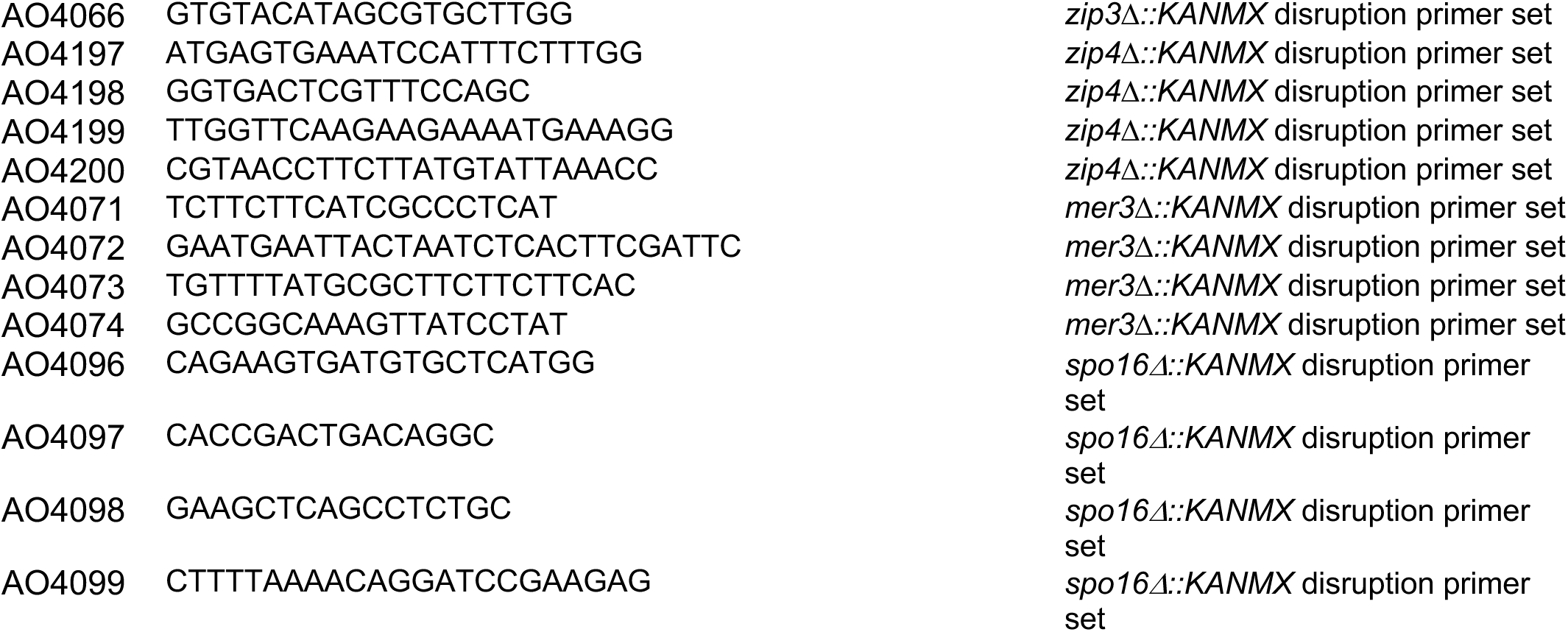
Oligonucleotides used in this study (shown 5’ to 3’).

## Notes

### Competing Interest Statement

The authors have declared no competing interest.

## REFERENCES

Abdullah, M.F., Hoffmann, E.R., Cotton, V.E., and Borts, R.H. (2004). A role for the MutL homologue MLH2 in controlling heteroduplex formation and in regulating between two different crossover pathways in budding yeast. Cytogenet. Genome Res. 107, 180–190.

Agarwal, S., and Roeder, G.S. (2000). Zip3 provides a link between recombination enzymes and synaptonemal complex proteins. Cell 102, 245–255.

Ahuja, J.S., Harvey, C.S., Wheeler, D.L., and Lichten, M. (2021). Repeated strand invasion and extensive branch migration are hallmarks of meiotic recombination. Mol. Cell, Ahead of Print; doi: 10.1016/j.molcel.2021.08.003

Al-Sweel, N., Raghavan, V., Datta, A., Ajith, V.P., Di Vietro, L., Khondakar, N., Manhart, C.M., Surtees, J.A., Nishant, K.T., and Alani, E. (2017). *mlh3* separation of function and endonuclease defective mutants display an unexpected effect on meiotic recombination outcomes. PLoS Genet. 13, e1006974.

Amin, N.S., Nguyen, M., Oh, S., and Kolodner, R.D. (2001). *exo1*-dependent mutator mutation: Model system for studying functional interactions in mismatch repair. Mol. Cell. Biol. 21, 5142–5155.

Argueso, J.L., Kijas, A.W., Sarin, S., Heck, J., Waase, M., and Alani, E. (2003). Systematic mutagenesis of the *Saccharomyces cerevisiae MLH1* gene reveals distinct roles for Mlh1p in meiotic crossing over and in vegetative and meiotic mismatch repair. Mol. Cell. Biol. 23, 873–886.

Argueso, J.L., Wanat, J., Gemici, Z., and Alani, E. (2004). Competing crossover pathways act during meiosis in *Saccharomyces cerevisiae*. Genetics 168, 1805–1816.

Arter, M., Hurtado-Nieves, V., Oke, A., Zhuge, T., Wettstein, R., Fung, J.C., Blanco, M.G., and Matos, J. (2018). Regulated crossing-over requires inactivation of Yen1/GEN1 resolvase during meiotic prophase I. Dev. Cell 45, 785–800.e6.

Bahler, J., Wu, J.Q., Longtine, M.S., Shah, N.G., McKenzie, A. 3rd, Steever, A.B., Wach, A., Philippsen, P., and Pringle, J.R. (1998). Heterologous modules for efficient and versatile PCR-based gene targeting in *Schizosaccharomyces pombe*. Yeast 14, 943–951.

Balakrishnan, L. and Bambara, R. A. (2013). Flap endonuclease 1. Annu. Rev. Biochem. 82, 119–138.

Bell, L.R., and Byers, B. (1983). Homologous association of chromosomal DNA during yeast meiosis. Cold Spring Harb. Symp. Quant. Biol. 47, 829–40. doi: 10.1101/sqb.1983.047.01.095.

Benjamini, Y., and Hochberg, Y. (1995). Controlling the false discovery rate: A practical and powerful approach to multiple testing. J. R. Statist. Soc. Ser. B. 57, 289–300.

Bhagwat, N.R., Owens, S.N., Ito, M., Boinapalli, J.V., Poa, P., Ditzel, A., Kopparapu, S., Mahalawat, M., Davies, O.R., Collins, S.R., Johnson, J.R., Kroganm, N.J., and Hunter, N. (2021). SUMO is a pervasive regulator of meiosis Elife 10, e57720.

Bishop, D.K. (1994). RecA homologs Dmc1 and Rad51 interact to form multiple nuclear complexes prior to meiotic chromosome synapsis. Cell 79, 1081–1092. doi: 10.1016/0092-8674(94)90038-8.

Börner, G.V., Kleckner, N., and Hunter, N. (2004). Crossover/noncrossover differentiation, synaptonemal complex formation, and regulatory surveillance at the leptotene/zygotene transition of meiosis. Cell 117, 29–45.

Bradford, M.M. (1976). A rapid and sensitive method for the quantitation of microgram quantities of protein utilizing the principle of protein-dye binding. Anal. Biochem. 72, 248–254.

Brar, G.A., Yassour, M., Friedman, N., Regev, A., Ingolia, N.T., and Weissman, J.S. (2012). High-resolution view of the yeast meiotic program revealed by ribosome profiling. Science 335, 552–557.

Cannavo, E., Sanchez, A., Anand, R., Ranjha, L., Hugener, J., Adam, C., Acharya, A., Weyland, N., Aran-Guiu, X., Charbonnier, J.B., et al. (2020). Regulation of the MLH1–MLH3 endonuclease in meiosis. Nature 586, 618–622.

Cao, L., Alani, E., and Kleckner, N. (1990). A pathway for generation and processing of double-strand breaks during meiotic recombination in *S. cerevisiae*. Cell 61, 1089–1101.

Ceska, T.A., Sayers, J.R., Stier, G., and Suck, D. (1996). A helical arch allowing single-stranded DNA to thread through T5 5’-exonuclease. Nature 382, 90–93.

Challa, K., Fajish, V.G., Shinohara, M., Klein, F., Gasser, S.M., and Shinohara, A. (2019). Meiosis-specific prophase-like pathway controls cleavage-independent release of cohesin by Wapl phosphorylation. PLoS Genet.15, e1007851. doi: 10.1371/journal.pgen.1007851.

Chapados, B.R., Hosfield, D.J., Han, S., Qiu, J., Yelent, B., Shen, B., and Tainer, J.A. (2004). Structural basis for FEN-1 substrate specificity and PCNA-mediated activation in DNA replication and repair. Cell 116, 39–50. doi: 10.1016/s0092-8674(03)01036-5.

Christianson, T.W., Sikorski, R.S., Dante, M., Shero, J.H. and Hieter, P. (1992). Multifunctional yeast high-copy-number shuttle vectors. Gene 110, 119–122.

Cole, F., Kauppi, L., Lange, J., Roig, I., Wang, R., Keeney, S., and Jasin, M. (2012). Homeostatic control of recombination is implemented progressively in mouse meiosis. Nat. Cell Biol. 14, 424–430. doi: 10.1038/ncb2451.

Dai, J., Sanchez, A., Adam, C., Ranjha, L., Reginato, G., Chervy, P., Tellier-Lebegue, C., Andreani, J., Guérois, R., Ropars, V., Le Du, M.H., Maloise, l.L., Martini, E., Legrand, P., Thureau, A., Cejka, P., Borde, V., and Charbonnier, J.B. (2021). Molecular basis of the dual role of the Mlh1-Mlh3 endonuclease in MMR and in meiotic crossover formation. Proc. Natl. Acad. Sci. USA 118, e2022704118. doi: 10.1073/pnas.2022704118.

de Boer, E., and Heyting, C. (2006). The diverse roles of transverse filaments of synaptonemal complexes in meiosis. Chromosoma 115, 220–234.

de los Santos, T., Loidl, J., Larkin, B., and Hollingsworth, N.M. (2001). A role for MMS4 in the processing of recombination intermediates during meiosis in *Saccharomyces cerevisiae*. Genetics 159, 1511–1525.

De Muyt, A., Pyatnitskaya, A., Andréani, J., Ranjha, L., Ramus, C., Laureau, R., Fernandez-Vega, A., Holoch, D., Girard, E., Govin, J., Margueron, R., Couté, Y., Cejka, P., Guérois, R., and Borde, V. (2018). A meiotic XPF-ERCC1-like complex recognizes joint molecule recombination intermediates to promote crossover formation. Genes Dev. 32, 283–296.

Devos, J.M., Tomanicek, S.J., Jones, C.E., Nossal, N.G., and Mueser, T.C. (2007). Crystal structure of bacteriophage T4 5’ nuclease in complex with a branched DNA reveals how flap endonuclease-1 family nucleases bind their substrates. J. Biol. Chem. 282, 31713–31724.

Feng, M., Patel, D., Dervan, J.J., Ceska, T., Suck, D., Haq, I., and Sayers, J.R. (2004). Roles of divalent metal ions in flap endonuclease–substrate interactions. Nat. Struct. Mol. Biol. 11, 450–456.

Fiorentini, P., Huang, K.N., Tishkoff, D.X., Kolodner, R.D., and Symington, L.S. (1997). Exonuclease I of *Saccharomyces cerevisiae* functions in mitotic recombination *in vivo* and *in vitro*. Mol. Cell. Biol. 17, 2764–2773.

Fricke, W.M., Bastin-Shanower, S.A., and Brill, S.J. (2005). Substrate specificity of the Saccharomyces cerevisiae Mus81-Mms4 endonuclease. DNA Repair 4, 243–251. doi: 10.1016/j.dnarep.2004.10.001.

Fung, J.C., Rockmill, B., Odell, M., and Roeder, G.S. (2004). Imposition of crossover interference through the nonrandom distribution of synapsis initiation complexes. Cell 116, 795–802.

García-Luis, J., and Machín, F. (2014). Mus81-Mms4 and Yen1 resolve a novel anaphase bridge formed by noncanonical Holliday junctions. Nat. Commun. 5, 5652. doi: 10.1038/ncomms6652.

Gary, R., Park, M.S., Nolan, J.P., Cornelius, H.L., Kozyreva, O.G., Tran, H.T., Lobachev, K.S., Resnick, M.A., and Gordenin, D.A. (1999). A novel role in DNA metabolism for the binding of Fen1/Rad27 to PCNA and implications for genetic risk. Mol Cell Biol. 19, 5373–5382.

Genschel, J., and Modrich, P. (2003). Mechanism of 5′-Directed Excision in Human Mismatch Repair. Mol. Cell 12, 1077–1086.

Giaever, G., and Nislow, C. (2014). The yeast deletion collection: A decade of functional genomics. Genetics 197, 451–465.

Gietz, R.D., Schiestl, R.H., Willems, A.R., and Woods, R.A. (1995). Studies on the transformation of intact yeast cells by the LiAc/SS-DNA/PEG procedure. Yeast 11, 355–360.

Gloor, J.W., Balakrishnan, L., and Bambara R.A. (2010). Flap endonuclease 1 mechanism analysis indicates flap base binding prior to threading. J. Biol. Chem. 285, 34922–34931. doi: 10.1074/jbc.M110.165902.

Goellner, E.M., Putnam, C.D., Graham, W.J. 5th, Rahal, C.M., Li, B.-Z., and Kolodner, R.D. (2018). Identification of Exo1-Msh2 interaction motifs in DNA mismatch repair and new Msh2-binding partners. Nat Struct Mol Biol. 25, 650–659.

Goellner, E.M., Putnam, C.D., and Kolodner, R.D. (2015). Exonuclease 1-dependent and independent mismatch repair. DNA Repair 32, 24–32.

Goldstein, A.L., and McCusker, J.H. (1999). Three new dominant drug resistance cassettes for gene disruption in *Saccharomyces cerevisiae*. Yeast 15, 1541–1553.

Hall, M.C., Wang, H., Erie, D.A., and Kunkel, T.A. (2001). High affinity cooperative DNA binding by the yeast Mlh1-Pms1 heterodimer. J. Mol. Biol. 312, 637–647.

Hassold, T., and Hunt, P. (2001). To err (meiotically) is human: the genesis of human aneuploidy. Nat. Rev. Genet. 2, 280–291.

Hatkevich, T., and Sekelsky, J. (2017). Bloom syndrome helicase in meiosis: Pro-crossover functions of an anti-crossover protein. BioEssays 39, 1–22.

Hillers, K.J. (2004). Crossover interference. Curr. Biol. 14, R1036–R1037.

Hoffman, C.S., and, Winston, F. (1987). A ten-minute DNA preparation from yeast efficiently releases autonomous plasmids for transformation of *Escherichia coli*. Gene 57, 267–272.

Hollingsworth, N.M., Ponte, L., and Halsey, C. (1995). MSH5, a novel MutS homolog, facilitates meiotic reciprocal recombination between homologs in *Saccharomyces cerevisiae* but not mismatch repair. Genes Dev. 9, 1728–1739.

Hunter, N. (2015). Meiotic recombination: The essence of heredity. Cold Spring Harb. Perspect. Biol. 7, a016618.

Hunter, N., and Kleckner, N. (2001). The single-end invasion: An asymmetric intermediate at the double-strand break to double-Holliday Junction transition of meiotic recombination. Cell 106, 59–70.

Hwang, K.Y., Baek, K., Kim, H.-Y., and Cho, Y. (1998). The crystal structure of flap endonuclease-1 from *Methanococcus jannaschii*. Nat. Struct. Mol. Biol. 5, 707–713.

Ip, S.C., Rass, U., Blanco, M.G., Flynn, H.R., Skehel, J.M., and West, S.C. (2008). Identification of Holliday Junction resolvases from humans and yeast. Nature 456, 357–361.

Jessop, L., Rockmill, B., Roeder, G.S., and Lichten, M. (2006). Meiotic chromosome synapsis-promoting rroteins antagonize the anti-crossover activity of Sgs1. PLoS Genet. 2, e155.

Jones, G. H., and Franklin, F.C.H. (2006). Meiotic crossing-over: Obligation and interference. Cell 126, 246–248.

Kane, S.M. and Roth, R. (1974) Carbohydrate metabolism during ascospore development in yeast. J. Bacteriol. 118, 8–14.

Katoh, K., Rozewicki, J., and Yamada, K.D. (2019). MAFFT online service: multiple sequence alignment, interactive sequence choice and visualization. Brief. Bioinform. 20, 1160–1166.

Kaur, H., De Muyt, A., and Lichten, M. (2015). Top3-Rmi1 DNA Single-Strand Decatenase Is Integral to the formation and resolution of meiotic recombination intermediates. Mol. Cell 57, 583–594.

Keelagher, R.E., Cotton, V.E., Goldman, A.S. and Borts, R.H. (2011). Separable roles for Exonuclease I in meiotic DNA double-strand break repair. DNA Repair 10, 126–137. DOI:10.1016/j.dnarep.2010.09.024.

Keeney, S., Giroux, C.N., and Kleckner, N. (1997). Meiosis-specific DNA double-strand breaks are catalyzed by Spo11, a member of a widely conserved protein family. Cell 88, 375–384.

Khazanehdari, K.A., and Borts, R.H. (2000). EXO1 and MSH4 differentially affect crossing-over and segregation. Chromosoma 109, 94–102.

Kim, Y., Furman, C.M., Manhart, C.M., Alani, E. and Finkelstein, I.J. (2019). Intrinsically disordered regions regulate both catalytic and non-catalytic activities of the MutLα mismatch repair complex. Nucleic Acids Res. 47, 1823–1835.

Kolas, N.K., Svetlanov, A., Lenzi, M.L., Macaluso, F.P., Lipkin, S.M., Liskay, R.M., Greally, J., Edelmann, W., and Cohen, P.E. (2005). Localization of MMR proteins on meiotic chromosomes in mice indicates distinct functions during prophase I. J. Cell Biol. 171, 447–458.

Krishnaprasad, G.N., Salim, S., Pankajam, A.V., Shinohara, M., Lin, G., Chakraborty, P., Farnaz, A., Steinmetz, L.M., Shinohara, A, and Nishant, K.T. (2021). Regulation of Msh4-Msh5 association with meiotic chromosomes in budding yeast. Genetics, In press, https://doi.org/10.1093/genetics/iyab102.

Kulkarni, D.S., Owens, S.N., Honda, M., Ito, M., Yang, Y., Corrigan, M.W., Chen, L., Quan, A.L., and Hunter, N. (2020). PCNA activates the MutL endonuclease to promote meiotic crossing over. Nature 586, 623–627.

Kunkel, T.A., and Erie, D.A. (2015). Eukaryotic Mismatch Repair in Relation to DNA Replication. Annu. Rev. Genet. 49, 291–313.

Lee, B.-I., Nguyen, L.H., Barsky, D., Fernandes, M., and Wilson, D.M. 3rd. (2002). Molecular interactions of human Exo1 with DNA. Nucleic Acids Res. 30, 942–949.

Lee, B.-I., and Wilson, D.M. (1999). The RAD2 domain of human exonuclease 1 exhibits 5′ to 3′ exonuclease and flap structure-specific endonuclease activities. J. Biol. Chem. 274, 37763–37769.

Li, Y., Shen, J., and Niu, H. (2019). DNA duplex recognition activates Exo1 nuclease activity. J. Biol. Chem. 294, 11559–11567.

Lynn, A., Soucek, R., and Börner, G.V. (2007). ZMM proteins during meiosis: Crossover artists at work. Chromosome Res. 15, 591–605.

Machín, F. (2020). Implications of metastable nicks and nicked Holliday Junctions in processing joint molecules in mitosis and meiosis. Genes 11, 1498. doi: 10.3390/genes11121498.

Maguire, M.P. (1974). The need for a chiasma binder. J. Theor. Biol. 48, 485–487.

Malkova, A., Swanson, J., German, M., McCusker, J.H., Housworth, E.A., Stahl, F.W., and Haber, J.E. (2004). Gene conversion and crossing over along the 405-kb left arm of Saccharomyces cerevisiae chromosome VII. Genetics 168, 49–63.

Mancera, E., Bourgon, R., Brozzi, A., Huber, W., and Steinmetz, L.M. (2008). High-resolution mapping of meiotic crossovers and non-crossovers in yeast. Nature 454, 479–485.

Manhart, C.M., Ni, X., White, M.A., Ortega, J., Surtees, J.A., and Alani, E. (2017). The mismatch repair and meiotic recombination endonuclease Mlh1-Mlh3 is activated by polymer formation and can cleave DNA substrates in trans. PLoS Biol. 15, e2001164.

Marsolier-Kergoat, M.C., Khan, M.M., Schott, J., Zhu, X., and Llorente, B. (2018). Mechanistic view and genetic control of DNA recombination during meiosis. Mol. Cell 70, 9–20.

Martini, E., Borde, V., Legendre, M., Audic, S., Regnault, B., Soubigou, G., Dujon, B., and Llorente, B. (2011). Genome-wide analysis of heteroduplex DNA in mismatch repair–deficient yeast cells reveals novel properties of meiotic recombination pathways. PLoS Genet. 7, e1002305.

Martini, E., Diaz, R.L., Hunter, N., and Keeney, S. (2006). Crossover homeostasis in yeast meiosis. Cell 126, 285–295.

McDonald, J.H. (2014). Handbook of Biological Statistics (3rd ed.). Sparky House Publishing, Baltimore, Maryland.

Mueser, T.C., Nossal, N.G., and Hyde, C.C. (1996). Structure of bacteriophage T4 RNase H, a 5′ to 3′ RNA–DNA and DNA–DNA exonuclease with sequence Ssimilarity to the RAD2 family of eukaryotic proteins. Cell 85, 1101–1112.

Nagaoka, S.I., Hassold, T.J., and Hunt, P.A. (2012). Human aneuploidy: mechanisms and new insights into an age-old problem. Nat. Rev. Genet. 13, 493–504.

Nicolette, M.L., Lee, K., Guo, Z., Rani, M., Chow, J.M., Lee, S.E., and Paull, T.T. (2010). Mre11-Rad50-Xrs2 and Sae2 promote 5’ strand resection of DNA double-strand breaks. Nat. Struct. Mol. Biol. 17, 1478–1485.

Nishant, K.T., Chen, C., Shinohara, M., Shinohara, A, and Alani, E. (2010). Genetic analysis of baker’s yeast Msh4-Msh5 reveals a threshold crossover level for meiotic viability. PLoS Genetics, 6, e1001083.

Nishant, K.T., Plys, A.J., and Alani, E. (2008). A mutation in the putative MLH3 endonuclease domain confers a defect in both mismatch repair and meiosis in Saccharomyces cerevisiae. Genetics 179, 747–755.

Novak, J.E., Ross-Macdonald, P.B., and Roeder, G.S. (2001). The Budding yeast Msh4 protein functions in chromosome synapsis and the regulation of crossover distribution. Genetics 158, 1013–1025.

Orans, J., McSweeney, E.A., Iyer, R.R., Hast, M.A., Hellinga, H.W., Modrich, P., and Beese, L.S. (2011). Structures of human exonuclease I DNA complexes suggest a unified mechanism for nuclease family. Cell 145, 212–223.

Padmore, R., Cao, L., and Kleckner, N. (1991). Temporal comparison of recombination and synaptonemal complex formation during meiosis in *S. cerevisiae*. Cell 66, 1239–1256.

Pan, J., Sasaki, M., Kniewel, R., Murakami, H., Blitzblau, H.G., Tischfield, S.E., Zhu, X., Neale, M.J., Jasin, M., Socci, N.D., Hochwagen, A., and Keeney, S. (2011). A Hierarchical Combination of Factors Shapes the Genome-Wide Topography of Yeast Meiotic Recombination Initiation. Cell 144, 719–731.

Papazian, H.P. (1952). The analysis of tetrad data. Genetics 37, 175–188.

Pelletier, H., M. R. Sawaya, W. Wolfle, S. H. Wilson, and J. Kraut, (1996). Crystal structures of human DNA polymerase β complexed with DNA: Implications for catalytic mechanism, processivity, and fidelity. Biochemistry 35, 12742–12761.

Perkins, D.D. (1949). Biochemical mutants in the smut fungus *Ustilago maydis*. Genetics 34, 607–626.

Peterson, S.E., Keeney, S., and Jasin, M. (2020). Mechanistic insight into crossing over during mouse meiosis. Mol. Cell. 78, 1252–1263.e3. doi: 10.1016/j.molcel.2020.04.009.

Qiu, J., Qian, Y., Chen, V., Guan, M.X., and Shen, B. (1999). Human exonuclease 1 functionally complements its yeast homologues in DNA recombination, RNA primer removal, and mutation avoidance. J Biol Chem. 274, 17893–17900. doi: 10.1074/jbc.274.25.17893.

Ranjha, L., Anand, R., and Cejka, P. (2014). The *Saccharomyces cerevisiae* Mlh1-Mlh3 heterodimer Is an endonuclease that preferentially binds to Holliday Junctions. J. Biol. Chem. 289, 5674–5686.

Reyes, G.X., Kolodziejczak, A., Devakumar, L.J.P.S., Kubota, T., Kolodner, R.D., Putnam, C.D., and Hombauer, H. (2021). Ligation of newly replicated DNA controls the timing of DNA mismatch repair. Curr. Biol. 31, 1268–1276.e6. doi: 10.1016/j.cub.2020.12.018.

Rogacheva, M.V, Manhart, C.M., Chen, C., Guarne, A., Surtees, J., and Alani, E. (2014). Mlh1-Mlh3, A meiotic crossover and DNA mismatch repair factor, is a Msh2-Msh3-stimulated endonuclease. J. Biol. Chem. 289, 5664–5673.

Rose, M.D., Winston, F., and Hieter, P. (1990). Methods in yeast genetics: A laboratory course manual. Cold Spring Harbor Laboratory Press, Cold Spring Harbor, NY.

Ross-Macdonald, P., and Roeder, G.S. (1994). Mutation of a meiosis-specific MutS homolog decreases crossing over but not mismatch correction. Cell 79, 1069–1080.

Sanchez, A., Adam, C., Rauh, F., Duroc, Y., Ranjha, L., Lombard, B., Mu, X., Wintrebert, M., Loew, D., Guarné, A., et al. (2020). Exo1 recruits Cdc5 polo kinase to MutL to ensure efficient meiotic crossover formation. Proc. Natl. Acad. Sci. USA 117, 30577–30588.

Santucci-Darmanin, S., Walpita, D., Lespinasse, F., Desnuelle, C., Ashley, T., and Paquis-Flucklinger, V. (2000). MSH4 acts in conjunction with MLH1 during mammalian meiosis. FASEB J. 14, 1539–1547 (2000).

Santucci-Darmanin, S., Neyton, S., Lespinasse, F., Saunières, A., Gaudray, P., and Paquis-Flucklinger, V. (2002). The DNA mismatch-repair MLH3 protein interacts with MSH4 in meiotic cells, supporting a role for this MutL homolog in mammalian meiotic recombination. Hum. Mol. Genet. 11, 1697–1706.

Schiltz, C.J., Lee, A., Partlow, E.A., Hosford, C.J., and Chappie, J.S. (2019). Structural characterization of Class 2 OLD family nucleases supports a two-metal catalysis mechanism for cleavage. Nucleic Acids Res. 47, 9448–9463. doi: 10.1093/nar/gkz703.

Schwacha, A., and Kleckner, N. (1994). Identification of joint molecules that form frequently between homologs but rarely between sister chromatids during yeast meiosis. Cell 76, 51–63.

Schwacha, A., and Kleckner, N. (1995). Identification of double Holliday Junctions as intermediates in meiotic recombination. Cell 83, 783–791.

Shen, B., Nolan, J.P., Sklar, L.A., and Park, M.S. (1996). Essential amino acids for substrate binding and catalysis of human flap endonuclease 1. J. Biol. Chem. 271, 9173–9176.

Shi, Y., Hellinga, H.W., and Beese, L.S. (2017). Interplay of catalysis, fidelity, threading, and processivity in the exo- and endonucleolytic reactions of human exonuclease I. Proc. Natl. Acad. Sci. 114, 6010–6015.

Shinohara, M., Sakai, K., Shinohara, A., and Bishop, D.K. (2003). Crossover interference in Saccharomyces cerevisiae requires a TID1/RDH54- and DMC1-dependent pathway. Genetics 163, 1273–1286.

Shinohara, M., Oh, S.D., Hunter, N. and Shinohara, A. (2008). Crossover assurance and crossover interference are distinctly regulated by the ZMM proteins during yeast meiosis. Nat. Genet. 40, 299–309.

Snowden, T., Acharya, S., Butz, C., Berardini, M. and Fishel, R. (2004). hMSH4-hMSH5 recognizes holliday junctions and forms a meiosis-specific sliding clamp that embraces homologous chromosomes. Mol. Cell 15, 437–451.

Song, B., Hamdan, S.M., and Hingorani, M.M. (2018). Positioning the 5’-flap junction in the active site controls the rate of flap endonuclease 1–catalyzed DNA cleavage. J. Biol. Chem. 293, 4792–4804.

Sugawara, N., Paques, F., Colaiacovo, M., and Haber, J. E. (1997). Role of *Saccharomyces cerevisiae* Msh2 and Msh3 repair proteins in double-strand break-induced recombination. Proc. Natl. Acad. Sci. USA 94, 9214–9219.

Sun, X., Thrower, D., Qiu, J., Wu, P., Zheng, L., Zhou, M., Bachant, J., Wilson, D. M., and Shen, B. (2003). Complementary functions of the Saccharomyces cerevisiae Rad2 family nucleases in Okazaki fragment maturation, mutation avoidance, and chromosome stability. DNA Repair 2, 925–940.

Sym, M., and Roeder, G.S. (1994). Crossover interference is abolished in the absence of a synaptonemal complex protein. Cell 79, 283–292.

Sym, M., Engebrecht, J., and Roeder, G.S. (1993). ZIP1 is a synaptonemal complex protein required for meiotic chromosome synapsis. Cell 72, 365–378.

Szankasi, P., and Smith, G.R. (1992). A single-stranded DNA exonuclease from *Schizosaccharomyces pombe*. Biochemistry 31, 6769–6773.

Szczelkun, M.D. (2002). Kinetic models of translocation, head-on collision, and DNA cleavage by type I restriction endonucleases. Biochemistry 41, 2067–2074.

Szostak, J.W., Orr-Weaver, T.L., Rothstein, R.J., and Stahl, F.W. (1983). The double-strand-break repair model for recombination. Cell 33, 25–35.

Thacker, D., Lam, I., Knop, M. and Keeney, S. (2011). Exploiting spore-autonomous fluorescent protein expression to quantify meiotic chromosome behaviors in *Saccharomyces cerevisiae*. Genetics 189, 423–439.

Tishkoff, D.X., Boerger A.L., Bertrand P., Filosi N., Gaida G.M., Kane M.F., Kolodner R.D. (1997). Identification and characterization of *Saccharomyces cerevisiae EXO1*, a gene encoding an exonuclease that interacts with *MSH2*. Proc. Natl. Acad. Sci. USA 94, 7487–7492.

Tomlinson, C.G., Atack, J.M., Chapados, B., Tainer, J.A., and Grasby, J.A. (2010). Substrate recognition and catalysis by flap endonucleases and related enzymes. Biochem. Soc. Trans. 38, 433–437.

Tran, P.T., Erdeniz, N., Dudley, S. and Liskay, R.M. (2002). Characterization of nuclease-dependent functions of Exo1p in *Saccharomyces cerevisiae*. DNA Repair 1, 895–912.

Tran, P.T., Erdeniz, N., Symington, L.S., and Liskay, R.M. (2004). EXO1-A multi-tasking eukaryotic nuclease. DNA Repair 3, 1549–1559.

Tran, P.T., Fey, J.P., Erdeniz, N., Gellon, L., Boiteux, S., and Liskay, R.M. (2007). A mutation in EXO1 defines separable roles in DNA mismatch repair and post-replication repair. DNA Repair 6, 1572–1583.

Tran, P.T., Simon, J.A., and Liskay, R.M. (2001). Interaction of EXO1 with components of MutLα in *Saccharomyces cerevisiae*. Proc. Natl. Acad. Sci. USA 98, 9760–9765. doi: 10.1073/pnas.161175998.

Tsubouchi, H., and Ogawa, H. (2000). Exo1 roles for repair of DNA double-strand breaks and meiotic crossing over in *Saccharomyces cerevisiae*. Mol. Biol. Cell 11, 2221–2233. doi: 10.1091/mbc.11.7.2221.

Tsutakawa, S.E., Thompson, M.J., Arvai, A.S., Neil, A.J., Shaw, S.J., Algasaier, S.I., Kim, J.C., Finger, L.D., Jardine, E., Gotham, V.J.B., Sarker, A.H., Her, M.Z., Rashid, F., Hamdan, S.M., Mirkin, S.M., Grasby, J.A., Tainer, J.A. (2017). Phosphate steering by Flap Endonuclease 1 promotes 5’-flap specificity and incision to prevent genome instability. Nat. Commun. 8, 15855. doi: 10.1038/ncomms15855.

Wild, P., Susperregui, A., Piazza, I., Dorig, C., Oke, A., Arter, M., Yamaguchi, M., Hilditch, A.T., Vuina, K., Chan, K.C., Gromova, T., Haber, J.E., Fung, J.C., Picotti, P., and Matos, J. (2019). Network Rewiring of Homologous Recombination enzymes during mitotic proliferation and meiosis. Mol. Cell 22, 859–874.e4. doi:10.1016/j.molcel.2019.06.022.

Xie, Y., Liu, Y., Argueso, J.L., Henricksen, L.A., Kao, H.I., Bambara, R.A., and Alani, E. (2001). Identification of *rad27* mutations that confer differential defects in mutation avoidance, repeat tract instability, and flap cleavage. Mol. Cell. Biol. 21, 4889–4899.

Zakharyevich, K., Ma, Y., Tang, S., Hwang, P.Y.-H., Boiteux, S., and Hunter, N. (2010). Temporally and biochemically distinct activities of Exo1 during meiosis: double-strand break resection and resolution of double Holliday junctions. Mol. Cell 40, 1001–1015.

Zakharyevich, K., Tang, S., Ma, Y., and Hunter, N. (2012). Delineation of joint molecule resolution pathways in meiosis identifies a crossover-specific resolvase. Cell 149, 334–347.

Zhang, L., Wang, S., Yin, S., Hong, S., Kim, K.P., and Kleckner, N. (2014). Topoisomerase II mediates meiotic crossover interference. Nature 511, 551–556.

Zickler, D., and Kleckner, N. (2015). Recombination, pairing, and synapsis of homologs during meiosis. Cold Spring Harb. Perspect. Biol. 7, a016626.

